# A mixture of innate cryoprotectants is key for freeze tolerance and cryopreservation of a drosophilid fly larva

**DOI:** 10.1101/2021.12.15.472769

**Authors:** Lukáš Kučera, Martin Moos, Tomáš Štětina, Jaroslava Korbelová, Petr Vodrážka, Lauren Des Marteaux, Robert Grgac, Petr Hůla, Jan Rozsypal, Miloš Faltus, Petr Šimek, Radislav Sedlacek, Vladimír Koštál

## Abstract

Insects that naturally tolerate internal freezing produce complex mixtures of multiple cryoprotectants (CPs). Better knowledge on composition of these mixtures, and on mechanisms of how the individual CPs interact, could inspire development of laboratory CP formulations optimized for cryopreservation of cells and other biological material. Here we identify and quantify (using high resolution mass spectrometry) a range of putative CPs in larval tissues of a subarctic fly, *Chymomyza costata* that survives long-term cryopreservation in liquid nitrogen. The CPs (proline, trehalose, glutamine, asparagine, glycine betaine, glycerophosphoethanolamine, glycerophosphocholine, and sarcosine) accumulate in hemolymph in a ratio of 313:108:55:26:6:4:3:0.5 mmol.L^-1^. Using calorimetry, we show that the artificial mixtures, mimicking the concentrations of major CPs’ in hemolymph of freeze-tolerant larvae, suppress the melting point of water and significantly reduce the ice fraction. We demonstrate in a bioassay that mixtures of CPs administered through the diet act synergistically rather than additively to enable cryopreservation of otherwise freeze-sensitive larvae. Using MALDI-MSI, we show that during slow extracellular freezing trehalose becomes concentrated in partially dehydrated hemolymph where it stimulates transition to the amorphous glass phase. In contrast, proline moves to the boundary between extracellular ice and dehydrated hemolymph and tissues where it likely forms a layer of dense viscoelastic liquid. We propose that amorphous glass and viscoelastic liquids may protect macromolecules and cells from thermomechanical shocks associated with freezing and transfer into and out of liquid nitrogen.

**Summary statement:** The composition of natural cryoprotectant mixture of the extremely freeze-tolerant insect is revealed. Components of the mixture work in synergy and behave differently during organismal freezing and cryopreservation.

## Introduction

Various organisms from bacteria to animals respond to environmental stressors such as heat, cold, freezing, drought, hypersalinity, or high hydrostatic pressure by accumulating a stereotypic set of cytoprotective compounds including amino acids, sugars, polyols, and methylamines (Somero, 1986; Yancey, 2005). This is also true for insects that naturally tolerate extracellular freezing during overwintering in temperate and polar habitats (Lee, 2010; Storey and Storey, 1988). In such insects, the accumulation of low molecular weight carbohydrates and free amino acids (cryoprotectants; CPs) is considered a fundamental tenet of their freeze tolerance (Storey, 1997; Storey and Storey, 1988; Storey and Storey, 1991; Toxopeus and Sinclair, 2018). Extensive knowledge has been accumulated over many decades about *which* CPs are present in many different species (Asahina, 1970; Salt, 1961; Sømme, 1982; Storey and Storey, 1991), but our understanding of *how* they protect the insect tissues and cells is far from complete (Toxopeus and Sinclair, 2018).

Here we focus on an important but previously neglected aspect of natural freeze tolerance: the components of complex *mixtures* of multiple CPs may contribute to freeze tolerance via additive or even synergistic mechanisms (Storey and Storey, 1986; Storey and Storey, 1988; Toxopeus et al., 2019b). It is well known from practice in clinical medicine and biotechnology that combinations of CPs protect the viability of cryopreserved cells or materials better than single substances (Elliott et al., 2017). The mixtures may even have emergent properties, i.e. those beyond the summation of properties of the individual components. Relatively recently, the existence of specific mixtures, so-called NADES (Natural Deep Eutectic Systems) in various organisms was proposed (Dai et al., 2013; Choi et al., 2011). These mixtures consist of stereotypic sets of natural primary metabolites such as sugars, sugar alcohols, organic acids, amino acids, and amines, and are characterized by a much lower melting point than that of the individual components, mainly due to the formation of intermolecular hydrogen bonds (Dai et al., 2015). These mixtures have been proposed to serve different physiological functions, including survival of organisms in extreme drought and cold (Castro et al., 2018; Gertrudes et al., 2017; Liu et al., 2018) and have also been proposed as new agents for cell cryopreservation (Hornberger et al., 2021). We believe that there may be major practical consequences of exploring the complex composition of insect CP mixtures, as they could aid in the development of applicable cryogenic techniques. Freeze-tolerant insects also offer the potential for the discovery of novel components, for the study of interactions between individual components, and for the analysis of the principles underlying survival after freezing in a complex organism composed of different cell types and tissues; a complexity that still poses a challenge for medical cryopreservation (Fahy et al., 2006; Pegg, 2001).

It is reasonable to first analyze the suite of CPs accumulated by the most freeze-tolerant model species, and the larvae of the malt fly, *Chymomyza costata* (Diptera: Drosophilidae) are among the most cold-hardy animals known (Des Marteaux et al., 2019). Overwintering (diapausing), cold-acclimated larvae can survive freezing of all osmotically active water (68% of total body water), have no apparent lower thermal limit while frozen, and even survive long-term cryopreservation in liquid nitrogen (LN_2_) (Moon et al., 1996; Rozsypal et al., 2018; Shimada and Riihimaa, 1988). We have previously shown that high concentrations of the free amino acid proline acquired by diapausing *C. costata* larvae during cold acclimation are essential for survival of freezing and cryopreservation stress (Koštál et al., 2011b; Rozsypal et al., 2018). Here we extend our previous results to include whole complex of CPs accumulated by these freeze-tolerant larvae. Using a combination of four chromatographic mass spectrometric (MS) analytical platforms, we established metabolite profiles of four different *C. costata* tissues in two contrasting larval phenotypes that differ greatly in freeze tolerance and cryopreservability in LN_2_. By combining the results of the analytical chromatographic MS analysis with matrix-assisted laser desorption/ionization (MALDI) mass spectrometry imaging (MSI), we show that freeze-tolerant larvae accumulate putative CPs in all tissues, but especially in hemolymph. Using the same technique, we observed changes in CP localization during slow extracellular freezing associated with partial dehydration of hemolymph and tissues. We then used differential scanning calorimetry (DSC) to describe the thermal phase transition properties of artificial aqueous CP mixtures mimicking the concentrations of the five most abundant metabolites in the hemolymph. Finally, we show in a bioassay that mixtures of CPs administered through the diet work in a synergistic rather than an additive way to induce strong freeze tolerance and cryopreservability in otherwise freeze-sensitive larvae.

## Materials and methods

### Fly rearing and acclimations

A colony of *C. costata*, Sapporo strain (Riihimaa and Kimura, 1988), was reared on artificial diet in MIR 154 incubators (Sanyo Electric, Osaka, Japan) as described previously (Kostal et al., 1998; Lakovaara, 1969). Two phenotypic variants (LD and SDA) of the 3^rd^ larval instar were generated according to our earlier acclimation protocols (Des Marteaux et al., 2019; Koštál et al., 2011b; Rozsypal et al., 2018) (Fig. A1a). The LD larvae (warm acclimated, active larvae reared at long-long day conditions) have limited survival after freezing (35% survive to -5°C, 10% to -10°C, and none survive freezing to -20°C or below). In contrast, practically all SDA larvae (dipause larvae reared at short-day conditions and gradually acclimated to cold) survive deep freezing to -30°C or even -70°C, and 42.5% survive at least six months of cryopreservation in LN_2_ (Rozsypal et al., 2018). In addition, SDA larvae were frozen (variant SDA-frozen) according to a previously developed optimal freezing protocol (Rozsypal et al., 2018) schematically presented in Fig. A1b. Survival after freezing and cryopreservation in LN_2_ was assessed as the ability to pupate and metamorphose into the adult stage within 42 d after thawing (at a constant 18°C/LD).

To collect the hemolymph, 30 larvae were gently pierced and torn on a piece of Parafilm, creating a large droplet of pooled hemolymph. This droplet was then extracted using a calibrated glass capillary (Drummond Sci., Broomall, PA, USA). To obtain fat body, muscle, and midgut tissues, the larvae were quickly dissected under binocular microscope in phosphate-buffered saline. Dissection of a single larva took less than 2 min, and we took care to sample a similar part of the respective tissue from each individual. This allowed us to express each metabolite concentration as a ‘pool per tissue’. Only in hemolymph samples, where we could measure the exact sample volume, we could also calculate the concentration in mmol.L^-1^.

### Extraction of metabolites and analytical MS platforms

Analyses were performed for whole *C. costata* larvae (pools of five larvae taken in four replicates) or dissected tissues (pools of 30 tissues in four replicates). Whole larvae were weighed to obtain fresh mass, immersed in LN_2_, and stored at −80°C until analysis. Dissected tissues were collected in ice-cold extraction buffer (see below) and stored at −80°C until analysis. Frozen samples of whole larvae/tissues were melted on ice and homogenized in 400 μL of extraction buffer (methanol:acetonitrile:deionized water in a volume ratio of 2:2:1). The methanol and acetonitrile (Optima LC/MS) were purchased from Fisher Scientific (Pardubice, Czech Republic) and the deionized water was prepared using Direct Q 3UV (Merck, Prague, Czech Republic). Internal standards, *p*-fluoro-DL-phenylalanine, methyl α-D-glucopyranoside (both from Sigma-Aldrich, Saint Luis, MI, USA) were added to the extraction buffer, both at a final concentration of 200 nmol.mL^−1^. Samples were homogenized using a TissueLyser LT (Qiagen, Hilden, Germany) set to 50 Hz for 5 min (with a rotor pre-chilled to −20°C). Homogenization and centrifugation (at 20,000 g for 5 min at 4°C) were repeated twice and the two supernatants were combined.

We performed targeted analyses of 49 select metabolites (Table A1) using a combination of four different mass spectrometry-based (MS) analytical platforms that were described previously: ECF-LC/MS and ECF-GC/MS (Štětina et al., 2018), SILYL-GC/MS (Škodová-Sveráková et al., 2020), and HILIC-LC/MS (Škodová-Sveráková et al., 2021). More details on instrumentation and chromatographic columns are in Table A1. All 49 metabolites were identified against relevant standards (Sigma-Aldrich) and subjected to quantitative analysis using a standard calibration curve method. The analytical results were validated by simultaneously running blank samples (no larvae in the sample), standard biological quality control samples (the periodic analysis of a standardized larva/tissue sample – the pool of all samples), and quality control mixtures of amino acids (AAS18, Sigma-Aldrich).

### Matrix-assisted laser desorption/ionization mass spectrometry imaging (MALDI-MSI)

The MALDI-MSI (Khalil et al., 2017; Tuthill II et al., 2020; Yang et al., 2020) was conducted in LD and SDA larvae and also on SDA-frozen larvae (Fig. A1). Larvae were rinsed in water, incubated for 3 min in 2% gelatin, and then submerged into 5 mL of 10% gelatin in a plastic mold (Tissue-Tek Cryomold, Sakura, Mdesa, Czech Republic) placed on an ice-cold stage. The molds with solid gelatin were then immersed in isopentane pre-cooled to -140°C in LN_2_ vapors for 3 min and transferred to -80°C for storage until further processing. Preparation of the SDA-frozen larvae was modified as shown in Fig. A2.

For cryo-sectioning, the frozen molds were placed into a cryostat chamber (Leica CM1950, Germany) pre-cooled to -20°C. Sections of 10 μm thickness were mounted on indium tin oxide coated glass slide (ITO, Bruker, Czech Republic), which was previously washed with hexane and then with 2-propanol. Sections of the LD and SDA larval variants were thaw-mounted on warm glass (standard procedure MALDI-TOF/MSI) (TOF, Time Of Flight), while sections of the SDA-frozen larval variant were mounted on a gold-coated ITO glass pre-cooled to -20°C in order to preserve the tissue organization as it formed during slow pre-freezing (extracellular ice formation and tissue freeze-dehydration). The sections were dried for 15 min in a desiccator at low air pressure (at room temperature for the LD and SDA variants, while at -20°C for the SDA-frozen variant), then sealed in plastic foil inside a plastic mailer, evacuated, and stored at -80°C until further processing. The ITO glasses with sections were transferred to room temperature, left to temper for 15 min, and dried in a desiccator for 15 min. Samples were then sprayed with a matrix solution of 9-aminoacridine hydrochloride, 7 mg.mL^-1^ in 70% ethanol (v/v). A TM-Sprayer 3 (HTX-Technologies, USA) was used to cover the tissue with the matrix solution using the following settings: Nozzle temperature 50°C, 12 passes, flow 0.035 mL.min^-1^, nozzle velocity 1000 mm.min^-1^, track spacing 2 mm, HH pattern, gas pressure 10 psi, gas flow rate 2 L.min^-1^, drying time 0 sec, nozzle height 40 mm and propelled with 50% methanol (v/v). After spraying, the samples were dried in a desiccator for 15 min.

Spectral images were acquired using a rapifleX MALDI-TOF/TOF spectrometer (Bruker, Germany) in reflector negative mode with a 355 nm smartbeam™ 3D laser with a spatial resolution of 10×10 μm (pixel) in the range of m/z 20-1000, at a constant laser fluence of 72% and laser frequency of 5 kHz. Two hundred images were accumulated from every position. The instruments were set up with Ion source 1 to 19.973 kV, PIE 2.664 kV, Lens 11.353 kV, Reflector 1 20.810 kV, Reflector 2 1.034 kV, and Reflector 3 8.577 kV. The pulsed Ion Extraction time was set to 90 ns and detector gain was 2302 V. Data were acquired with digitizer speed 2.5 GS/s. Samples were measured in random order. Calibration was done externally using red phosphorus, achieving precision up to 5 ppm (Kolářová et al., 2016).

After MALDI-MSI, tissues on ITO glass were washed three times for 2 min with 70% ethanol and afterwards washed with water. The samples were then stained with hematoxylin & eosin (H&E) and scanned with Axio Scan.Z1 (Zeiss, Germany) operated by Zen 2 software (blue edition; Zeiss, Germany). Single raw data files on the intensities of the MALDI-TOF signals of individual m/z peaks were processed using SCiLS Lab software (version 2021c, SCiLS, Bruker, Germany). The raw data were smoothed with a convolution algorithm (width 20) and all subsequent computations and images rendering were done with Total Ion Count square normalization. We collected MALDI-TOF signal-intensity datasets for 5 transversal sections of LD and SDA larvae and for 6 sections of SDA-frozen larvae, plus one longitudinal section of an SDA larva. For relative quantification of signal intensities and statistical analyses, the transversal sections were used. Galleries showing the complete results for all transversal sections are included in separate Appendice.

Regions (or tissues) of interest in the sections were manually delimited using H&E stained images in QuPath software (version 0.2.3, University of Edinburgh, United Kingdom) (Bankhead et al., 2017) and imported back into SCiLS software. The summary spectrum was exported into mMass software (version 5.5.0) (Niedermeyer and Strohalm, 2012), internally re-calibrated, and annotated based on m/z values with 50 ppm tolerance using an in-house modified database based on the HMDB metabolite database considering [M-H]^-^ and [M+Cl]^-^ adducts (Wishart et al., 2018). Selected peaks were then analyzed using SCiLS software with mean spectral intensity scaled in arbitrary units (A.U.) for each region and treatment. These spectral intensity data were then statistically compared using ANOVA followed by Bonferroni’s multiple comparison test (Prism, v. 6.07, GraphPad Software, San Diego, USA). In addition, we verified the identities of m/z peaks for trehalose and proline in the experiment where LD larvae were fed diet augmented by ^13^C6 glucose (for details, see Figs. A13 and A14). The ^13^C6 glucose was purchased from Sigma Aldrich (product ID 389374).

### Differential scanning calorimetry (DSC)

Thermal analysis was performed using the Q2000 calorimeter (TA Instruments, New Castle, DE, USA) as previously described (Pokorna et al., 2020). The analyzed solutions (see Table A2 for a complete list) were loaded and hermetically sealed into aluminum pans. The amount of loaded solution (ca. 5 μL) was controlled by weighing with precision to 0.01 mg using a Sartorius CP 225 D balance (Sartorius AG, Goettingen, Germany). The solutions were frozen and thawed according to the following protocol: (i) start at 25°C, (ii) cool to −90°C at 10°C.min^-1^, (iii) hold for 5 min at −90°C, and (iv) warm to 25°C at 10°C.min^-1^. An empty pan was used as a reference, and samples were run in technical triplicates (same solution analyzed thrice). DSC results were analyzed using TA Universal Analysis 2000 software (version 4.5A Build 4.5.0.5). The onset of exotherm on the cooling scan was taken as the supercooling point (temperature of ice crystallization). Two major thermal events were analyzed on heating scans above -60°C: (i) glass transition: an inflection point of the second-order phase transition was read as the temperature of vitrification (T_g_). The change in specific heat capacity (ΔCp) was derived from the difference in heat flow between the onset and the end of the glass transition; (ii) melting of bulk water: an onset of melting endotherm (first-order phase transition) was read as the melting point (m.p.), while the enthalpy of melting (ΔH, calculated from the area under the endothermic peak) served to estimate the fraction of melted (i.e. crystallized) water using the standard heat of fusion for the ice/water transition of 334 J g^-1^. The remaining fraction of unfrozen water was considered osmotically inactive (%OIW).

### Larval feeding on CP-augmented diets

Seventeen day-old LD larvae (i.e. roughly mid-3^rd^ instar, voracious feeding stage) were extracted from the standard diet and groups of 20 individuals were moved to one of a variety of specific diets: (i) fresh standard diet (control); (ii) ‘CP mix 508’ diet – a standard diet augmented with five select CPs (proline, trehalose, glutamine, asparagine, and betaine) in concentrations corresponding to those observed in SDA larval hemolymph (i.e. 508 mmol.kg^-1^ water in total, see Table A3); (iii) five different standard diets augmented with the select CPs individually; (iv) ‘CP mix minus Pro 313’ diet – a mixture of the four select CPs but without the 313 mmol.kg^-1^ proline. After three days exposure to these diets, the larvae were removed and used for the LN_2_ survival bioassay (Fig. A1b).

## Results

### Characterization of the CP mixture from *C. costata* larvae

Results of targeted quantitative analysis of 49 metabolites are summarized in Table A1 and shown diagrammatically in Fig. 1a (hemolymph) and Fig. A3 (other tissues). For most metabolites, the pool in the hemolymph was significantly larger than the pools in other tissues (Fig. A4). The volume of hemolymph in one larva is approximately 200 nL (see Table A1) representing only 10% of the total body volume or body mass (about 2 mg). Nevertheless, the hemolymph contained approximately 50% of the total metabolite pool in both LD and SDA larvae (Table A1). Entry into diapause and cold acclimation (associated with the acquisition of extreme freeze tolerance) induced accumulation of specific metabolites (putative CPs) in SDA larvae (Fig. A1). The sum of the molar concentrations of the 49 metabolites in LD hemolymph was 182 mmol.L^-1^, but this had increased to 554 mmol.L^-1^ in SDA hemolymph (Table A1). Four compounds that contributed most to this increase were: proline (increase of 277 mmol.L^-1^), trehalose (increase of 59 mmol.L^-^), glutamine (increase of 22 mmol.L^-1^), and asparagine (increase of 17 mmol.L^-1^). The same four compounds also ranked first in order of concentration in SDA hemolymph: proline (313 mmol.L^-1^), trehalose (108 mmol.L^-1^), glutamine (55 mmol.L^-1^), and asparagine (26 mmol.L^-1^). Glycine-betaine (hereafter ‘betaine’, 6.1 mmol.L^-1^) was fifth in concentration, although it increased only moderately (by 1.7 mmol.L^-1^) from LD to SDA. In addition, three other compounds had statistically higher hemolymph concentrations in SDA compared to LD larvae, although their concentrations in SDA hemolymph were relatively low in absolute terms: glycerophosphoethanolamine (GPE, 4.0 mmol.L^-1^), glycerophosphocholine (GPC, 2.9 mmol.L^-1^), and sarcosine (0.5 mmol.L^-1^). For the purposes of this article, we consider the above compounds as putative CPs.

**Figure 1:**
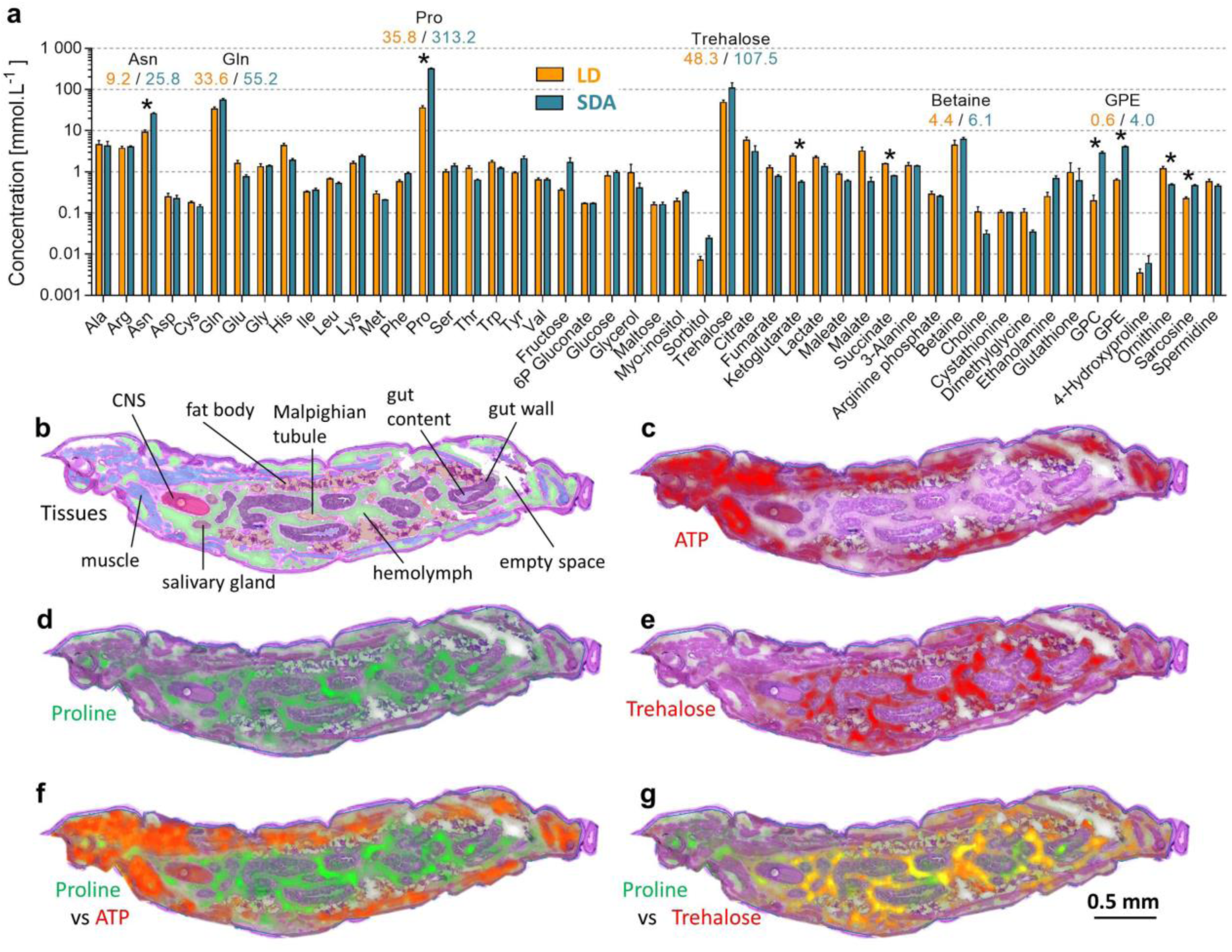
Metabolite profiles and localization in hemolymph and tissues of *C. costata* larvae. **(a)** Forty-nine target metabolites were absolutely quantified in the hemolymph of freeze sensitive (LD) and freeze tolerant (SDA) larvae using four MS-based analytical platforms. Each column represents a metabolite concentration (mean ± SD; *n* = 4 biological replicates, each containing hemolymph from 30 larvae). Differences between the means of LD and SDA were assessed by unpaired t-tests corrected for multiple comparisons using the Holm-Sidak method with α = 0.01. Asterisks indicate significantly different means. **(b)** Longitudinal section of a SDA larva stained with hematoxylin and eosin, with tissues highlighted by different colors. (c-e) Intensities of the MALDI-TOF signals corresponding to the three analyzed compounds: **(c)** ATP; **(d)** proline; **(e)** trehalose. **(f)** While proline (green signal) is most evident in hemolymph, ATP (red signal) is mainly found in muscles. **(g)** Proline (green) and trehalose (red) co-localize in hemolymph (a yellow color results from the mixing of green and red). See Fig. A3 for MALDI-MSI images of other compounds.

### Localization of CP mixture components in larval tissues

Using MALDI-MSI we localized select metabolites in larval sections. A longitudinal section through the SDA larva provides the best overview of tissue structure (Fig. 1b). The highest intensity of the MALDI-TOF signal for ATP is localized predominantly in muscle tissue (corresponding to its role as the most important intracellular phosphagen) (Fig. 1c). In contrast, the highest intensities of proline and trehalose signals are localized in the hemolymph (Fig. 1d, e) where they are also colocalized with the high-intensity signals of glutamine, asparagine, and GPE (Fig. A5).

As it was difficult to obtain a sufficient number of high quality longitudinal sections, we used transverse sections through the middle region of the larvae for the statistical analysis of the relative intensities of the MALDI-TOF signals. Quantitative analysis of these signals is technically difficult for small metabolite molecules freely dissolved in biological aqueous solutions (Fig. A6). Nevertheless, our analysis suggests that there are no differences in the overall localization patterns of the five putative CPs (proline, trehalose, glutamine, asparagine, and GPE) between LD and SDA larvae. However, the signal intensities for all five CPs were much stronger in SDA than in LD larvae (compare the *y*-axes of Figs. A7 vs A8). The MALDI-MSI thus confirms the results of the quantitative MS analysis (Fig. 1a) that the putative CPs are accumulated in all tissues, but especially in the hemolymph, of SDA larvae (for details, see Appendice containing all MALDI-MSI images).

During slow inoculative freezing to -30°C (rate of 0.1°C.min^-1^ ; see Fig. A1b for the freezing protocol) large masses of extracellular ice crystals develop in between partially dehydrated pools of larval hemolymph and tissues (Fig. 2a-d). We observed a striking difference in the ‘behavior’ of two major components of the CP mixture during slow extracellular freezing: while trehalose was concentrated in the pool of dehydrated hemolymph of SDA larvae, a large quantity of proline apparently ‘migrated’ from the hemolymph toward the ice region (Fig. 2e, f) and formed thin ‘layers’ of concentrated proline at the boundary between the extracellular ice and the dehydrated hemolymph and tissues (Fig. 2g-i). In these layers, proline is colocalized with high-intensity signals from glutamine and asparagine, while trehalose is colocalized with GPE in the dehydrated hemolymph (Fig. A9; for details, see Appendice containing all MALDI-MSI images).

**Figure 2:**
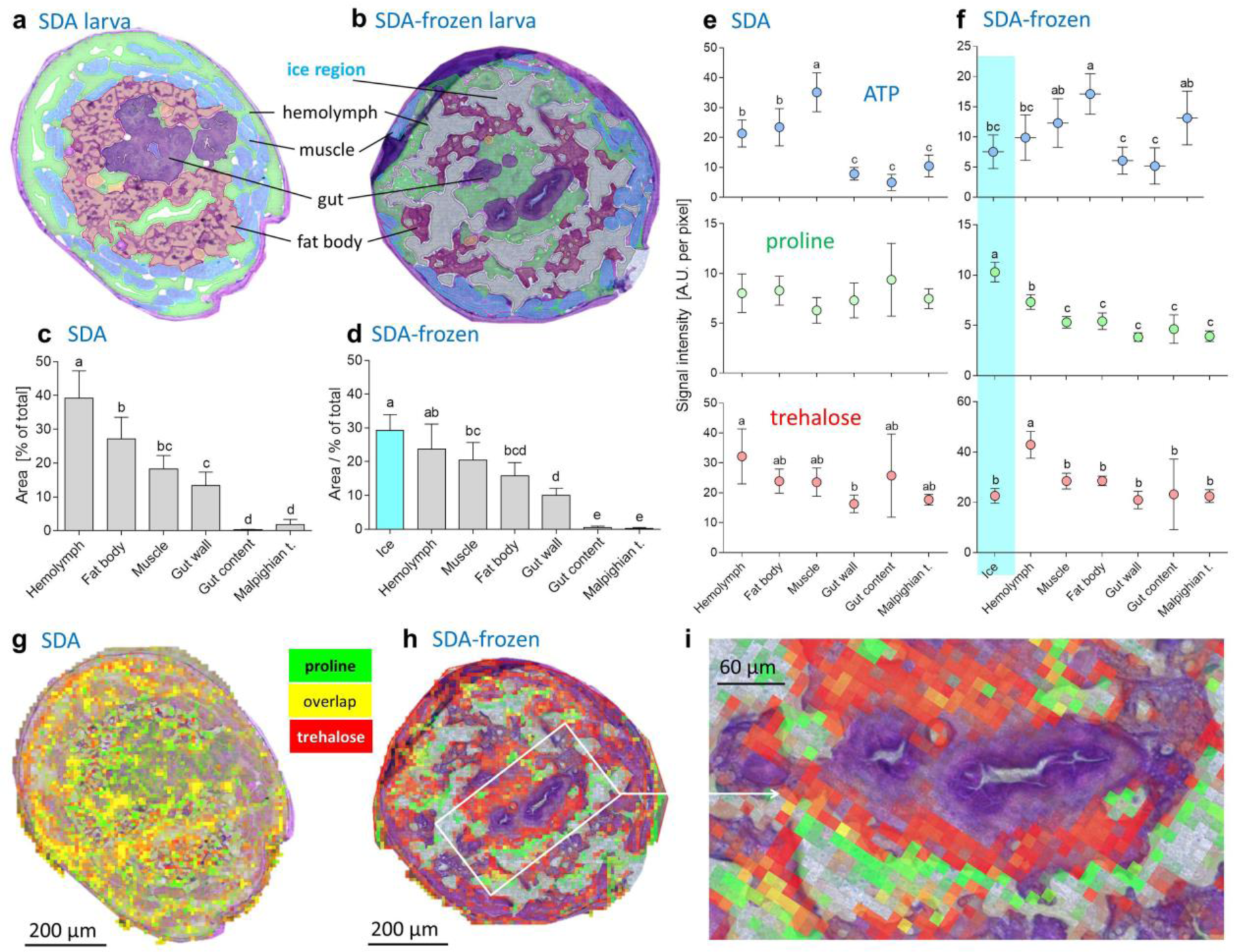
Localization of proline and trehalose *in C. costata* SDA larvae prior to and after slow freezing. **(a, b)** Examples of transverse sections through the middle region of SDA-larva (a) and SDA-frozen-larva (b). The tissues are highlighted by different colors. Note the large masses of extracellular ice between the partially dehydrated tissues of SDA-frozen-larva. **(c, d)** Relative proportions of total area occupied by different tissues or ice. Each column is a mean ± S.D. of five or six sections (SDA or SDA-frozen larvae, respectively). Means flanked by different letters are significantly different (one way ANOVA followed by Bonferroni’s multiple comparison test). **(e, f)** Results of MALDI-TOF signal intensity quantification (arbitrary units, A.U. per pixel of 10×10 um) for three select metabolites (ATP, proline, trehalose). Each point is a mean ± S.D. of five or six sections (SDA or SDA-frozen larvae, respectively). Mean values flanked by different letters are significantly different (one way ANOVA followed by Bonferroni’s multiple comparison test). **(g, h, i)** Examples of MALDI-TOF signal intensities for proline (green) and trehalose (red). Note that proline and trehalose signals markedly overlap (yellow color) in the hemolymph of SDA larva (g), while they apparently de-localize in SDA-frozen-larva (h, i). Trehalose concentrates in the partially dehydrated hemolymph, while proline migrates to the border between extracellular ice and partially dehydrated tissues of the SDA-frozen larva.

### Thermal phase transitions in artificial CP mixtures

To assess the influence of different CP mixture components on water binding and thermal phase transitions, we compared heating curves from DSC scans of aqueous solutions containing single components or different CP mixtures with real SDA hemolymph. We focused on melting of bulk water (Fig. 3a) and glass transition (Fig. 3b). The full DSC analysis results can be found in Table A2.

**Figure 3:**
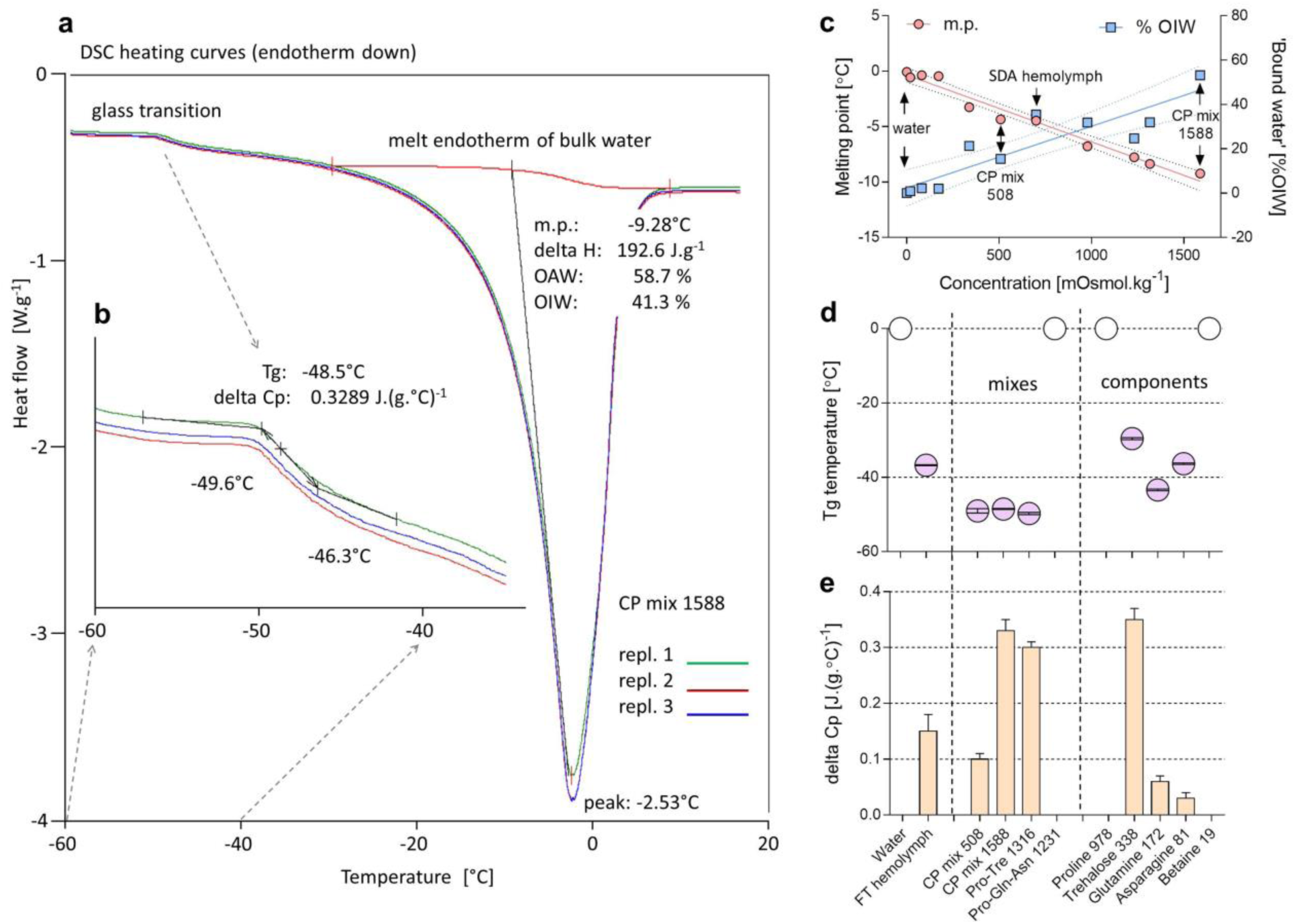
Thermal phase transitions in the aqueous solutions of CP mixtures and individual components. **(a)** Example of a heating curves recorded by differential scanning calorimetry (DSC) of the artificial CP mix 1588 (for explanation, see text). Three technical replicates are shown in different colors. Results calculated by TA Universal Analysis 2000 software are shown for the 1st replicate only (green line). The analysis of melting endotherm shows that melting of bulk water starts at an m.p. of -9.28°C, while only 58.7% of the total water in the sample is osmotically active (OAW), i.e. melts (the enthalpy of melting transition of ΔH of 334 J.g^-1^ would correspond to melting of 100% total water) while 41.3% of total water is osmotically inactive (OIW) or ‘bound’. **(b)** A glass transition was observed at -48.5°C (T_g_, inflection point) associated with change in specific thermal capacity ΔCp of 0.3289 J.(g.°C)^-1^. **(c)** Linear associations of the melting point (m.p., red line) or the fraction of bound water (%OIW, blue line) with the osmolality of the solution. Each point represents a specific solution and all solutions are shown in detail in Table A2. Points corresponding to water, SDA hemolymph, and CP mixes 508 and 1588 are identified for clarity. The broken lines show 95% confidence intervals of the linear regressions. **(d, e)** Temperatures of the glass transition, Tg (d) and changes in thermal capacity, ΔCp (e) of the different solutions. Empty points in (d) or a missing column in (e) indicates that no glass transition was observed, while the violet points and orange columns show mean ± S.D. values of T_g_ or ΔCp, respectively, for solutions where a glass transition was observed (see Table A2 for the full dataset).

We designed the CP mixtures according to the concentrations of the five most abundant (and commercially available / affordable) compounds in the SDA larval hemolymph. CP mix 508 mimics the composition of the major CPs in the hemolymph of SDA larvae prior to freezing, while CP mix 1588 takes into account a 3.125-fold increase in the concentration of all components resulting from extracellular freezing of osmotically active water (68%). In addition, we tested all components individually at concentrations equivalent to those estimated in SDA hemolymph after extracellular freezing, and we also designed two reduced-composition mixes: Pro-Tre 1316 (proline 978 + trehalose 338) and Pro-Gln-Asn 1231 (proline 978 + glutamine 172 + asparagine 81) (see Table A2 for details).

We found that depression of the bulk water melting point, as well as the fraction of ‘bound’ water (unfrozen, osmotically inactive water, %OIW), correlate linearly with the molal concentration of a solution, regardless of whether it is a single component solution or a mixture. Even the values for real hemolymph collected from SDA larvae fit well to the linear relationships (Fig. 3c).

Whenever trehalose was present in a mixture, a clear glass transition was observed (Fig. A10). The temperatures of glass transition (T_g_) in different trehalose-containing solutions varied from -29.6°C (trehalose 338) to -49.7°C (Pro-Tre 1316) (Fig. 3d). The change of thermal capacity (ΔCp) during de-glassing was slightly above 0.3 J.(g.°C)^-1^ in the solutions containing a relatively high concentration of trehalose (338 mmol.kg^-1^) but lower, ∼ 0.1 J.(g.°C)^-1^, in the solution containing a relatively low concentration of trehalose (CP mix 508 contains 108 mmol.kg^-1^ of trehalose) and in real SDA hemolymph (which also contains 108 mmol.kg^-1^ of trehalose) (Fig. 3e). No glass transition was observed in the concentrated proline 978 solution (Fig. A11). The presence of proline in a mixture with glutamine and asparagine (Pro-Gln-Asn 1231) resulted in the elimination of small glass transitions observed in single-component mixtures of glutamine or asparagine (Fig. 3d, e). We also observed minor exotherms in heating curves of glutamine and asparagine at -22.8°C and -14.6°C, respectively (Fig. A12). The presence of such exotherms was also eliminated by the addition of proline in the Pro-Gln-Asn 1231 mix (Table A2).

### CPs in mixture protect larvae from cryopreservation stress better than individual CPs

To evaluate the ability of a complex cryoprotective mixture to increase freeze tolerance compared to individual components of the CP mix, we fed LD larvae with CP-augmented diets and then performed freezing survival assays. We prepared the CP mix 508 diet by augmenting the standard larval diet with a CP mix 508 solution (see Table A3). When LD larvae were fed with the CP mix 508 diet from hatching, more than 90% died (often as 1^st^ instars). We therefore exposed only 3^rd^ instar LD larvae to this diet and for only three days (at ages 17-19 days), which reduced larval mortality to 20%. The surviving larvae were significantly smaller at day 19 and pupariation was delayed by two days (Table A4), indicating that even short-term exposure to the CP mix 508 appears to have some toxic effects.

A three-day exposure to the CP mix 508 diet significantly increased whole-body concentrations of most CP mix components: proline by 6.8-fold, trehalose by 1.7-fold, glutamine by 5.4-fold, and asparagine by 19.9-fold (see Table A5 for more details). In the hemolymph, the concentrations of proline, glutamine, and asparagine reached high levels of 289, 80, and 49 mmol.L^-1^, respectively, while trehalose and betaine remained relatively low at 29 and 7 mmol.L^-1^, respectively (Fig. 4a, see Table A5 for more details). The sum molarity of the five CPs in hemolymph reached 454 mmol.L^-1^ in LD larvae fed with the CP mix 508 diet, which is broadly similar to the sum of 508 mmol.L^-1^ seen in SDA larvae.

**Figure 4:**
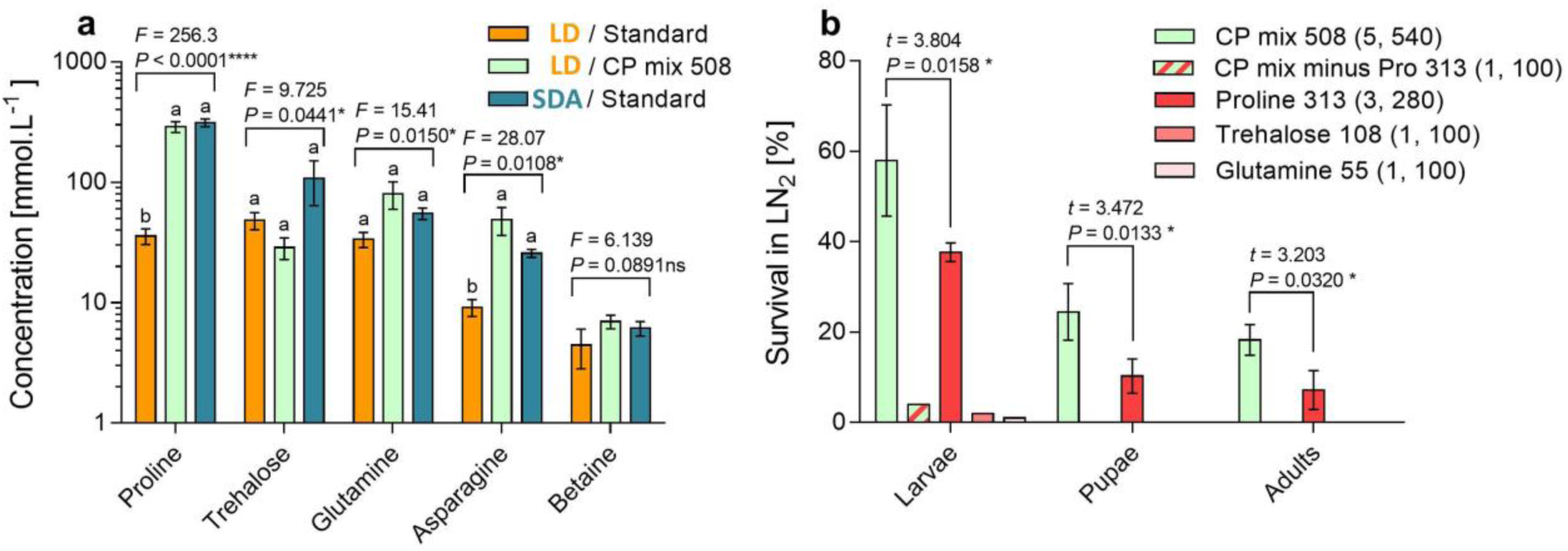
Effects of CP-augmented diets on *C. costata* larvae. **(a)** Change in the hemolymph concentration of five CPs in response to feeding 17 day-old LD larvae with a CP mix 508 diet for three days. Concentrations are compared with LD and SDA larvae fed on a standard diet, using one way ANOVAs followed by Bonferroni’s multiple comparison tests (columns flanked by different letters differ significantly). **(b)** Percent survival of larvae cryopreserved in LN_2_ (note: only diets where at least one larva survived are shown in panel b). The 17 day-old LD larvae were fed different diets for 3 days, then exposed to LN_2_ for 1 h, returned to the standard diet at a constant 18°C, and their survival (movements 24 h later), pupation, and adult emergence were observed over the next 14 days. The two numbers in parentheses (*n, N*) show: *n* = the number of fly generations, and *N* = the total number of larvae in all replicates. The mean survival for larvae fed a CP mix 508 diet versus a proline diet (Pro 313) were directly compared using a two-tailed Student’s *t*-test.

No LD larvae that fed exclusively on the standard diet survived cryopreservation in LN_2_. In striking contrast, 58% of the LD larvae survived cryopreservation after only three days of feeding with the CP mix 508 diet; 24.5% went on to pupate and 18.3% successfully metamorphosed to the adult stage (Fig. 4b). The components of the CP mix 508 administered via the diet had much less (or no) effect on larval freeze tolerance when they acted in isolation: betaine and asparagine had absolutely no effect; glutamine and trehalose resulted in 1 and 2 larvae surviving (out of 100), respectively, but these larvae did not pupate; proline augmentation was most effective, resulting in 37.7%, 10.3% and 7.2% survival for larvae, pupae, and adults, respectively. When proline was excluded from the CP mix, the remaining components allowed only 4 larvae (out of 100) to survive LN_2_ exposure, demonstrating that proline is the most important component. Nevertheless, the complete CP mix 508 was approximately 2-fold more effective at bolstering LN_2_ survival than proline alone. This result was consistent at all three levels of survival (see *t*-tests in Fig. 4b) and also across five different generations of assayed insects (Table A6). Simple addition of the individual effects of the three components (proline + trehalose + glutamine) was not sufficient to explain the effect of the complete CP mix 508. These results suggest a synergy between CP mix 508 components that should be investigated in the future.

## Discussion

### Components of the fly’s innate CP mixture behave differently during slow extracellular freezing

We identified the composition of the innate putative CP mixture which bathes the tissues of extremely freeze-tolerant *C. costata* larvae: proline, trehalose, glutamine, asparagine, betaine, GPE, GPC, and sarcosine, which occur in hemolymph in a stoichiometric ratio of 313:108:55:26:6:4:3:0.5 mmol.L^-1^, respectively. It may not be surprising that the mixture is broadly similar to the cocktails known from other organisms exposed to various environmental stressors (Gertrudes et al., 2017; Choi et al., 2011; Somero, 1986; Yancey, 2005). The occurrence of the same classes of organic cytoprotectants and compatible osmolytes in different organisms is likely based on convergent evolution which favors accumulation of polar molecules with high solubility in water (high accumulation capacity) countered by low tendency to interact with charged groups of macromolecules and membranes (low toxicity) (Hochachka and Somero, 2002).

We performed a bioassay showing that supplementation of the larval diet with major components of the CP mixture in their native stoichiometric ratio changes freeze sensitive-larvae to freeze-tolerant/cryopreservable-larvae within just three days. This result convincingly demonstrates that the accumulation of CPs has a strong functional (adaptive) significance in *C. costata*. However, this adaptation has probably evolved to cope with conditions typical for larvae overwintering in thermally buffered microhabitats under the bark of fallen trees, often under a blanket of snow (Band and Band, 1982; Grimaldi, 1986). Therefore, there may be different functional explanations for the roles of CPs in ecologically relevant situation in the field and for deep freezing and cryopreservation in LN_2_ in the laboratory. Here we avoid the eco-physiological aspects of CP-based freeze tolerance in animals and insects in the field, as these have been discussed for animals (Sinclair, 1999; Storey and Storey, 1988; Toxopeus and Sinclair, 2018; Zachariassen, 1985) and also specifically for *C. costata* (Rozsypal et al., 2018) elsewhere. Instead, we limit further discussion to the potential relevance of our results for deep freezing and cryopreservation in the laboratory.

We applied the MALDI-MSI technique to localize CPs in larval tissues and, for the first time in cryobiology literature, to determine how this localization changes during slow extracellular freezing, i.e. under conditions that allow larvae to survive deep freezing and subsequent cryopreservation in LN_2_. So far, concentrations of CPs in insects have mostly been reported in whole body samples for two general reasons: first, insect tissues are small, making it technically difficult to estimate precise concentrations; and second, native CPs – being standard products of metabolism – have been considered to move freely across membranes (Somero, 1986). In cases where CP analyses were performed on separate tissues (Koštál et al., 2011a; Rozsypal et al., 2013; Toxopeus et al., 2019a), the CPs were generally found in all examined tissues. Here we have confirmed that SDA larvae of *C. costata* accumulate putative CPs in all examined tissues but that the highest concentrations occur in the hemolymph.

One of the most salient findings of this study is the observation that the two major components of the CP mixture, proline and trehalose, ‘behave’ differently during slow extracellular freezing. The MALDI-MSI analysis suggests that the bulk of trehalose molecules remain in their original location (before freezing) and concentrate in partially freeze-dehydrated hemolymph and tissues. In contrast, some of proline molecules appear to move out of the hemolymph and concentrate in thin layers separating the extracellular ice crystals from freeze-dehydrated tissues and hemolymph. At this stage of research, we can only offer hypothetical/speculative explanations for the physico-chemical mechanisms responsible for de-localization of proline and trehalose and for its possible functional significance in a cryopreserved insect. We suggest that the mechanism behind the de-localization could be explained by differences between trehalose and proline in water solubility, propensity to induce glass transition, and mobility in highly viscous freeze-concentrated solutions.

### Trehalose stimulates glass transition in partially freeze-dehydrated hemolymph and tissues

We have previously shown that in gradually frozen SDA larvae a glass transition of the residual solution occurs between -20°C and -30°C. Moreover, only larvae pre-frozen to temperatures below -30°C (i.e. below the temperature of glass transition) were able to survive abrupt submersion and cryopreservation in LN_2_ and successfully develop into adults (Rozsypal et al., 2018). This result clearly demonstrated the importance of the glass transition for survival in LN_2_. In our earlier work (Rozsypal et al., 2018), we attributed glass formation to high concentrations of proline in accordance with results of the DSC analysis performed by (Rudolph and Crowe, 1986). However, this observation was challenged in later DSC studies (Liu et al., 2020; Rasmussen et al., 1997) and our own analysis confirms that proline at a concentration of 978 mmol.kg^-1^ does not induce glass transition, at least not at temperatures above -60°C. Instead, trehalose is known in physical chemistry as one of the strongest glass transition inducers in aqueous systems (Cesaro et al., 2008; Green and Angell, 1989; Chen et al., 2000). Nicolajsen and Hvidt (Nicolajsen and Hvidt, 1994) reported that a water:trehalose mixture (%weight, 89.7:10.3) de-glasses at -30.5°C with a ΔCp of 0.38 J.(g.°C)^-1^. These values agree well with our DSC analysis results showing de-glassing at -29.6°C with a ΔCp of 0.33 J.(g.°C)^-1^ for the trehalose 338 solution (which is 88.7:11.3 %weight of water:trehalose). Accurate interpretation of the thermal behavior and glass transition parameters is a challenge for physical chemistry of simple binary mixtures such as trehalose:water (Cesaro et al., 2008; Chen et al., 2000; Nicolajsen and Hvidt, 1994; Olgenblum et al., 2020; Weng and Elliott, 2014). This limitation becomes almost insurmountable for more complex mixtures and, especially, for biological solutions. Nevertheless, our DSC analysis suggests that trehalose rather than proline may be the component mainly responsible for the glass transition in larval body fluids: (i) all artificial CP mixtures containing trehalose showed clear glass transitions with broadly similar T_g_ and ΔCp parameters; (ii) the parameters were similar for real hemolymph collected from SDA larvae (containing 108 mmol.L^-1^ trehalose) and for the artificial CP mix 508 (simulating composition of major CPs in SDA hemolymph and also containing 108 mmol.kg^-1^ trehalose); (iii) no glass transition was observed in the concentrated proline 978 solution.

During glass transition, the viscosity of the supercooled liquid increases rapidly as the frequency, strength, and lifetime of hydrogen bonds in a system increases. In dilute systems, most hydrogen bonds are formed between water molecules, but as the concentration of solute molecules increases the interactions between water-trehalose and trehalose-trehalose become more frequent until trehalose molecules form large hydrogen-bonded clusters that exclude water (Olgenblum et al., 2020; Weng and Elliott, 2014). The increase in viscosity slows the molecular dynamics and macromolecules become trapped in an amorphous hydrogen-bonded matrix, which probably prevents undesirable transitions such as protein unfolding. In addition, trehalose can directly form hydrogen-bonds with macromolecules which probably further increases their stability in the amorphous glass (Crowe et al., 1998; Olgenblum et al., 2020). Trehalose molecules involved in glass formation are relative immobile. Moreover, large proportions of water molecules are ‘bound’, i.e. osmotically inactive in SDA larval hemolymph (35.5%) or in whole-body *C. costata* (42% (Rozsypal et al., 2018)), and therefore do not contribute much to trehalose solvation. As a result, trehalose can even locally reach the solubility limit (about 1.7 mol.kg^-1^ according to ALOGPS, http://www.vcclab.org/lab/alogps/), partly crystallize, and form a eutectic system with ice crystals. Both crystallization and glass formation would make trehalose molecules immobile, i.e. gradually concentrated in dehydrated tissues and hemolymph, which we observed.

### Proline-rich viscoelastic liquid forms at the boundary between extracellular ice and dehydrated tissues

In contrast to trehalose, proline does not induce a glass transition (Liu et al., 2020; Rasmussen et al., 1997) and its solubility in water is enormous (it may reach over 15 mol.kg^-1^ (Held et al., 2014; Qiu et al., 2019)). Nevertheless, physical chemistry shows that proline in high concentration also extensively hydrogen bonds with water and together they form a viscoelastic liquid (de Molina et al., 2017; McLain et al., 2007; Troitzsch et al., 2008). The formation of dense, viscoelastic liquids is a guiding principle underlying the theory of NADES systems (Choi et al., 2011). The components in a complex mixture forming NADES are connected to one another through a well-organized tridimensional system with optimum interactions via inter- and intramolecular hydrogen bonding (Dai et al., 2013; Dai et al., 2015). Through these interactions, NADES systems exhibit emergent properties: (i) the melting point is deeply suppressed (usually below - 30°C); (ii) ice crystallization is completely (or at least strongly) inhibited; (iii) while the tendency to transition to amorphous glasses is enhanced (Castro et al., 2018; Liu et al., 2018; Martins et al., 2019). However, work with NADES has also shown that the interactions between the NADES components are weakened with water dilution and even broken with water content exceeding 50 % weight, when each component recovers its own specific properties (Castro et al., 2018; Dai et al., 2015; Durand et al., 2016). Our artificial CP mixtures have relatively high water content (between 79 and 99 %weight), as do the SDA larval hemolymph (74.4 %weight) and whole larvae (69.3 %weight). Using DSC analysis, we show that these solutions exhibit significant suppression of the melting point (e.g. to -9.2°C in CP mix 1588, and to -4.5°C in SDA hemolymph) and significant water-binding capacity (41.1 %OIW in CP mix 1588, and 35.5 %OIW in SDA hemolymph). Both parameters (the decrease in melting point and the increase in OIW fraction) were linearly dependent on the osmolality of a solution (from 0 for water to 1,588 mmol.kg^-1^ for CP mix 1588). These results suggest that neither the CP mixes 508 and 1588 nor the SDA hemolymph are ‘deep eutectic systems’ (DES, in the strict sense of (Martins et al., 2019)), but regular aqueous solutions with relatively low transition temperature from liquid to solid phase (LTTM, in the sense of (Durand et al., 2016)).

We suggest that a proline-dominated viscoelastic solution with NADES-like properties may form on a microscale level, locally, in slowly freeze-dehydrating larvae. As water molecules are osmotically driven from the gradually dehydrating hemolymph and tissues and migrate toward the extracellular ice during extracellular freezing, the highly soluble and relatively mobile proline molecules could move together with water until they reach a barrier of ice where they form a thin layer of concentrated proline, which we observed in MALDI-MSI. Such a viscoelastic liquid could contribute to the stabilization of deeply frozen *C. costata* larvae by forming a rubber-like zone between extracellular ice crystals and freeze-dehydrated tissues, thus reducing the thermo-mechanical stresses associated with temperature fluctuations during immersion in LN_2_ and rewarming (Rubinsky et al., 1980).

## Conclusions

Overall, our study reveals the composition of a specific mixture of putative cryoprotectants accumulated by the freeze-tolerant larvae of *C. costata*. We demonstrate in a bioassay that as little as three-days of exposure to diet enriched with artificial mixture of CPs allows an otherwise freeze-sensitive larva to survive not only deep freezing but also cryopreservation in LN_2_. The components of CP mixture act synergistically rather than additively to improve the freeze tolerance. These findings open avenues for functional experiments to better understand the mechanisms underlying cryoprotection and cryopreservation of complex multicellular organisms. Toward this goal, we demonstrate that the accumulated CPs suppress the melting point of body water, increase the fraction of osmotically inactive water, and significantly reduce the ice fraction. During slow extracellular freezing of whole larvae, trehalose becomes concentrated in the partially dehydrated hemolymph and tissues and stimulates glass transition, while proline moves to the boundary between extracellular ice and dehydrated tissues where it probably creates a layer of dense viscoelastic liquid. We suggest that these mechanisms contribute to larval extreme freeze tolerance and enable their cryopreservation by protecting their cells against thermomechanical shocks associated with freezing and transfers into and out of LN_2_.

## Supporting information

Appendice_MALDI MSI images

## Supplementary Information

(Supplementary Figures.pdf)

**Supplementary Figure A1** Phenotypic/acclimation variants and freeze tolerance bioassay.

**Supplementary Figure A2** Preparation of SDA-frozen larvae for MALDI-TOF/MS analysis.

**Supplementary Figure A3** Metabolomic profiles in *C. costata* tissues.

**Supplementary Figure A4** Sizes of metabolite pools in *C. costata* tissues.

**Supplementary Figure A5** MALDI-MSI of Gln, Asn and GPE on a longitudinal section of an SDA larva.

**Supplementary Figure A6** Relative quantification of MALDI-TOF signals: method and limitations.

**Supplementary Figure A7** Relative quantification of MALDI-TOF signals in LD larvae.

**Supplementary Figure A8** Relative quantification of MALDI-TOF signals in SDA larvae.

**Supplementary Figure A9** Relative quantification of MALDI-TOF signals in SDA-frozen larvae.

**Supplementary Figure A10** Analysis of DSC heating curves of trehalose solution (338 mmol.kg^-1^) in water.

**Supplementary Figure A11** Analysis of DSC heating curves of proline solution (978 mmol.kg^-1^) in water.

**Supplementary Figure A12** Analysis of DSC heating curves of glutamine solution (172 mmol.kg^-1^) in water.

**Supplementary Figure A13** Validation of trehalose m/z peak identity using ^13^C6 experiment.

**Supplementary Figure A14** Validation of proline m/z peak identity using ^13^C6 experiment.

(Supplemenatry Tables.xlsx)

**Supplementary Table A1** Tissue metabolomics results

**Supplementary Table A2** Results of DSC analysis

**Supplementary Table A3** CP mix 508 diet recipe

**Supplementary Table A4** Effect of CP mix 508 diet on larval survival and pupariation time

**Supplementary Table A5** Effect of CP Mix 508 diet on larval metabolite composition

**Supplementary Table A6** Effect of CP mix 508 and Pro 313 diets on LN2 survival

(Appendice.pdf)

**Supplementary Images** MALDI-MSI images of all transversal sections of *C. costata* larvae

## Acknowledgements

We thank Irena Vacková, Jana Járová, and Karolina Kowalska for maintenence of *C. costata* colonies, processing samples for metabolomics, and assistance during preparation of MALDI-MSI samples, respectively. This study was supported by a Grantová Agentura České Republiky (GAČR) grant no. 19-13381S to VK. Development of metabolomics platforms were supported by a GAČR grant no. 17-22276S to PŠ. The MALDI-MS analysis was supported by the Czech Academy of Science project no. RVO 68378050, Ministry of Education, Youth and Sports (MEYS) grant no. LM2018126, and MEYS in cooperation with European Structural and Investing Funds grant no. CZ.02.1.01/0.0/0.0/16_013/0001789 to RS. The DSC analysis was supported by the Ministry of Agriculture, grant number MZE RO0418 to MF.

## Author contributions

Conceptualization: VK

Experiments: MM, LK, JK, PV, LDM, RG, PH, JR, TŠ, MF

Formal analysis: VK, MM, LK

Funding acquisition. VK, PŠ, RS, MF

Supervision and methodology validation: VK, PŠ, RS, MF

Writing – original draft: VK

Writing – review and editing: all authors

## Notes

### Competing Interest Statement

The authors have declared no competing interest.

### Summary of Updates

This version of the manuscript has been revised just to change the formal layout of chapters.

