## Appendice_MALDI MSI images for "A mixture of innate cryoprotectants is key for freeze tolerance and cryopreservation of a drosophilid fly larva"

*Chymomyza costata*, 3rd instar larvae, 3-week-old

**phenotypic/acclimation variant LD (freeze sensitive)**

transversal sections through the middle part of larva

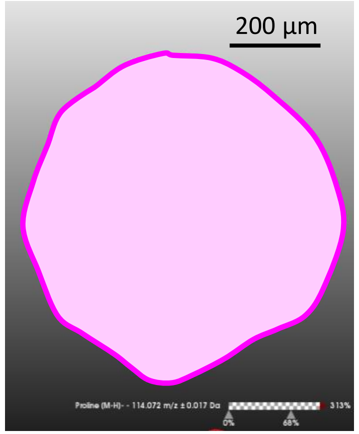


**H&E stain**

**Size bar** 200 µm

**Color scale**

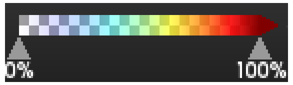


### Legend

For each section, the signal intensities of five putative CPs (proline, asparagine, glutamine, trehalose, and GPE) plus ATP are shown using arbitrary color scale overlaid over H&E stained transversal section. The same scale bar (200 µm applies to all sections). Medium denoising (SCiLS tool) of signal intensities among neighboring pixels of 10x10 µm was applied.

**Proline** [M-H]-
m/z 114,072 ± 0,017Da

**Asparagine** [M-H]-
m/z 131,062 ± 0,02Da

**Glutamine** [M-H]-
m/z 145,078 ± 0,022Da

**Trehalose** [M+Cl]-
m/z 377,101 ± 0,057Da

**GPE** [M-H]-
m/z 214,064 ± 0,032Da

**ATP**[M-H]-
m/z 506,017 ± 0,027Da

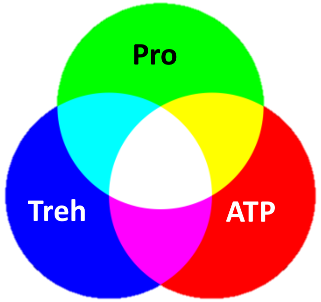


**Color mixing**

Next, signals of three select compounds (proline, trehalose, and ATP) are shown in color mixing mode.

**Pro** **ATP**

**Pro** **Treh**

**Treh** **ATP**

0258_LD-3A-06

Next, the raw signals (without denoising)
of putative CPs are shown in color mixing mode.

Pro Treh

GPE Treh

For comparison, the raw signal of ATP is shown
(no denoising)

ATP

Pro Gln

Pro Asn

Legend

# 0258_LD-3A-06

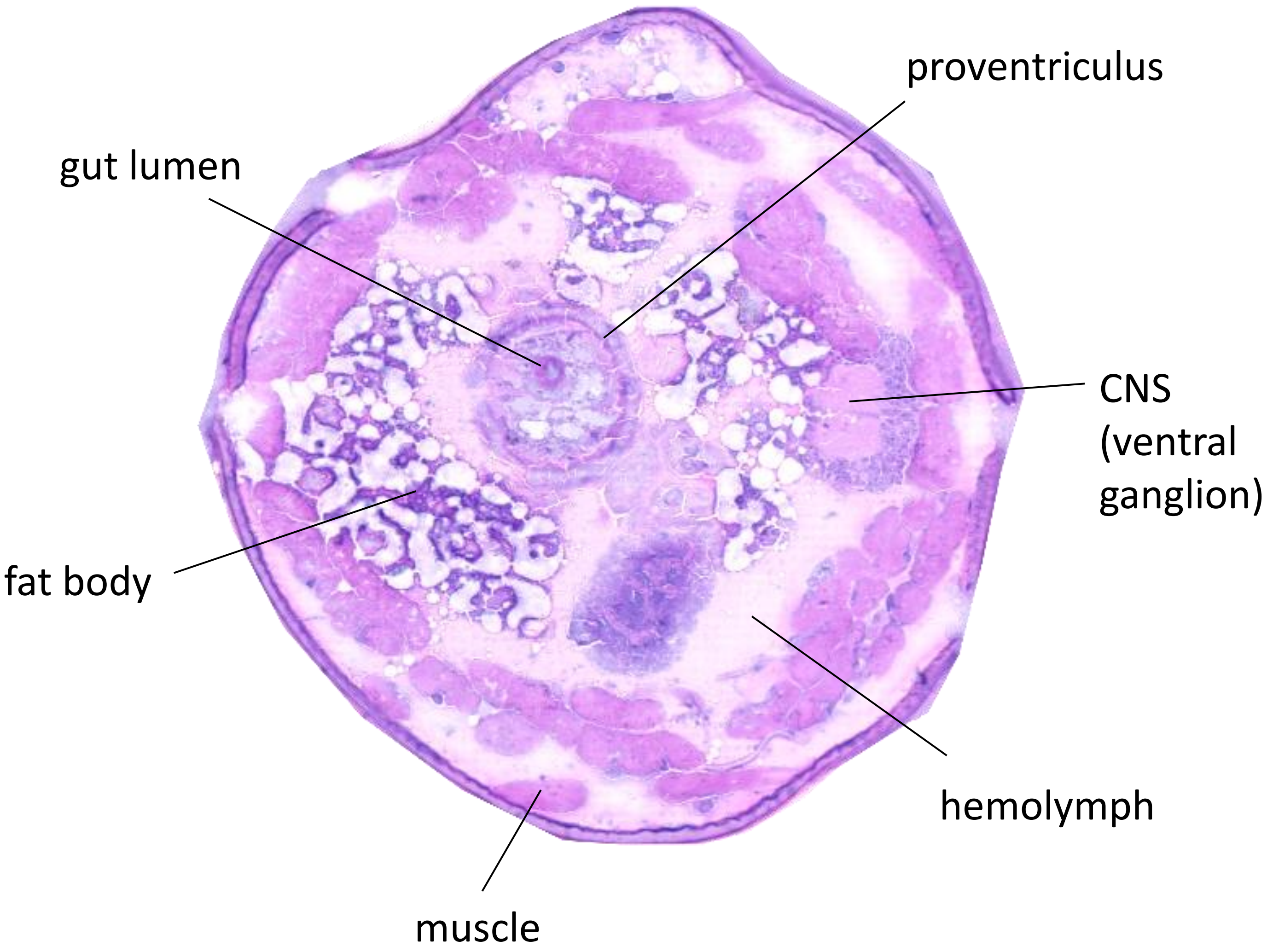
proventriculus
gut lumen
CNS
(ventral
ganglion)
fat body
hemolymph
muscle

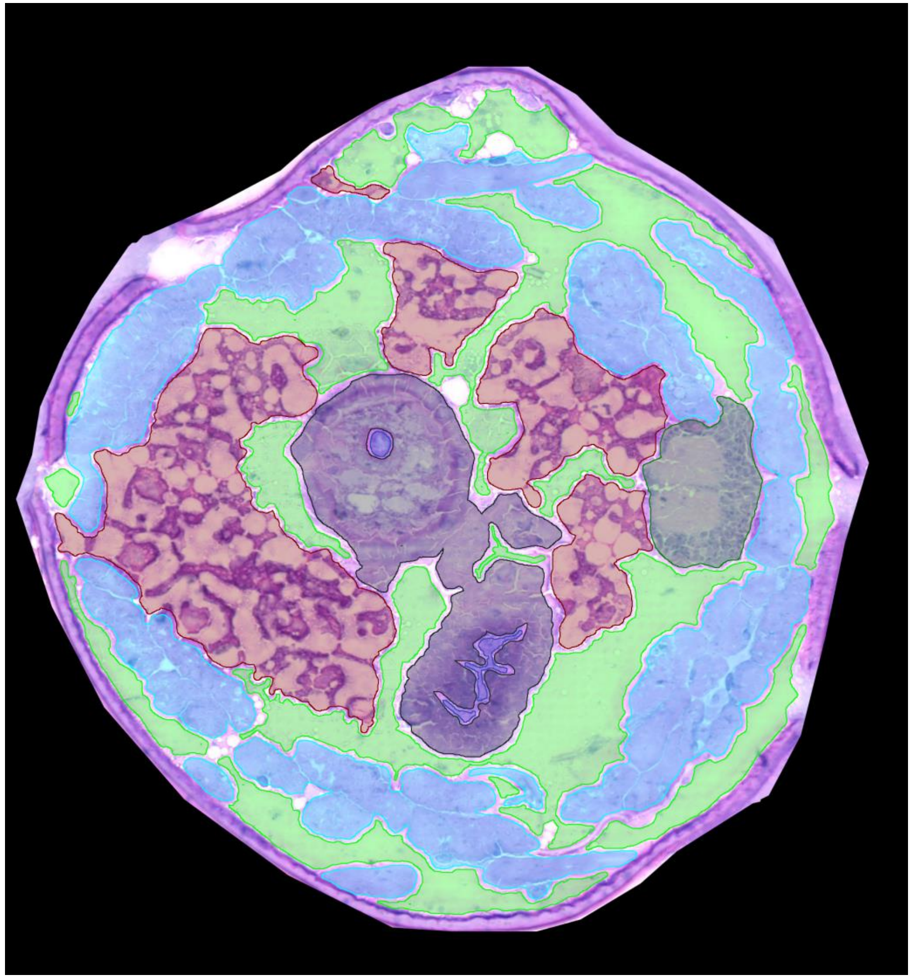

0258_LD-3A-06

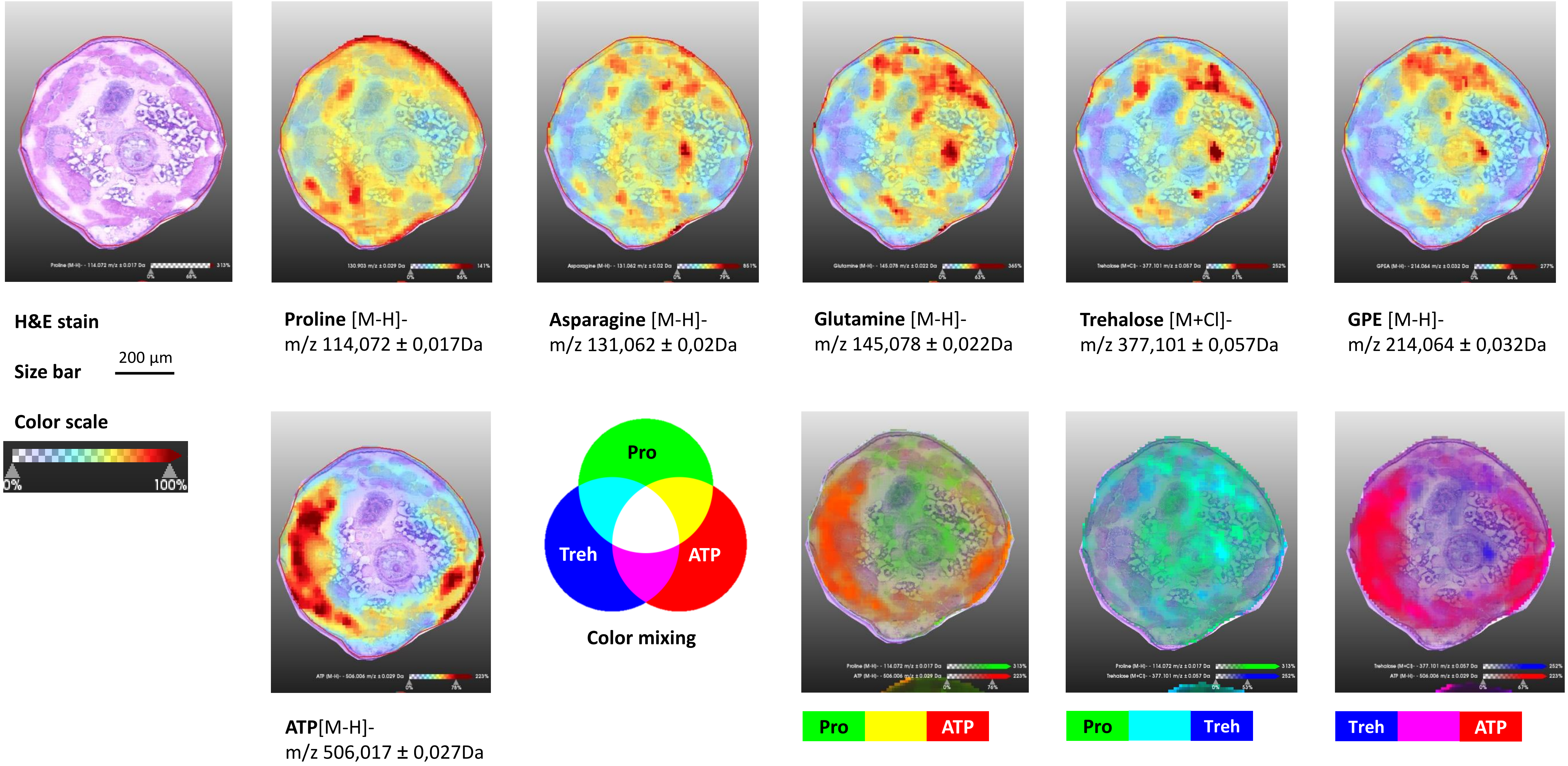
H&E stain
Size bar
200 µm
Color scale
0%
100%
Proline [M-H]-
m/z 114,072 ± 0,017Da
Asparagine [M-H]-
m/z 131,062 ± 0,02Da
Glutamine [M-H]-
m/z 145,078 ± 0,022Da
Trehalose [M+Cl]-
m/z 377,101 ± 0,057Da
GPE [M-H]-
m/z 214,064 ± 0,032Da
Pro
Treh
ATP
Color mixing
ATP[M-H]-
m/z 506,017 ± 0,027Da
Pro
ATP
Pro
Treh
Treh
ATP

0258_LD-3A-06

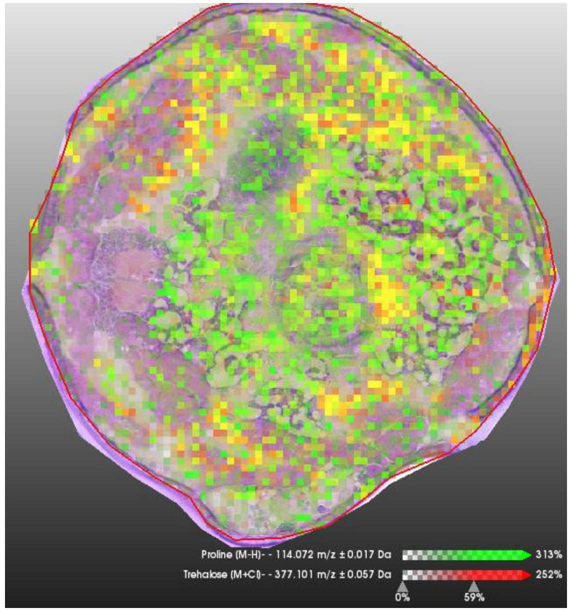


Pro Treh

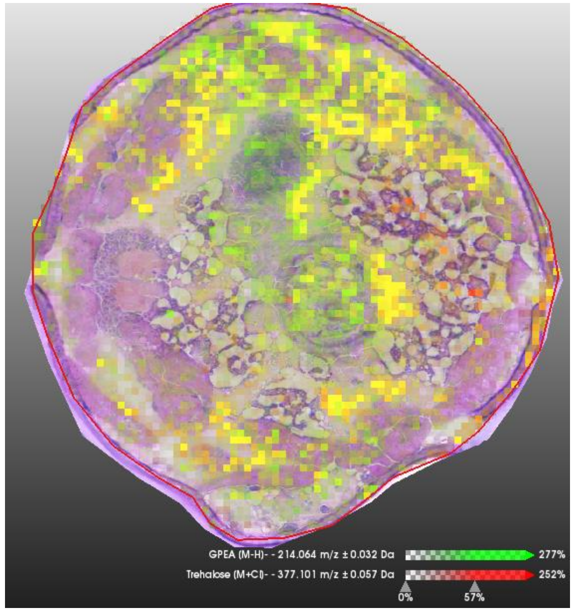


GPE Treh

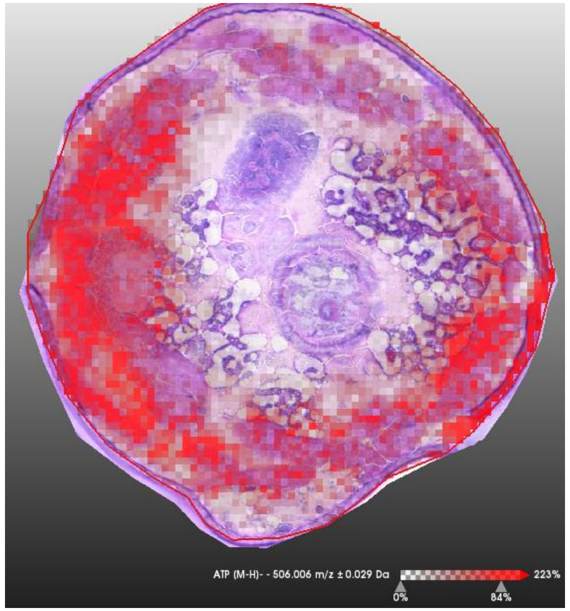


ATP

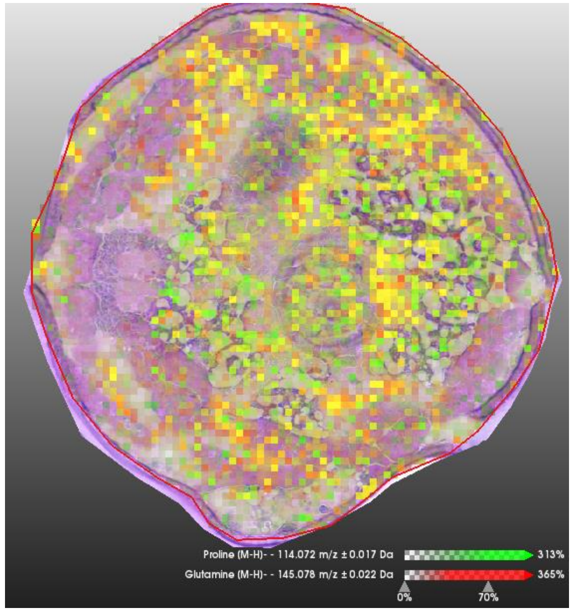


Pro Gln

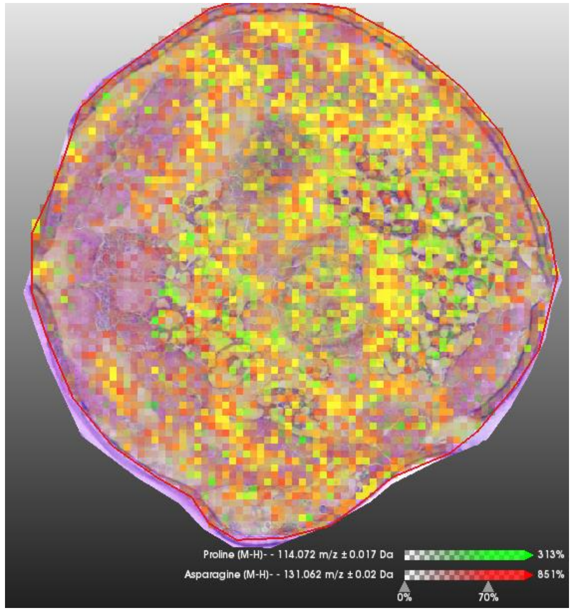


Pro Asn

# 0261_LD-2A-15

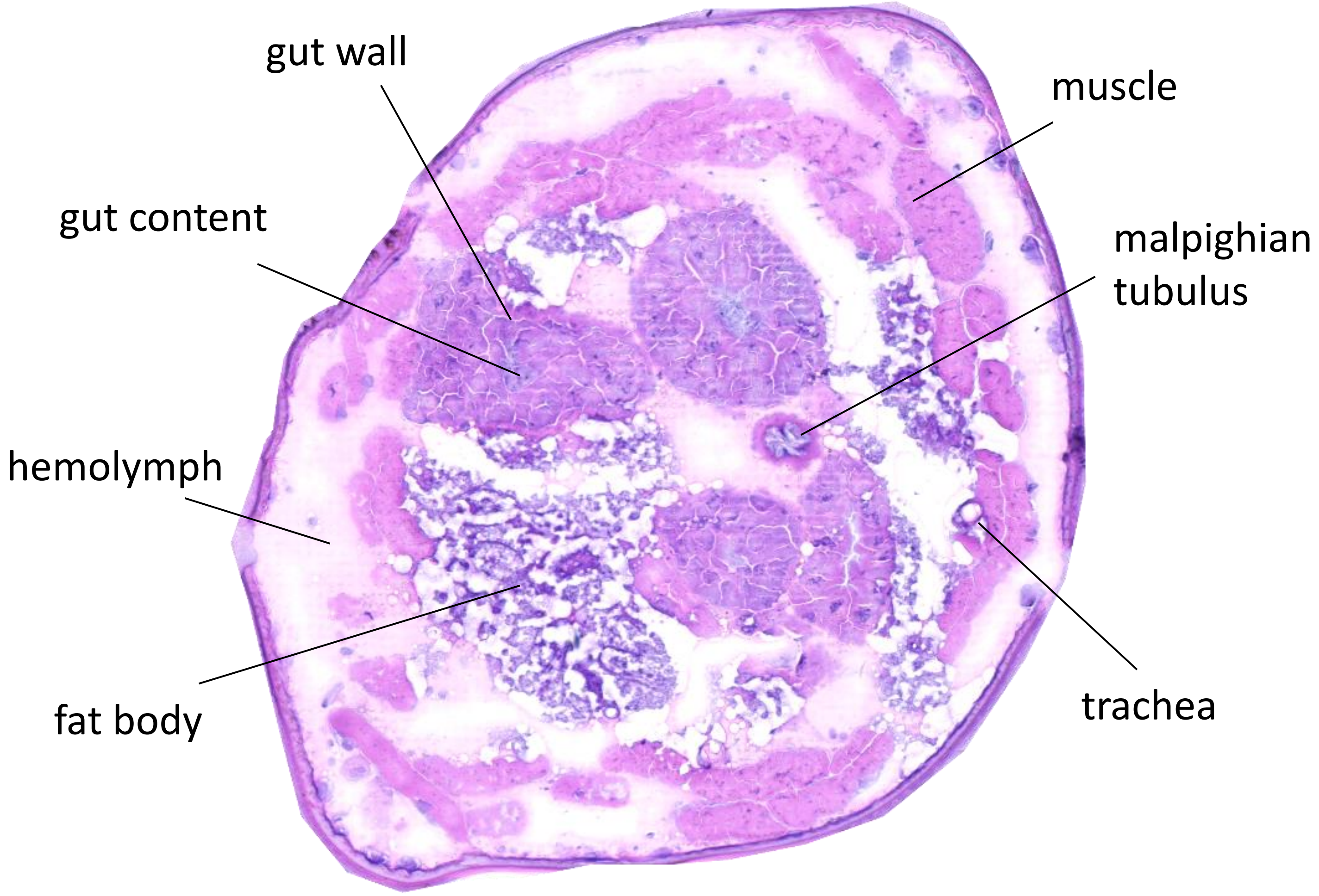

gut wall
muscle
gut content
malpighian tubulus
hemolymph
trachea
fat body


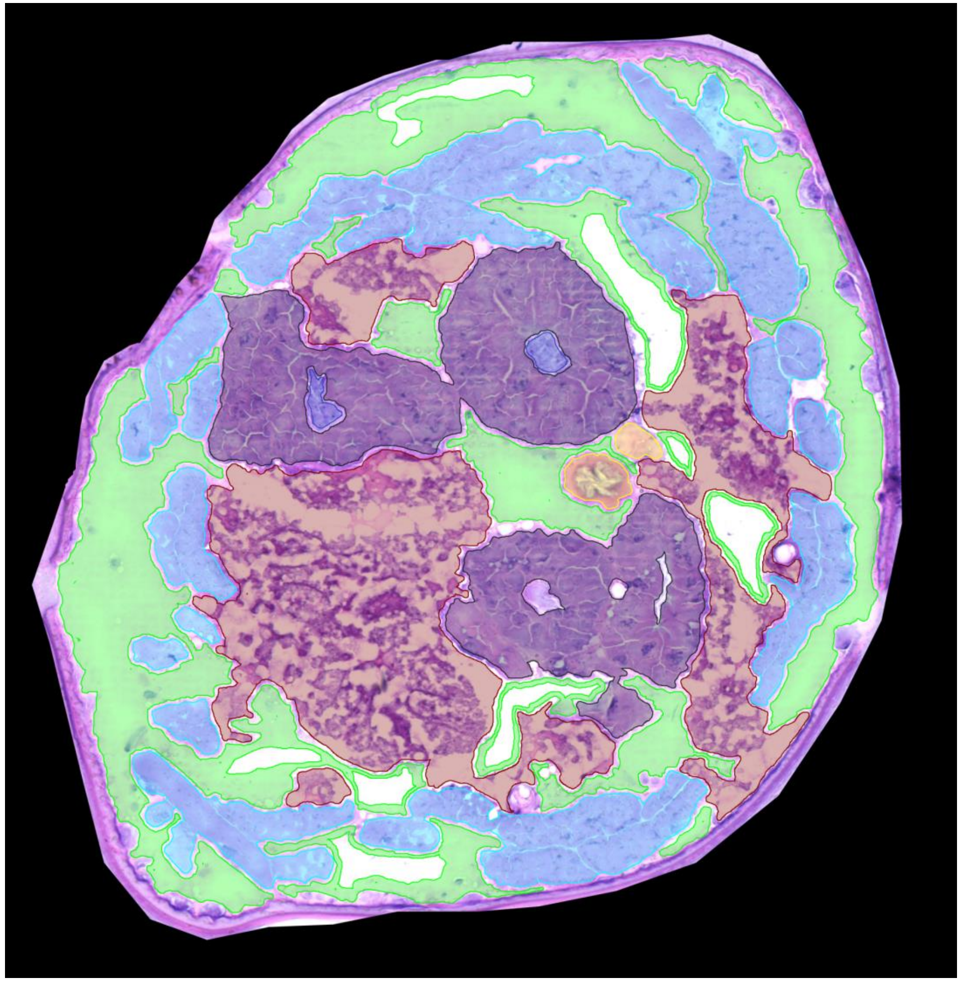

0261_LD-2A-15

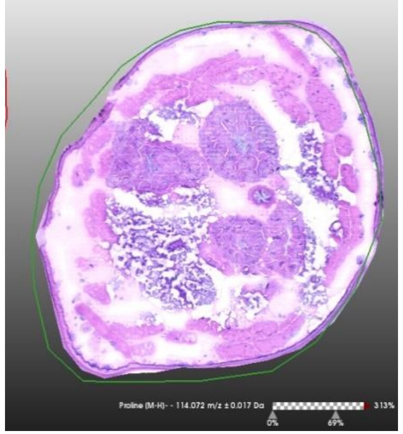

**H&E stain**

**Size bar** 200 µm

**Color scale**

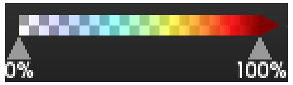


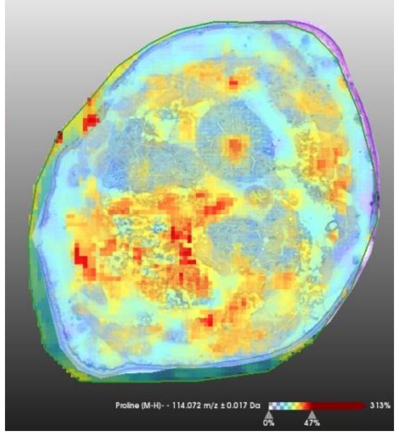

**Proline** [M-H]-
m/z 114,072 ± 0,017Da

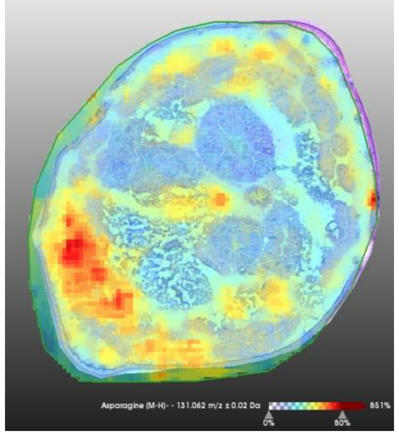

**Asparagine** [M-H]-
m/z 131,062 ± 0,02Da

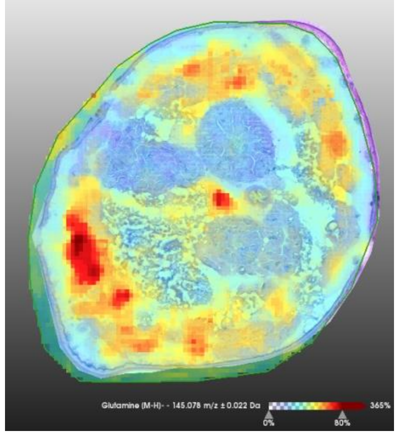

**Glutamine** [M-H]-
m/z 145,078 ± 0,022Da

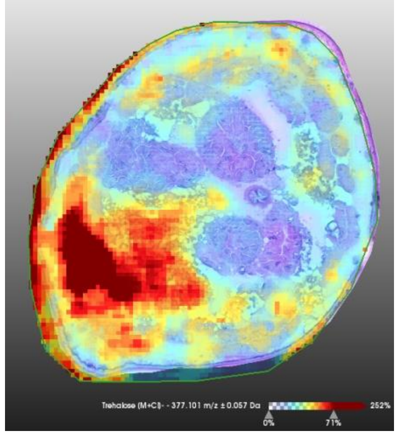

**Trehalose** [M+Cl]-
m/z 377,101 ± 0,057Da

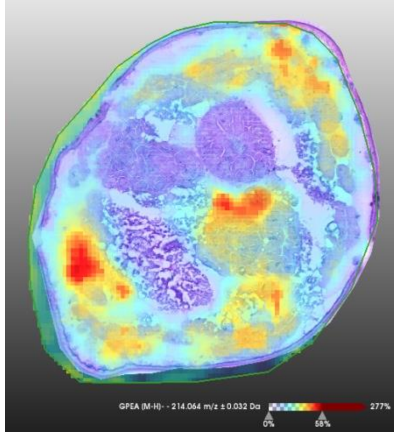

**GPE** [M-H]-
m/z 214,064 ± 0,032Da

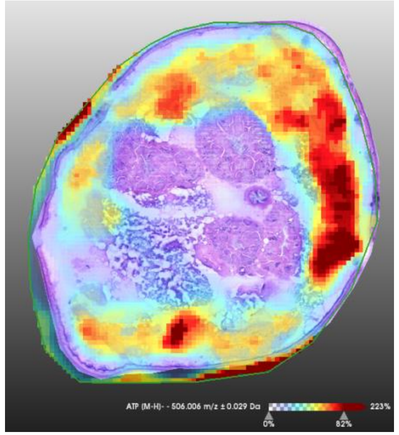

**ATP**[M-H]-
m/z 506,017 ± 0,027Da

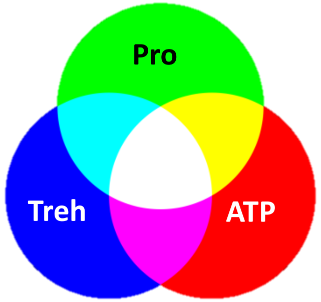


**Color mixing**

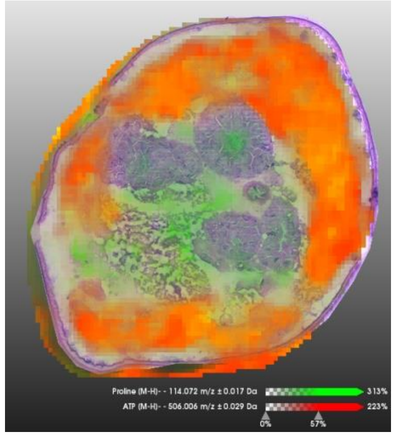

Pro | ATP

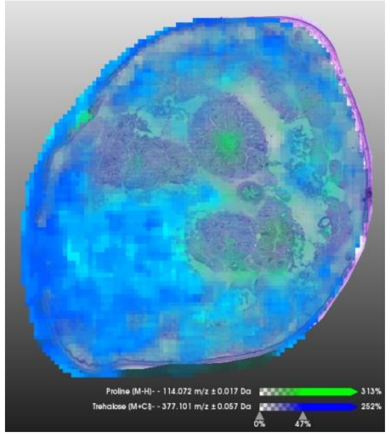

Pro | Treh

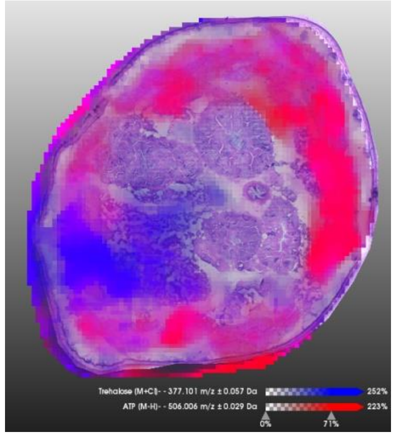

Treh | ATP

0261_LD-2A-15

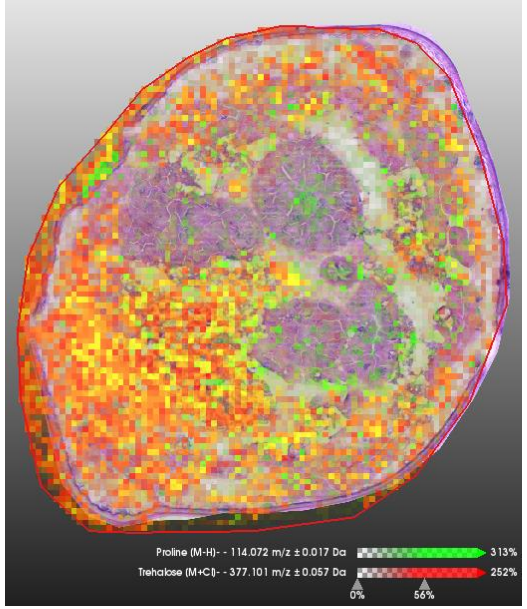


Pro | | Treh

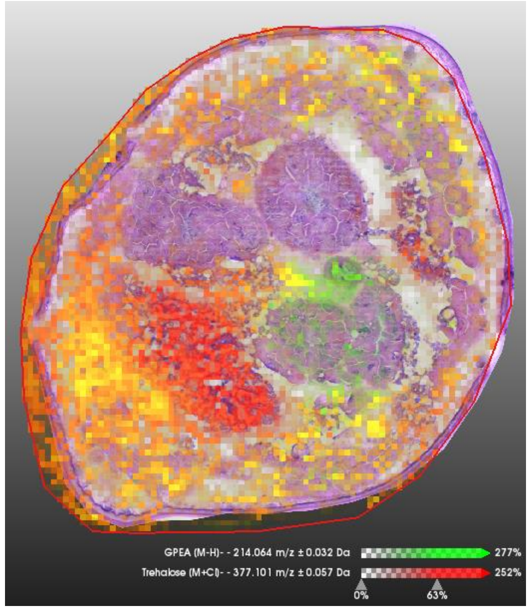


GPE | | Treh

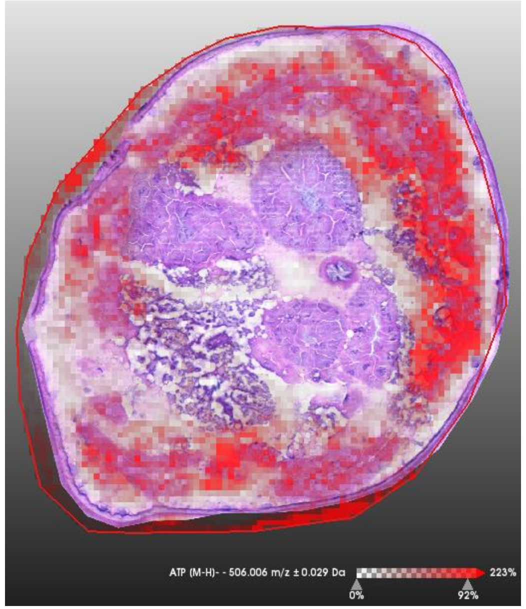


ATP

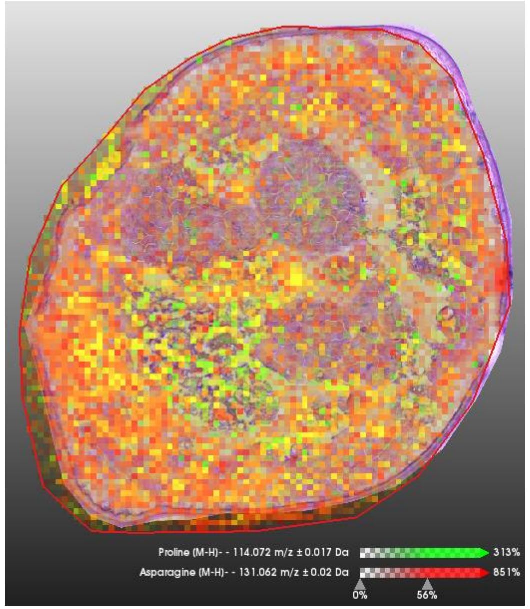


Pro | | Gln

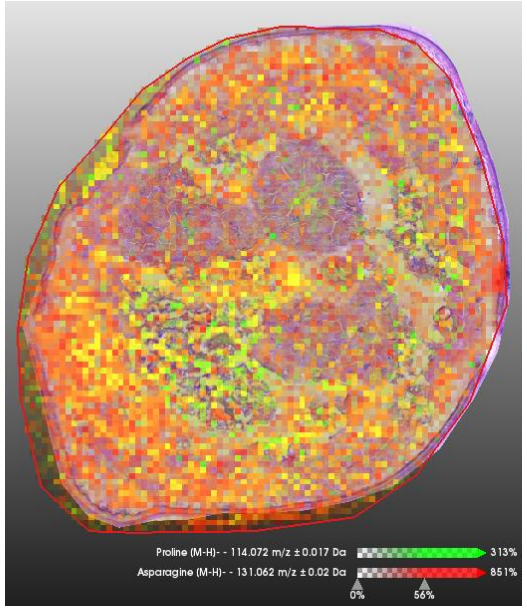


Pro | | Asn

# 0258_LD-3A-09

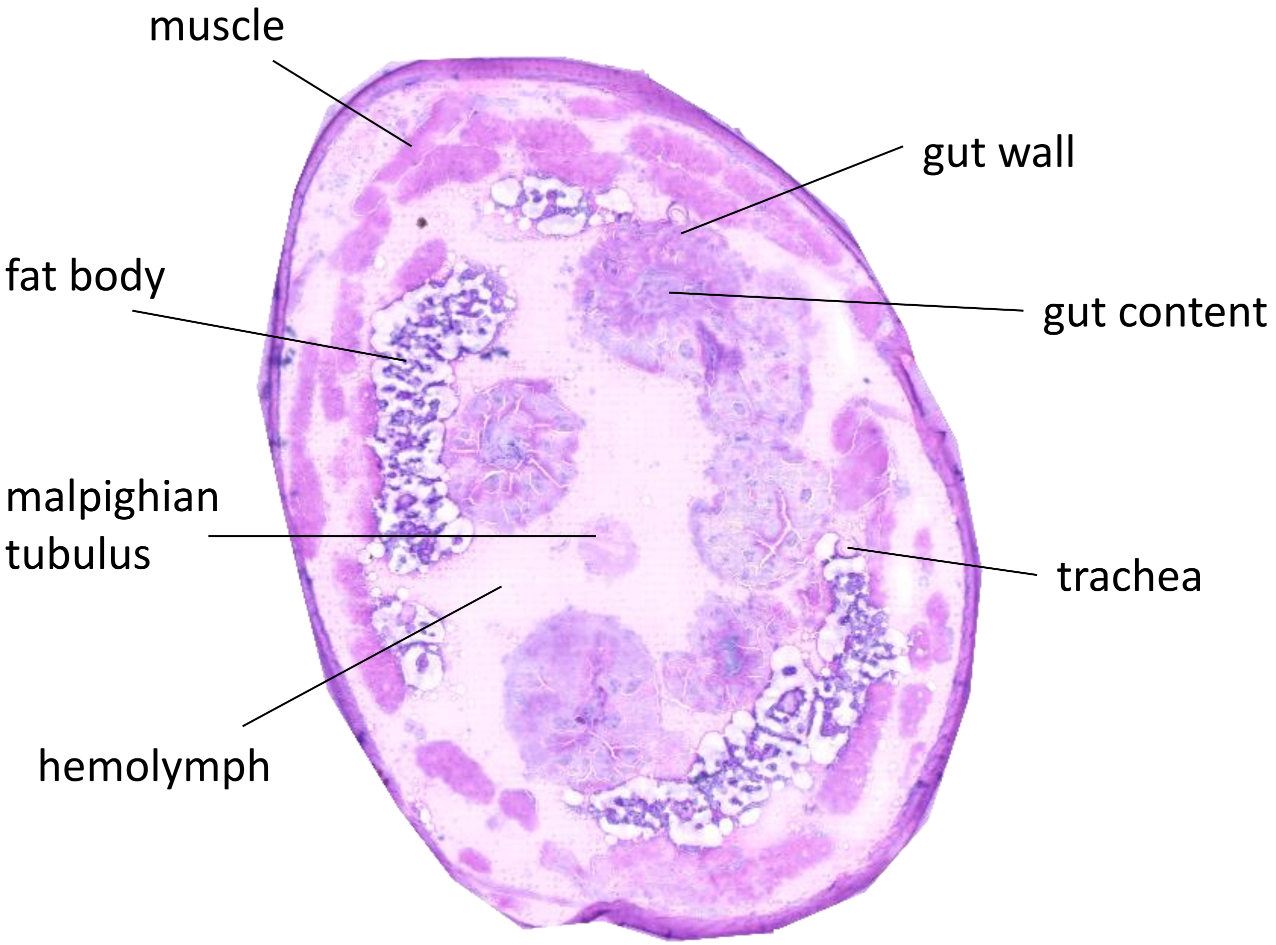
muscle
gut wall
fat body
gut content
malpighian
tubulus
trachea
hemolymph

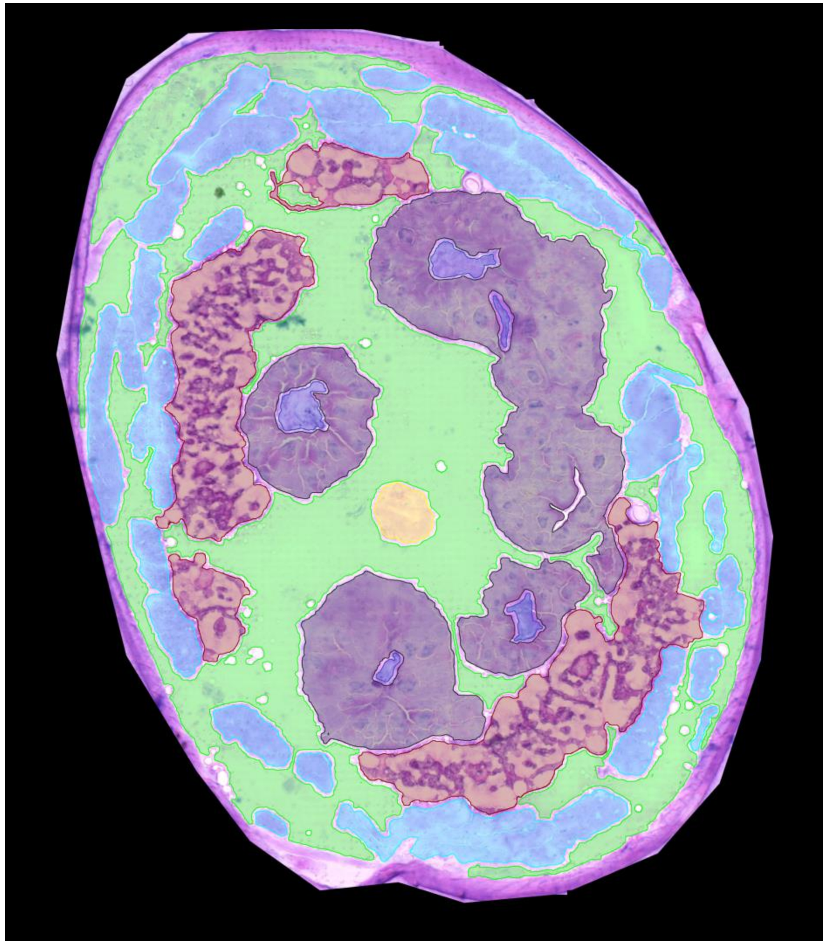

0258_LD-3A-09

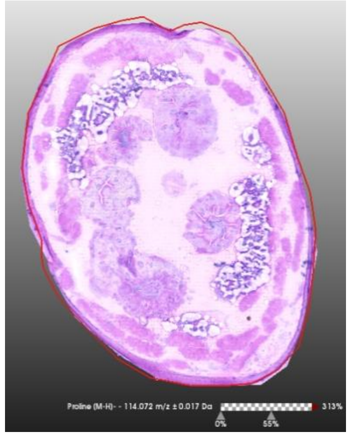

**H&E stain**

**Size bar** 200 µm

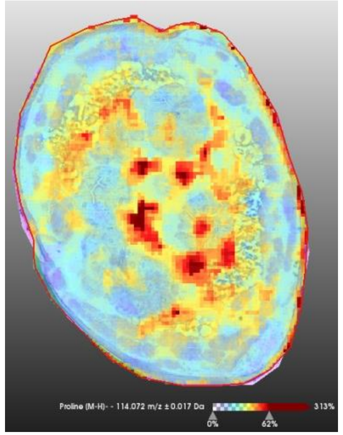

**Proline** [M-H]-
m/z 114,072 ± 0,017Da

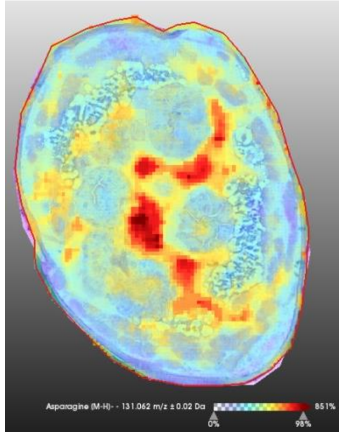

**Asparagine** [M-H]-
m/z 131,062 ± 0,02Da

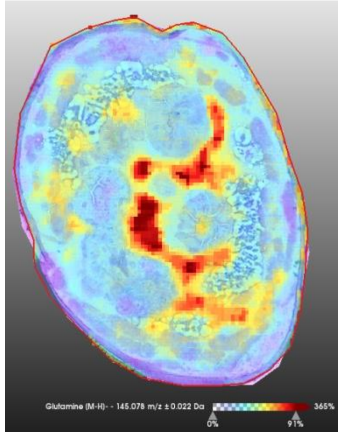

**Glutamine** [M-H]-
m/z 145,078 ± 0,022Da

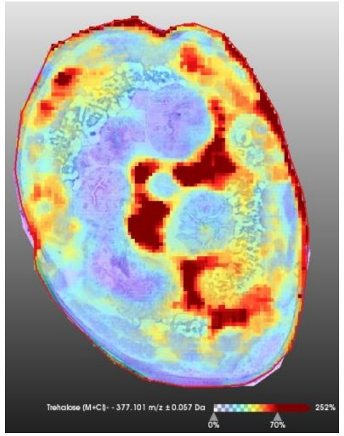

**Trehalose** [M+Cl]-
m/z 377,101 ± 0,057Da

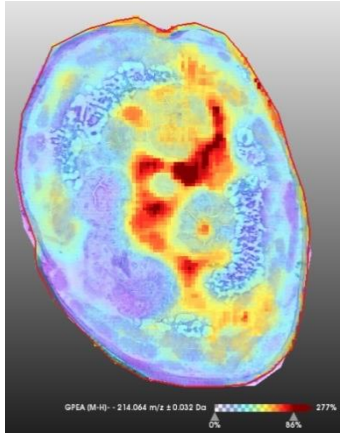

**GPE** [M-H]-
m/z 214,064 ± 0,032Da

**Color scale**

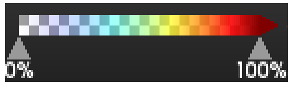


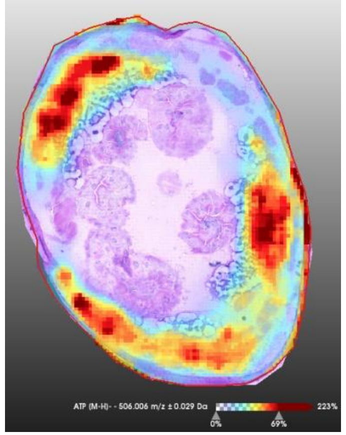

**ATP**[M-H]-
m/z 506,017 ± 0,027Da

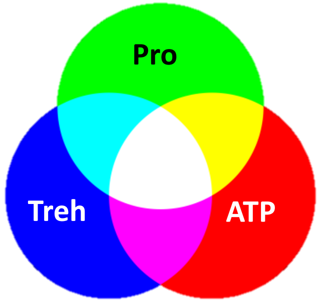


**Color mixing**

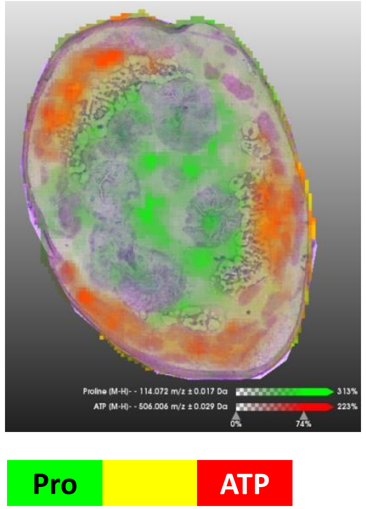


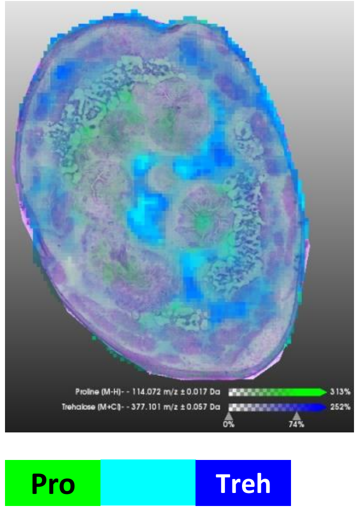


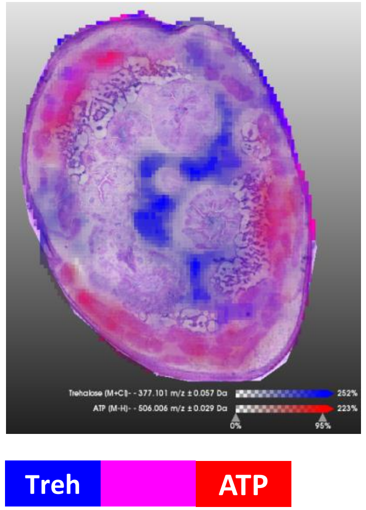

0258_LD-3A-09

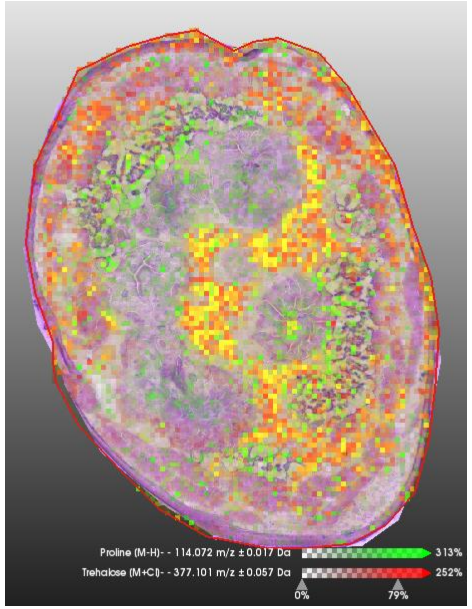


Pro Treh

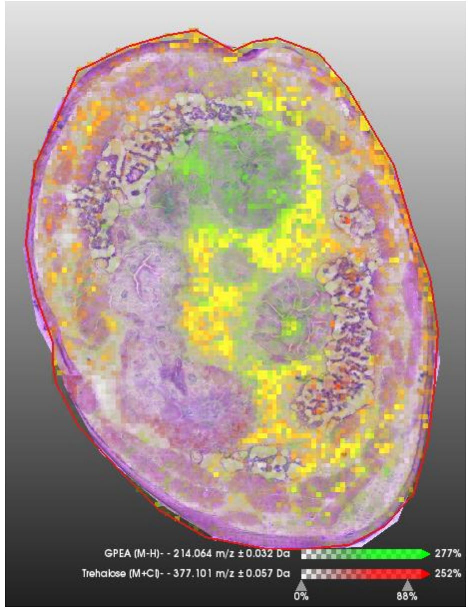


GPE Treh

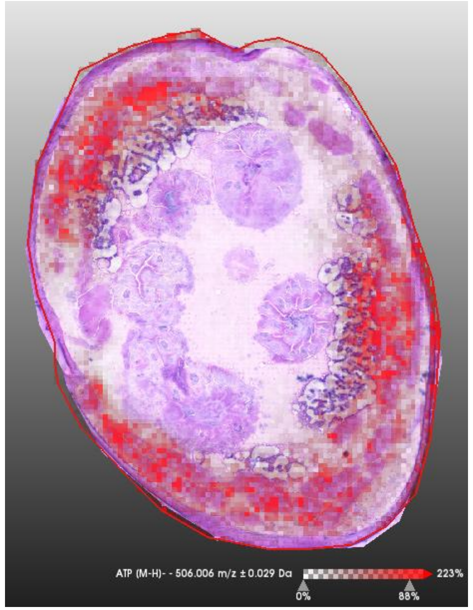


ATP

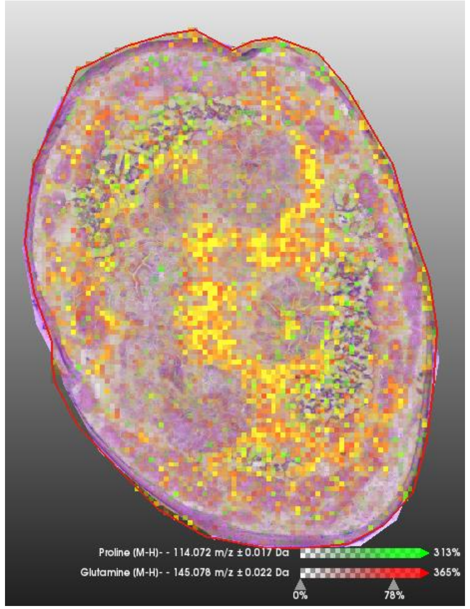


Pro Gln

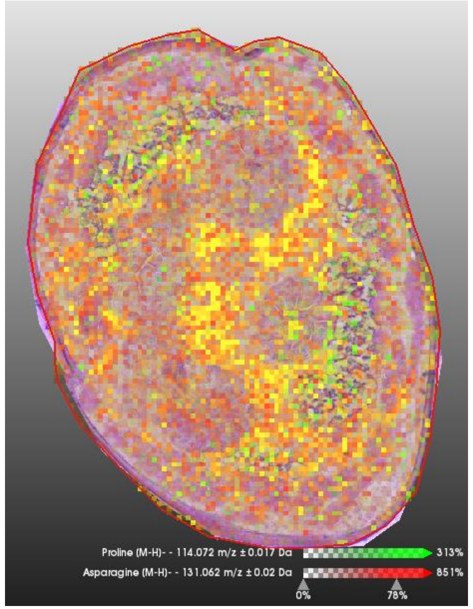


Pro Asn

# 0258_LD-3A-10

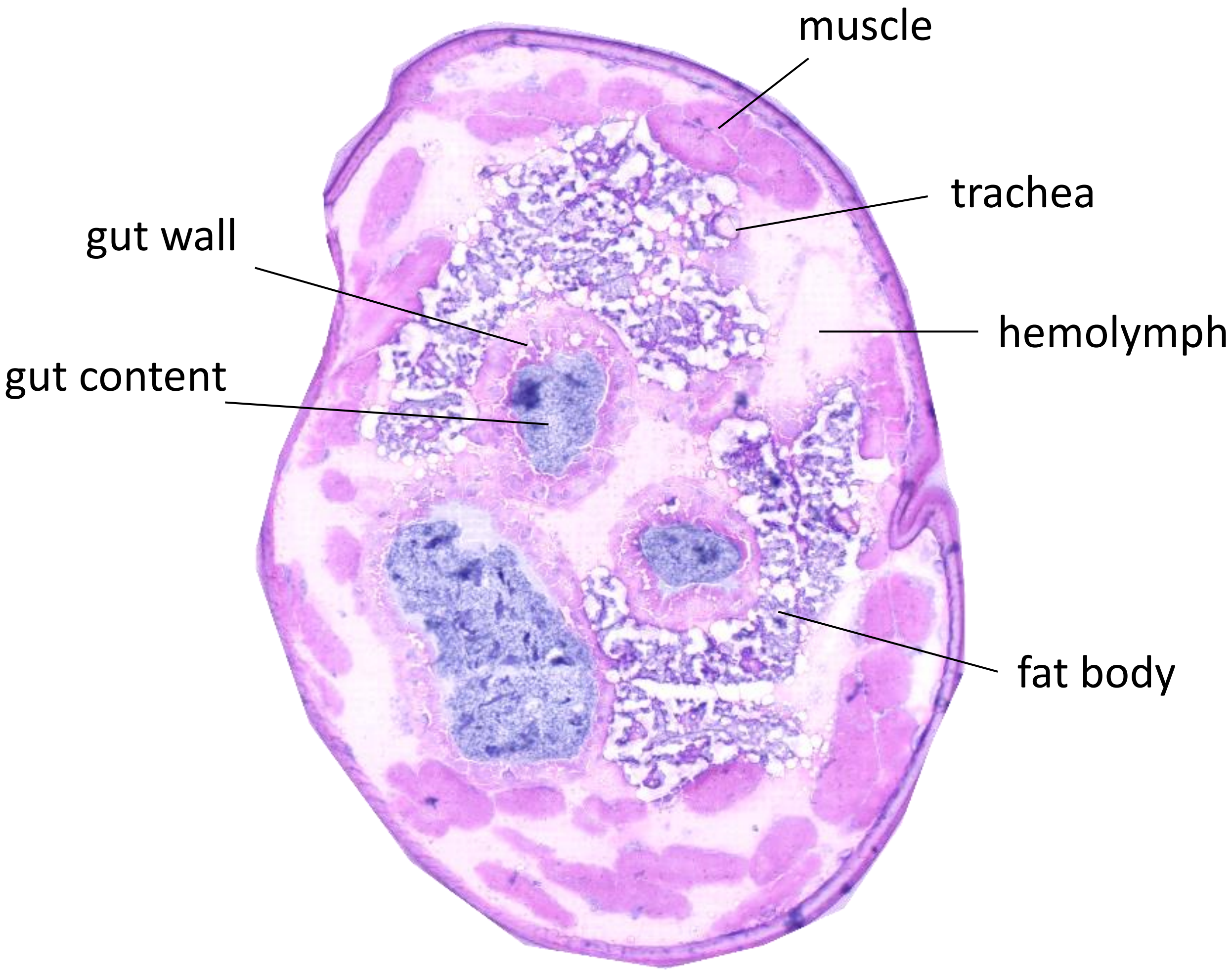


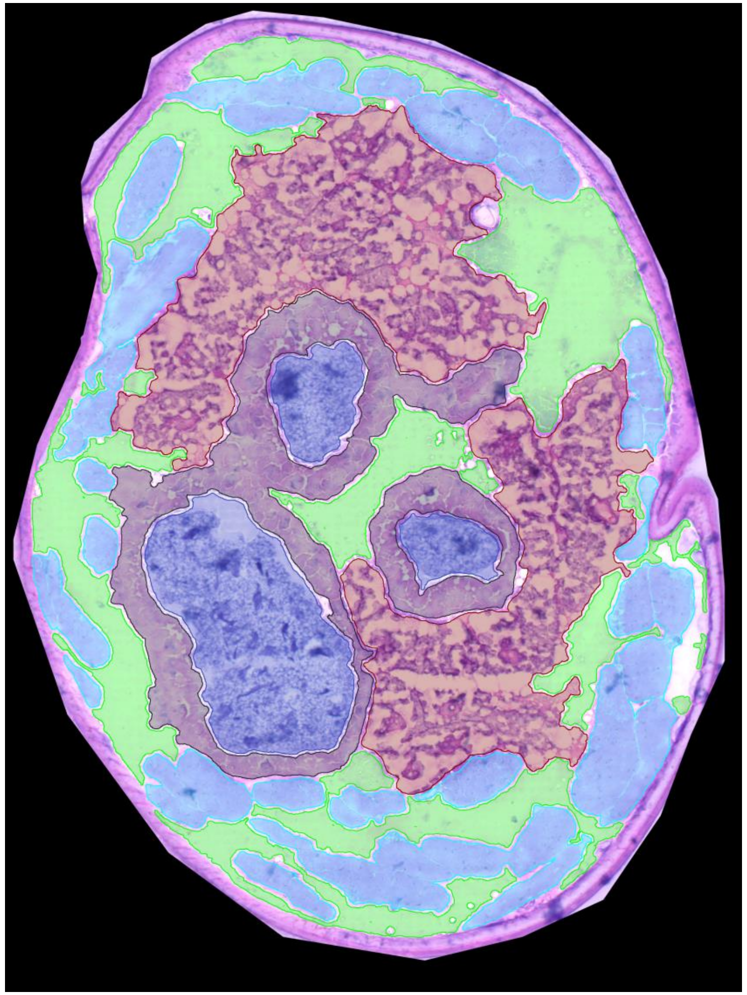

0258_LD-3A-10

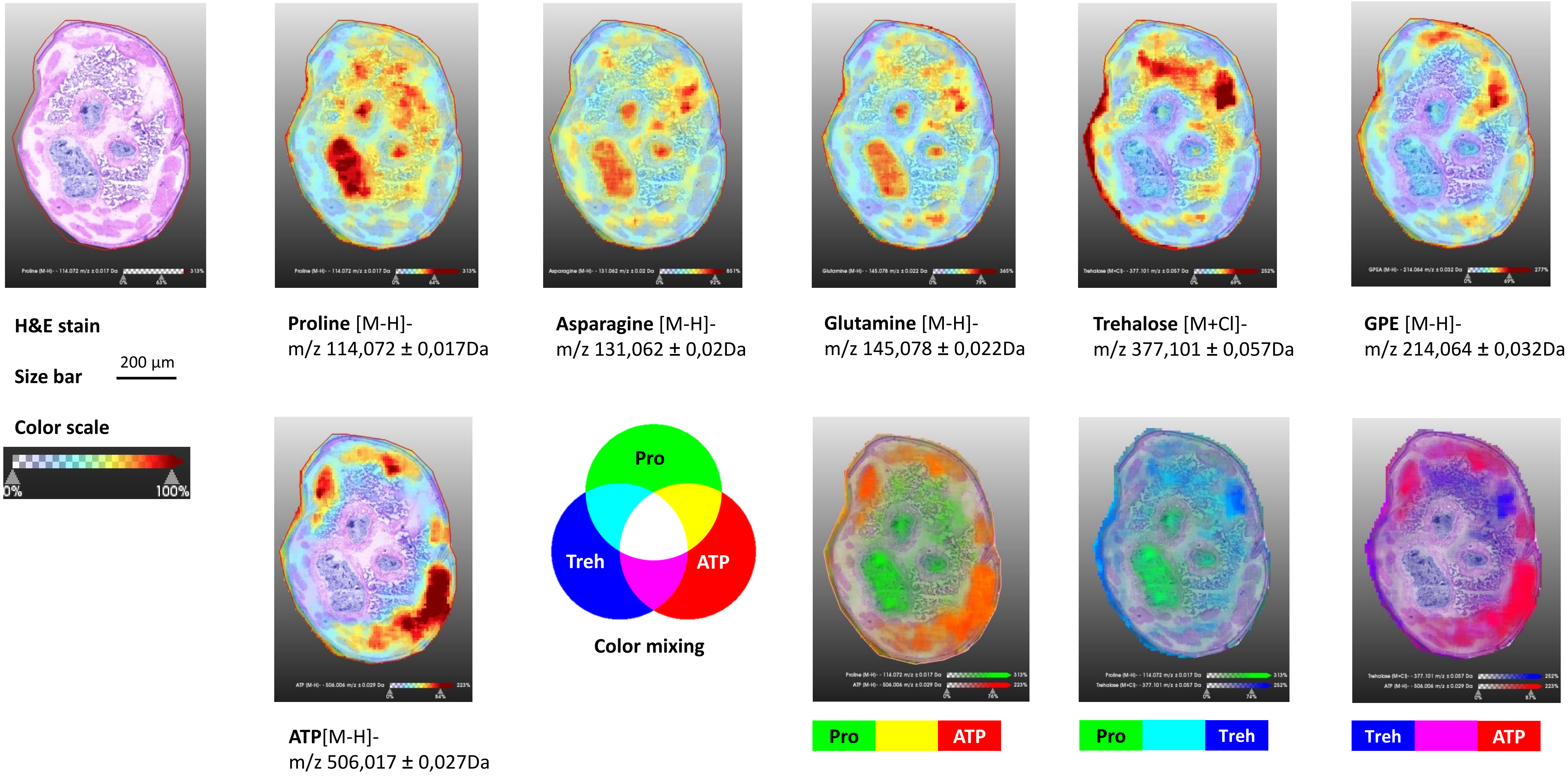
H&E stain
Size bar
200 µm
Color scale
0%
100%
Proline [M-H]-
m/z 114,072 ± 0,017Da
Asparagine [M-H]-
m/z 131,062 ± 0,02Da
Glutamine [M-H]-
m/z 145,078 ± 0,022Da
Trehalose [M+Cl]-
m/z 377,101 ± 0,057Da
GPE [M-H]-
m/z 214,064 ± 0,032Da
ATP[M-H]-
m/z 506,017 ± 0,027Da
Pro
Treh
ATP
Color mixing
Pro
ATP
Pro
Treh
Treh
ATP

0258_LD-3A-10

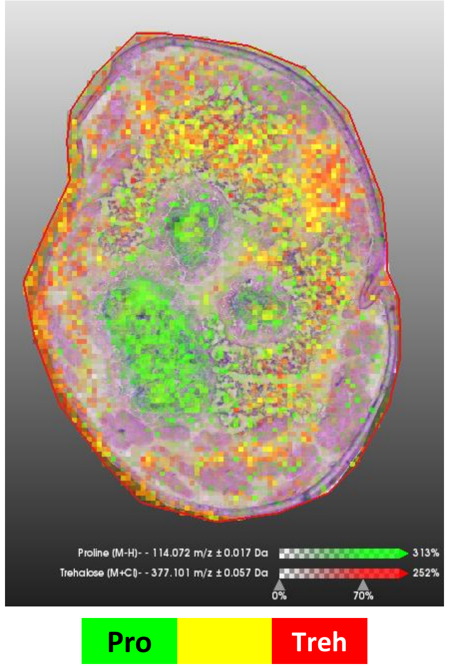

Proline (M-H)- - 114.072 m/z ±0.017 Da
313%
Trehalose (M+Cl)- - 377.101 m/z ±0.057 Da
252%
0%
70%
Pro
Treh


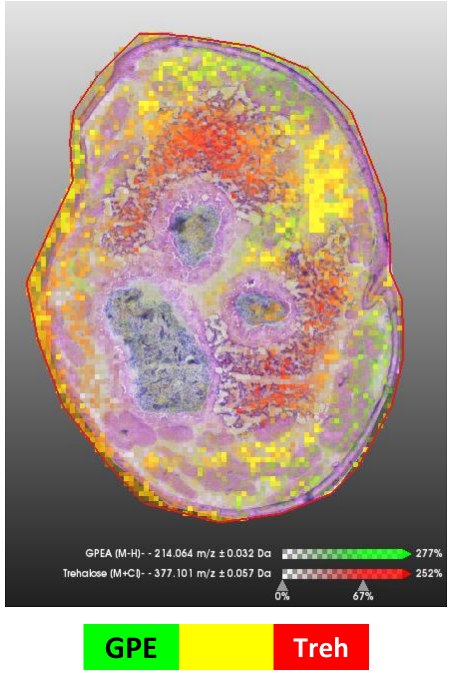

GPEA (M-H)- - 214.064 m/z ±0.032 Da
277%
Trehalose (M+Cl)- - 377.101 m/z ±0.057 Da
252%
0%
67%
GPE
Treh


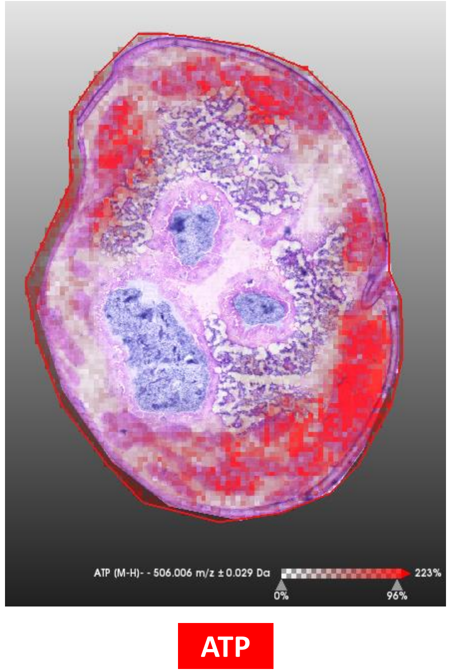

ATP (M-H)- - 506.006 m/z ±0.029 Da
223%
0%
96%
ATP


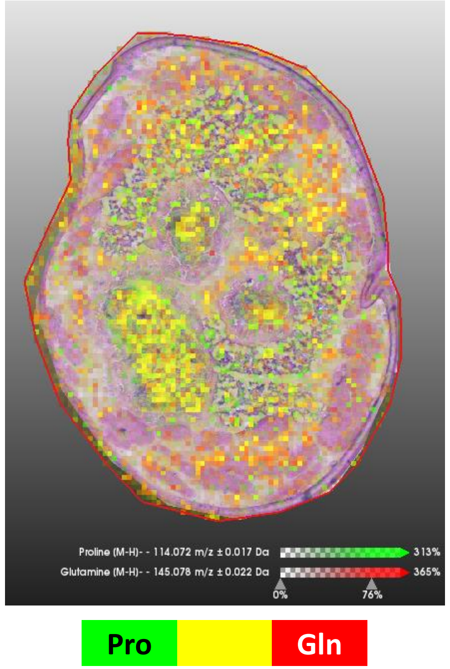

Proline (M-H)- - 114.072 m/z ±0.017 Da
313%
Glutamine (M-H)- - 145.078 m/z ±0.022 Da
365%
0%
76%
Pro
Gln


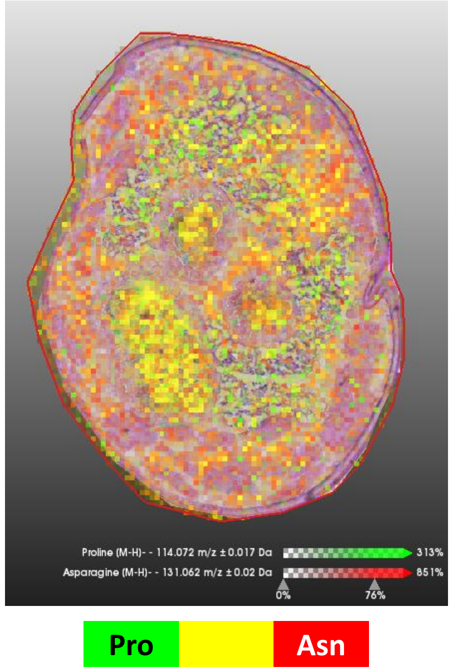

Proline (M-H)- - 114.072 m/z ±0.017 Da
313%
Asparagine (M-H)- - 131.062 m/z ±0.02 Da
851%
0%
76%
Pro
Asn

# 0261_LD-2A-17

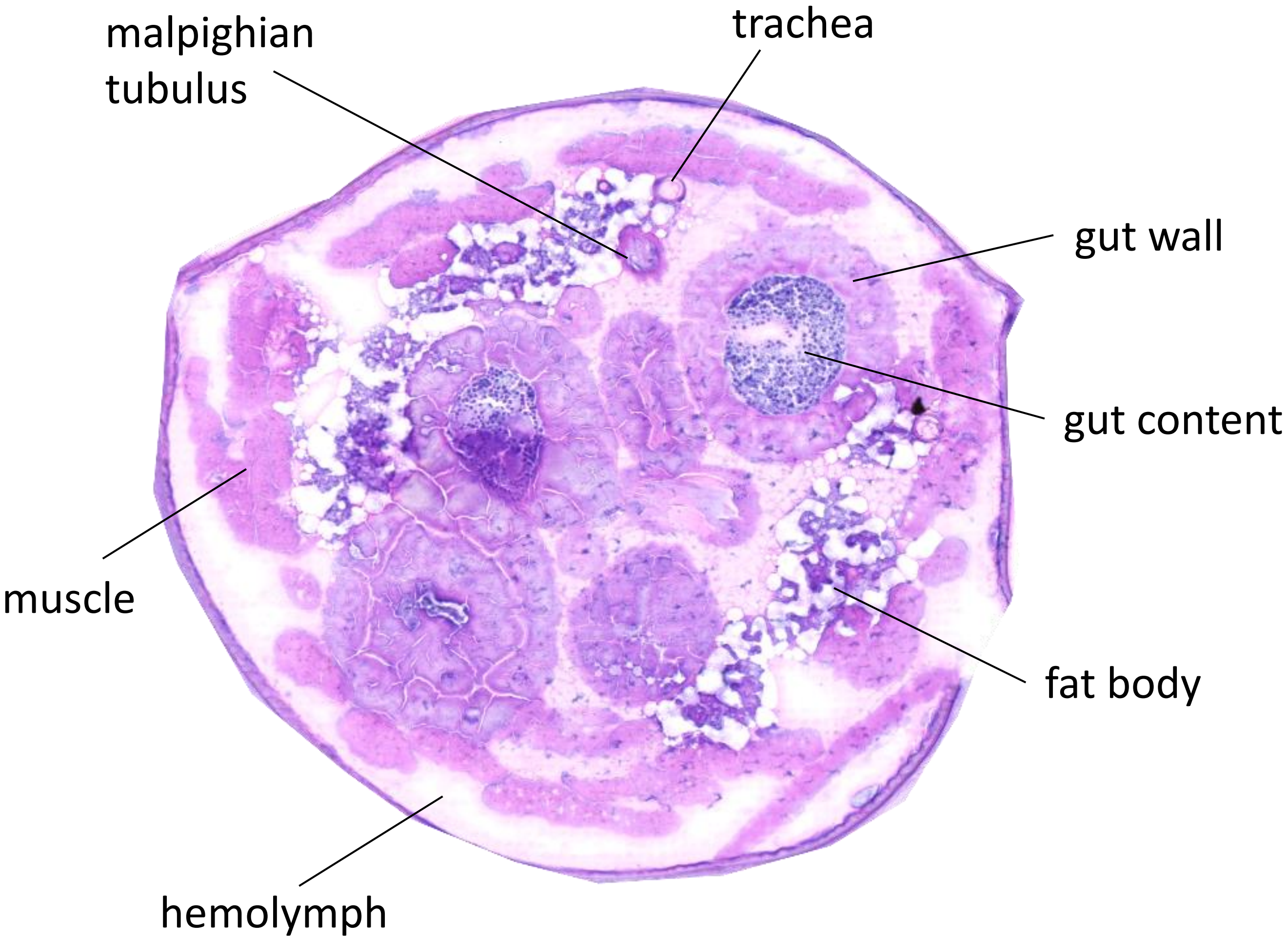
malpighian
tubulus
trachea
gut wall
gut content
muscle
fat body
hemolymph

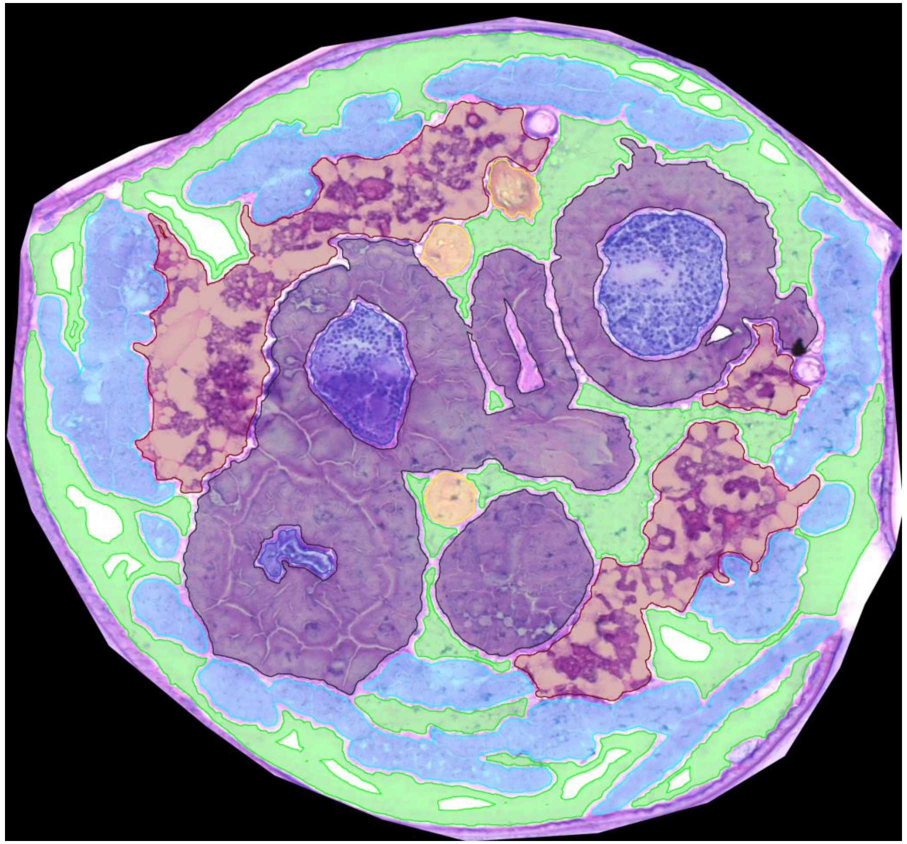

0261_LD-2A-17

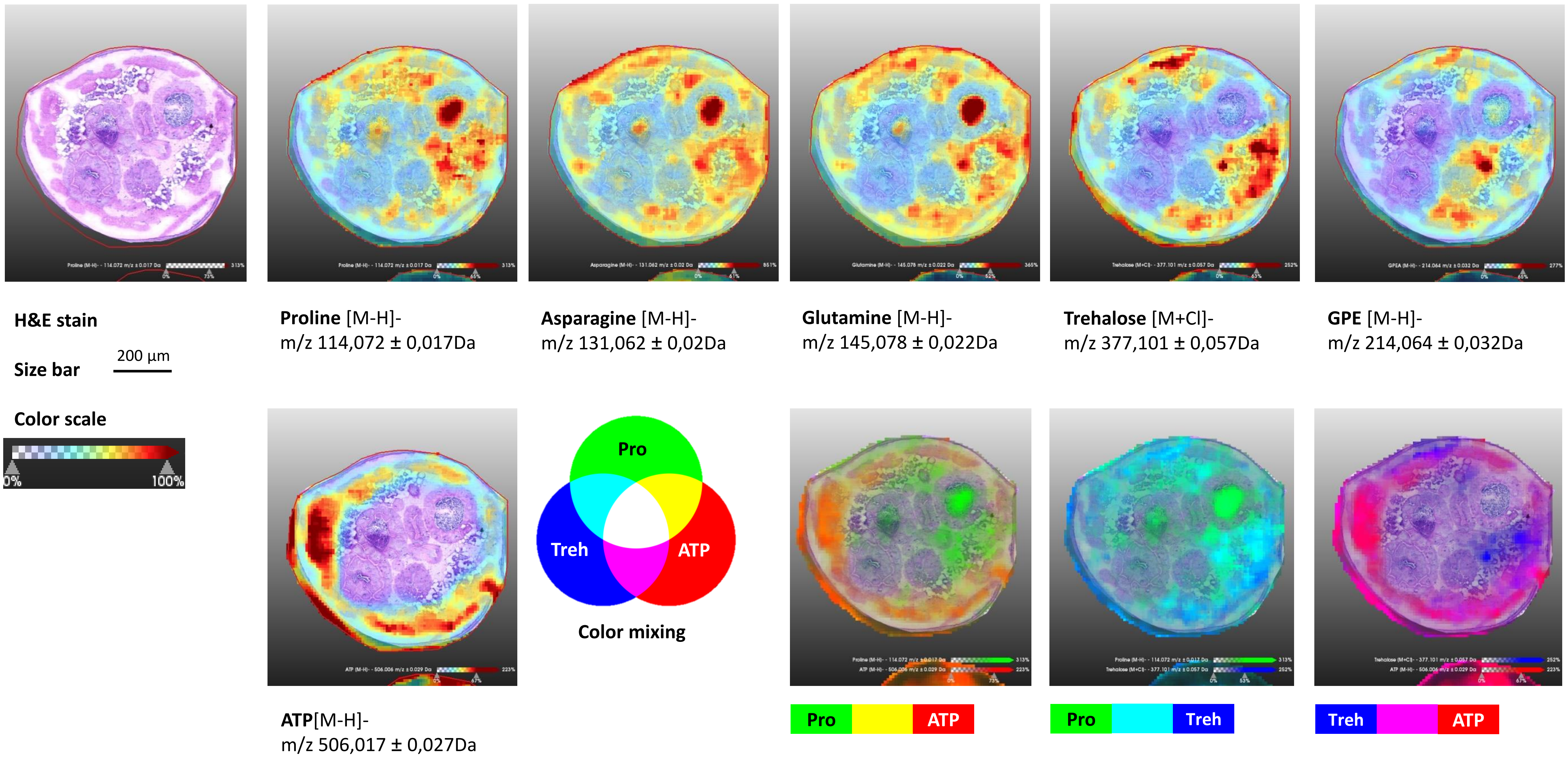

H&E stain
Size bar
200 µm
Color scale
0%
100%
Proline [M-H]-
m/z 114,072 ± 0,017Da
Asparagine [M-H]-
m/z 131,062 ± 0,02Da
Glutamine [M-H]-
m/z 145,078 ± 0,022Da
Trehalose [M+Cl]-
m/z 377,101 ± 0,057Da
GPE [M-H]-
m/z 214,064 ± 0,032Da
ATP[M-H]-
m/z 506,017 ± 0,027Da
Pro
Treh
ATP
Color mixing
Pro
ATP
Pro
Treh
Treh
ATP

0261_LD-2A-17

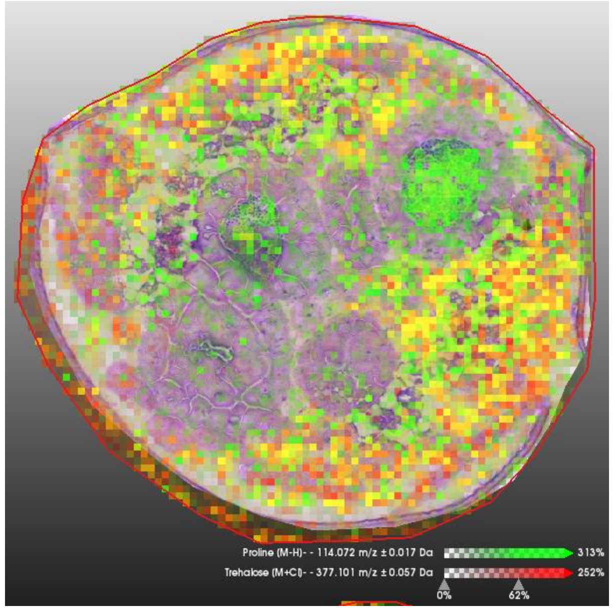


Pro Treh

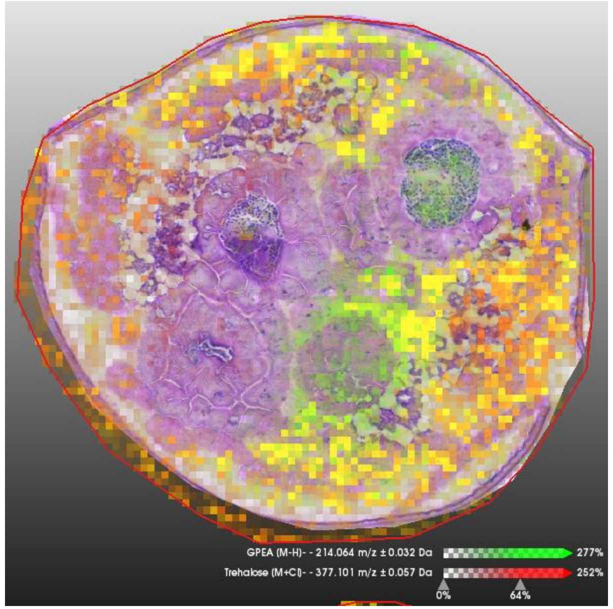


GPE Treh

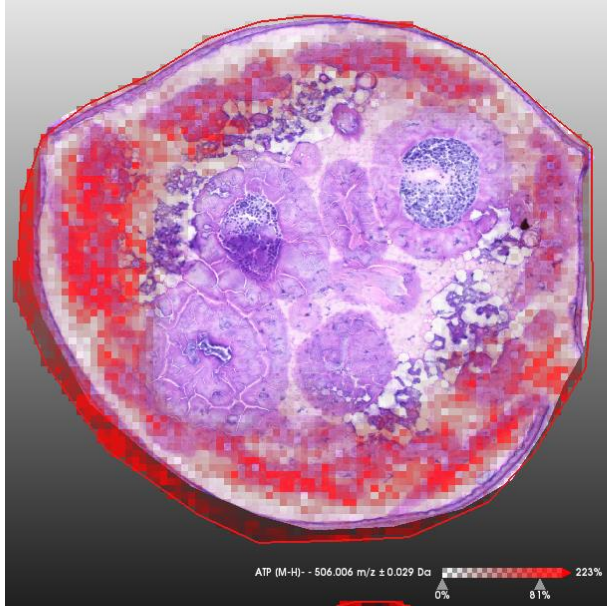


ATP

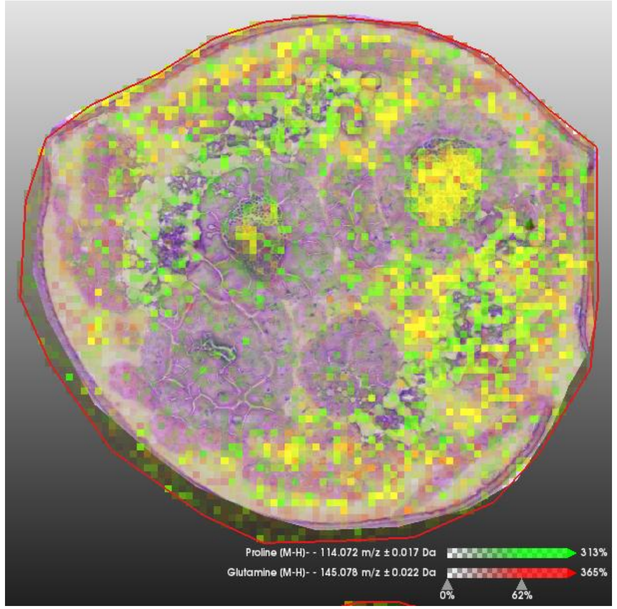


Pro Gln

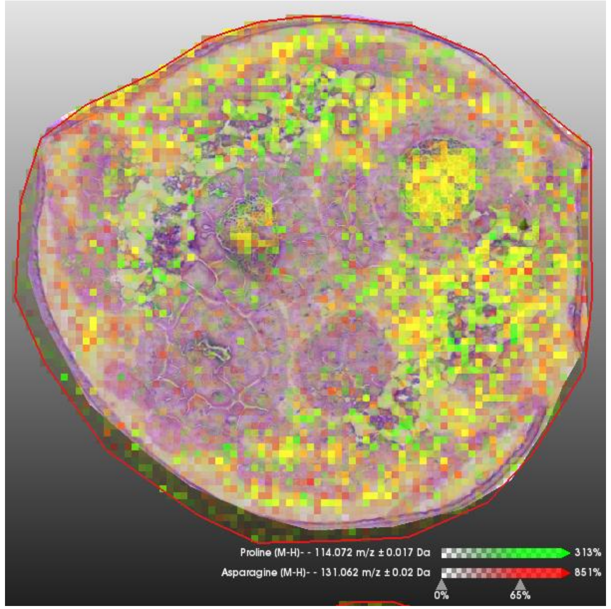


Pro Asn

*Chymomyza costata*, 3rd instar larvae, 3-week-old

**phenotypic/acclimation variant SDA (freeze tolerant)**
prior to slow freezing to -30°C

transversal sections through the middle part of larva

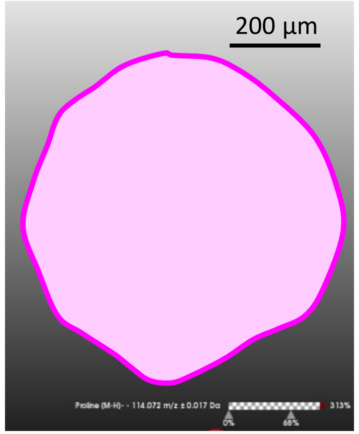


**H&E stain**

**Size bar** 200 µm

**Color scale**

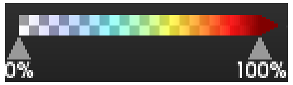


For each section, the signal intensities of five putative CPs (proline, asparagine, glutamine, trehalose, and GPE) plus ATP are shown using arbitrary color scale overlaid over H&E stained transversal section. The same scale bar (200 µm applies to all sections). Medium denoising (SCiLS tool) of signal intensities among neighboring pixels of 10x10 µm was applied.

**Proline** [M-H]-
m/z 114,072 ± 0,017Da

**Asparagine** [M-H]-
m/z 131,062 ± 0,02Da

**Glutamine** [M-H]-
m/z 145,078 ± 0,022Da

**Trehalose** [M+Cl]-
m/z 377,101 ± 0,057Da

**GPE** [M-H]-
m/z 214,064 ± 0,032Da

**ATP**[M-H]-
m/z 506,017 ± 0,027Da

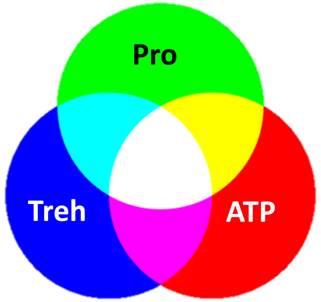


**Color mixing**

Next, signals of three select compounds (proline, trehalose, and ATP) are shown in color mixing mode.

**Pro** **ATP**

**Pro** **Treh**

**Treh** **ATP**

### Legend

0258_LD-3A-06

Next, the raw signals (without denoising)
of putative CPs are shown in color mixing mode.

For comparison, the raw signal of ATP is shown
(no denoising)

Pro Treh

GPE Treh

ATP

Legend

Pro Gln

Pro Asn

### 0256_SDA-1A-08

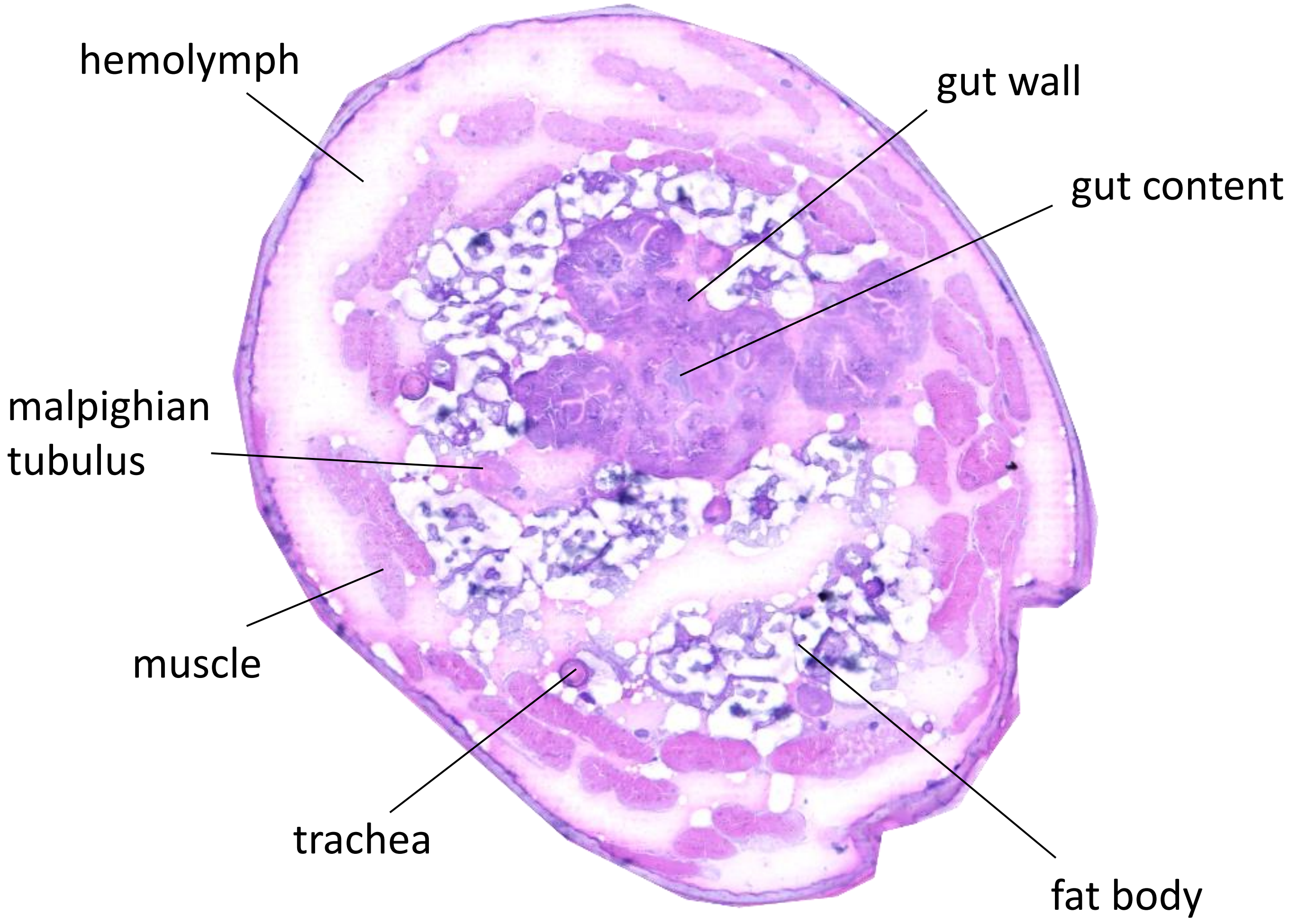


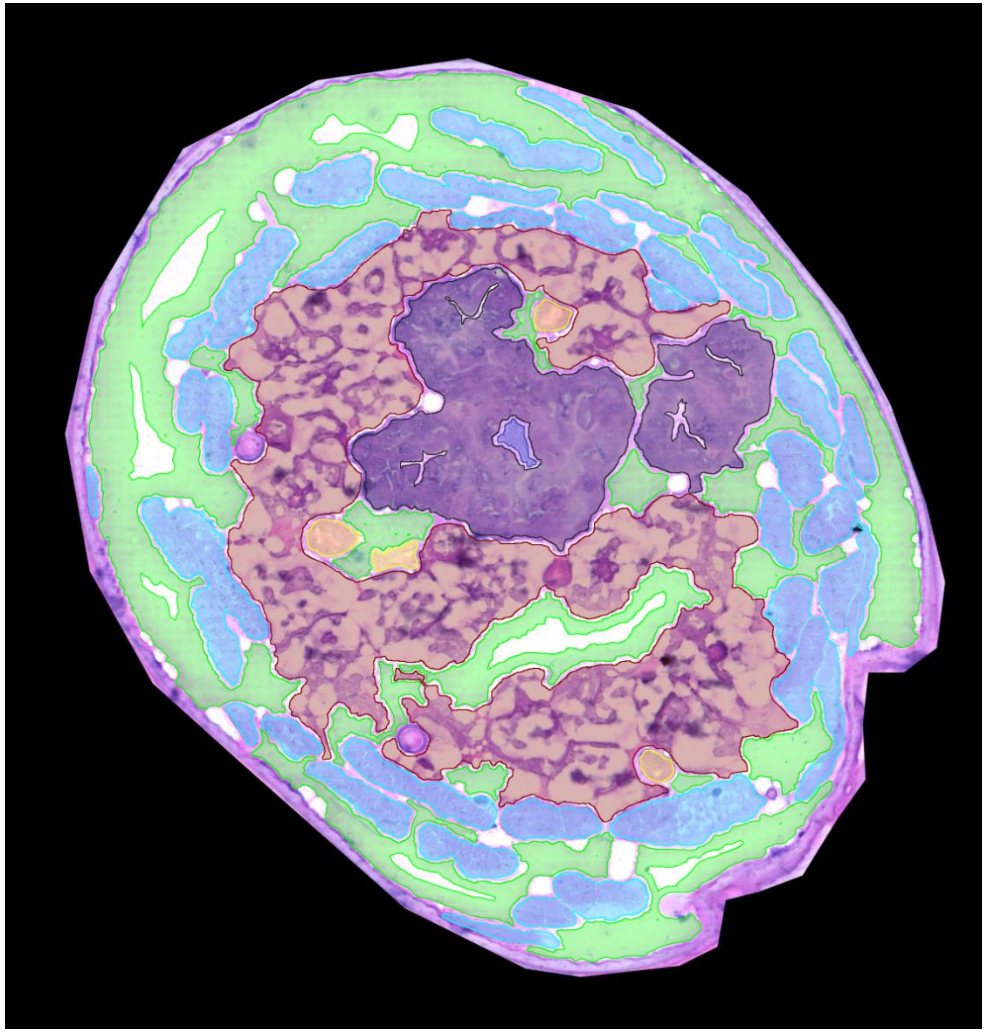

0256_SDA-1A-08

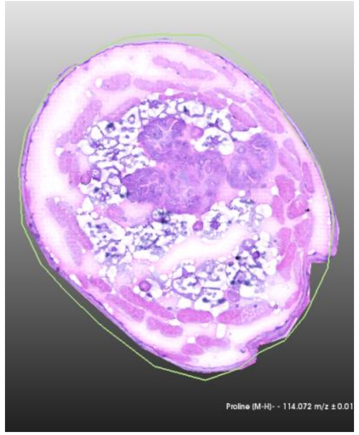

**H&E stain**

**Size bar** 200 µm

**Color scale**

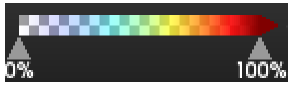


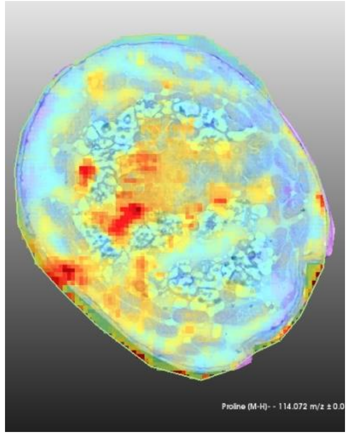

**Proline** [M-H]-
m/z 114,072 ± 0,017Da

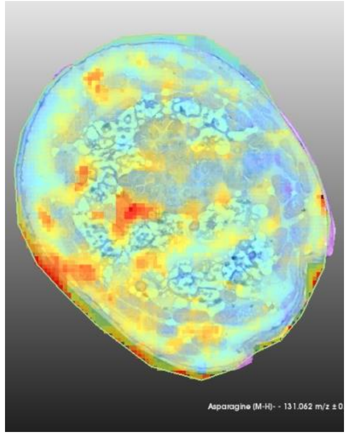

**Asparagine** [M-H]-
m/z 131,062 ± 0,02Da

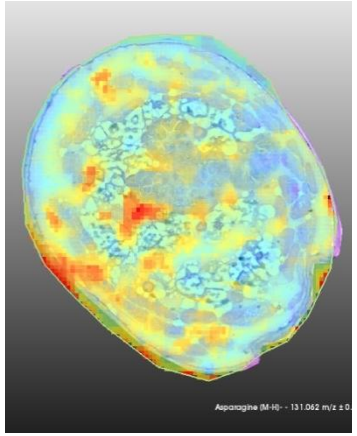

**Glutamine** [M-H]-
m/z 145,078 ± 0,022Da

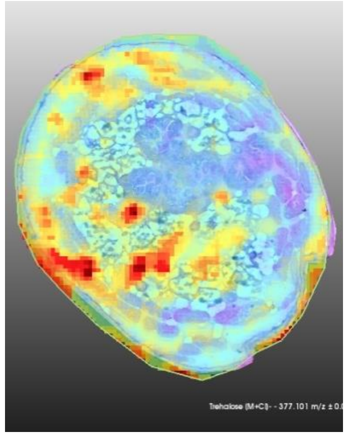

**Trehalose** [M+Cl]-
m/z 377,101 ± 0,057Da

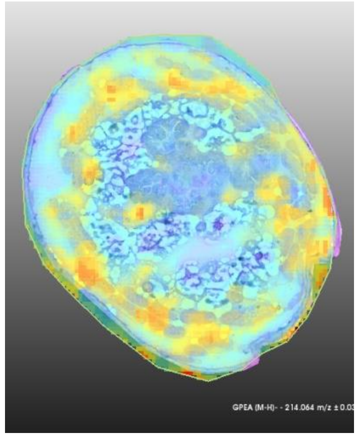

**GPEtn** [M-H]-
m/z 214,064 ± 0,032Da

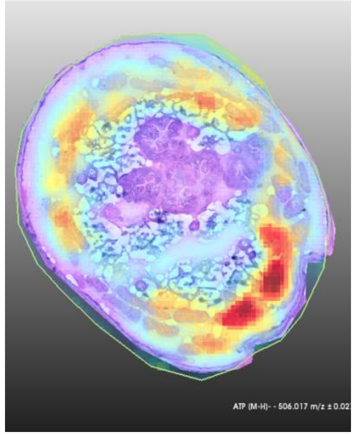

**ATP**[M-H]-
m/z 506,017 ± 0,027Da

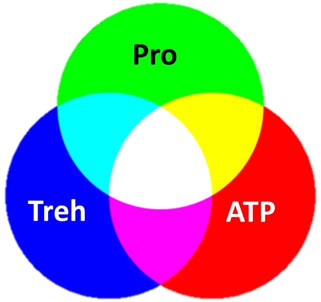


**Color mixing**

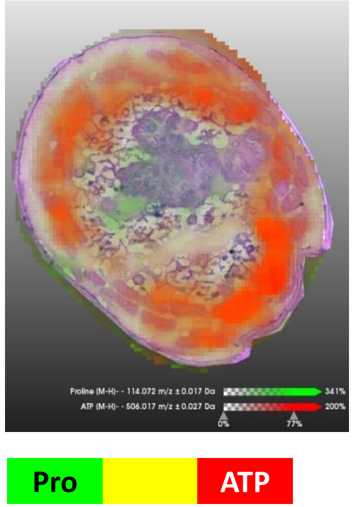


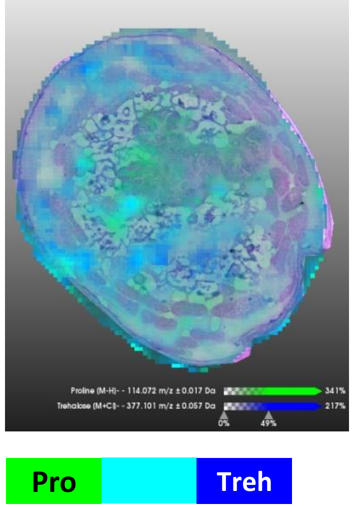


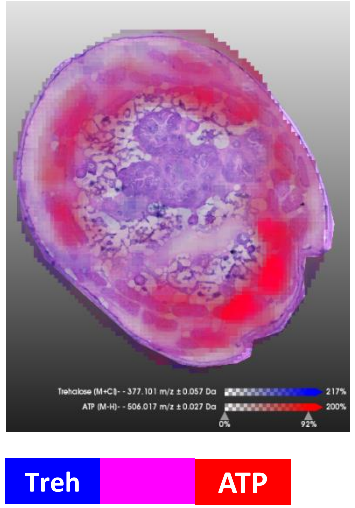

0256_SDA-1A-08

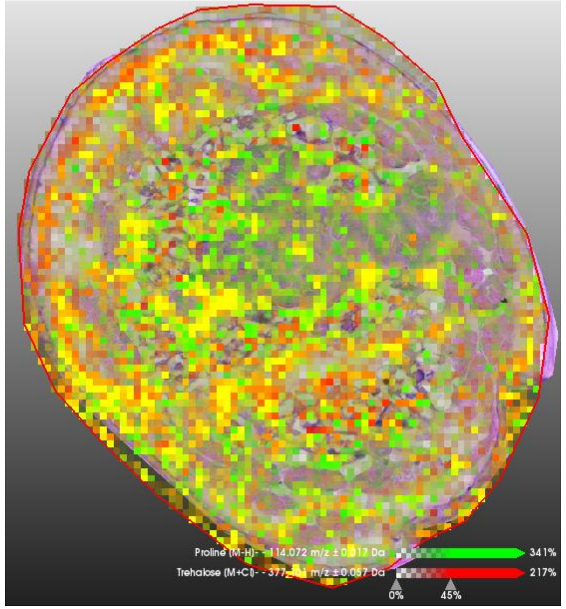


Pro | Treh

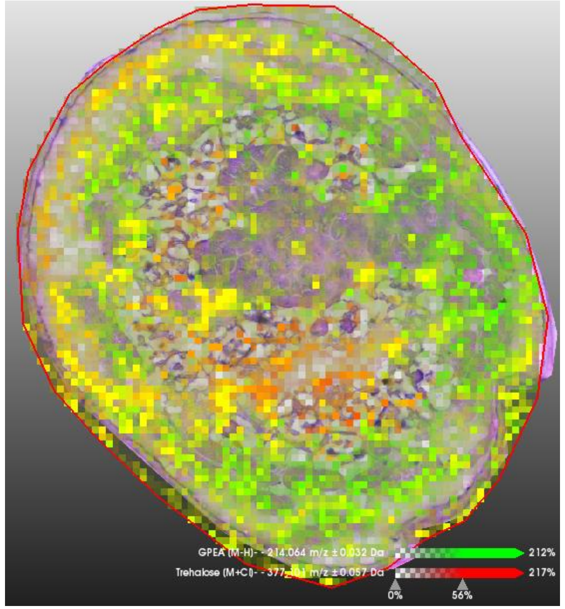


GPE | Treh

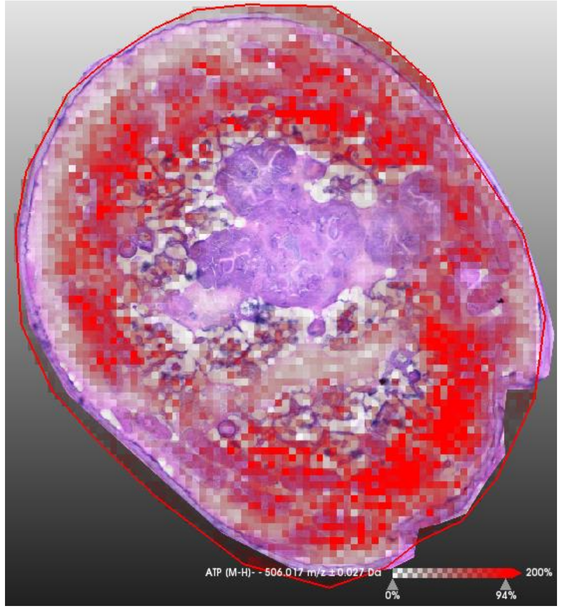


ATP

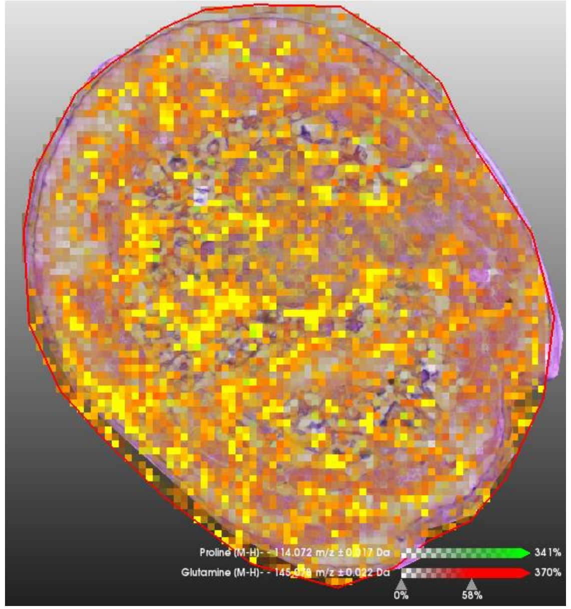


Pro | Gln

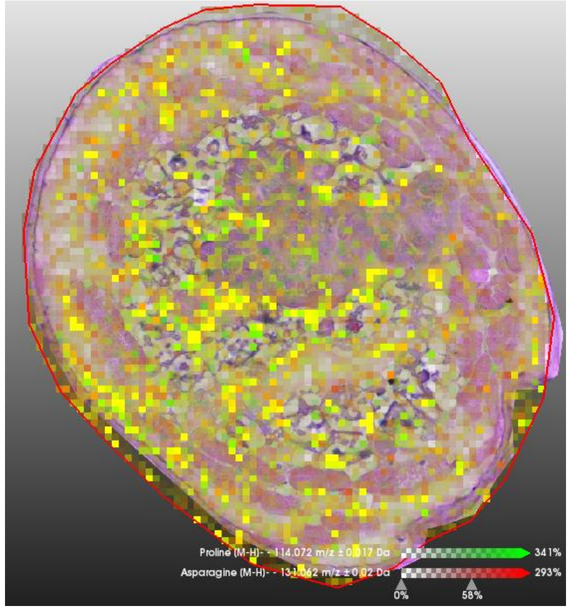


Pro | Asn

### 0256_SDA-1A-10

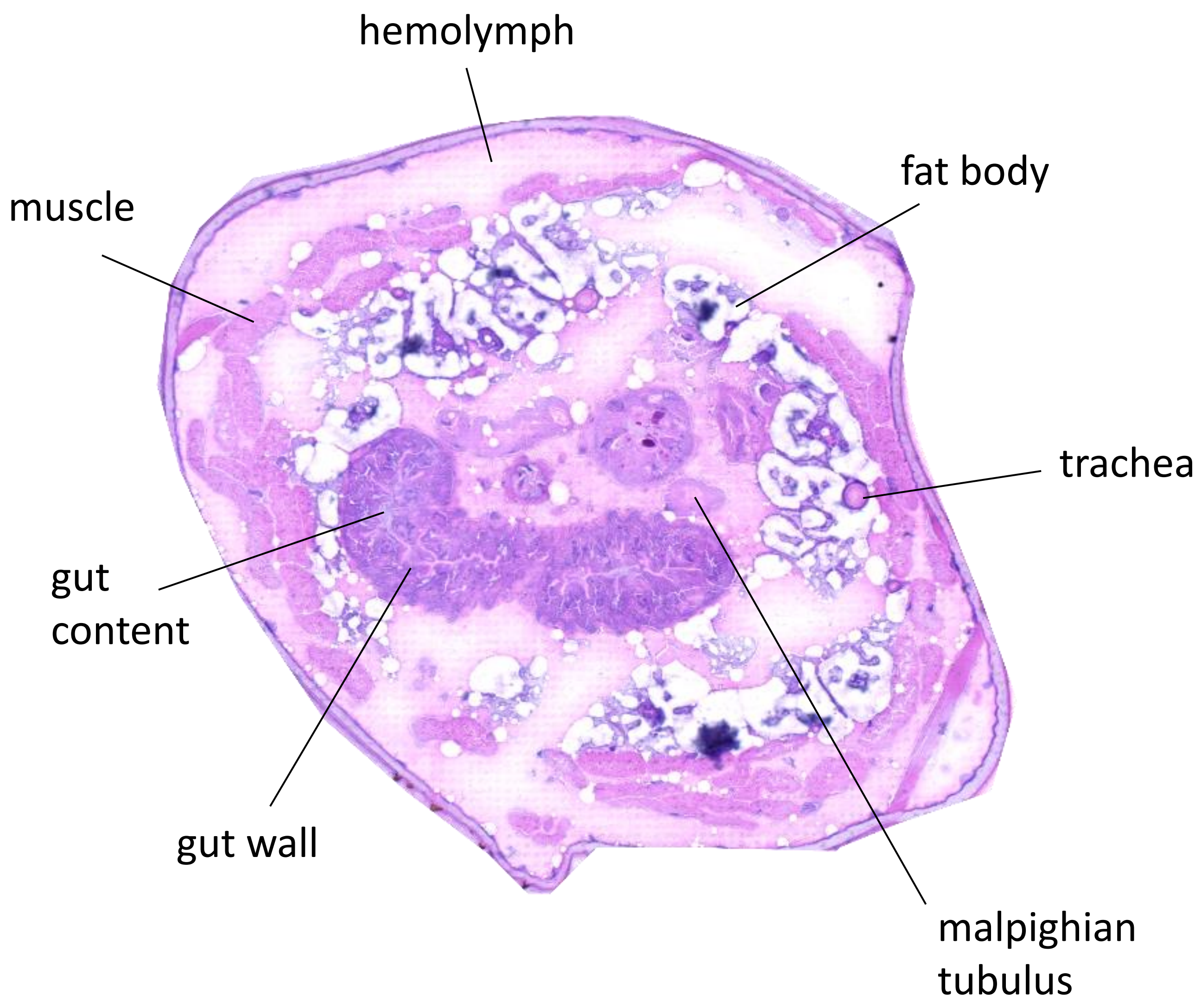


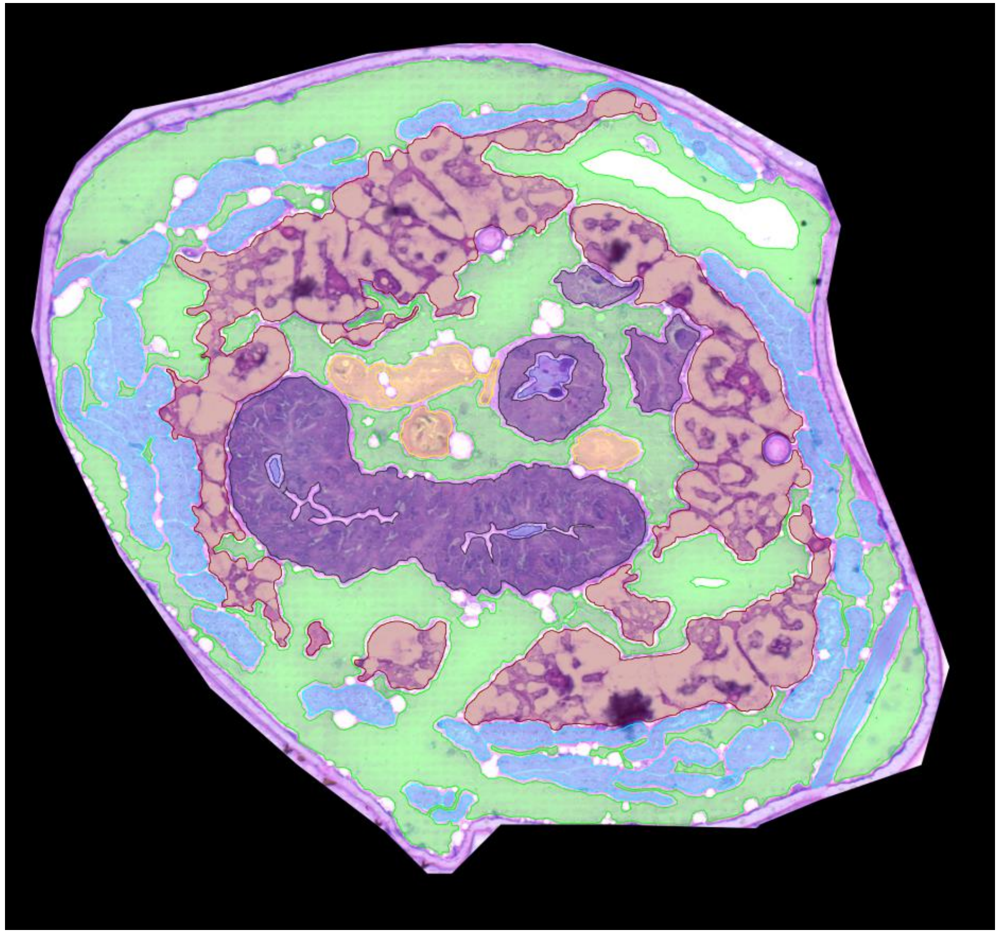

0256_SDA-1A-10

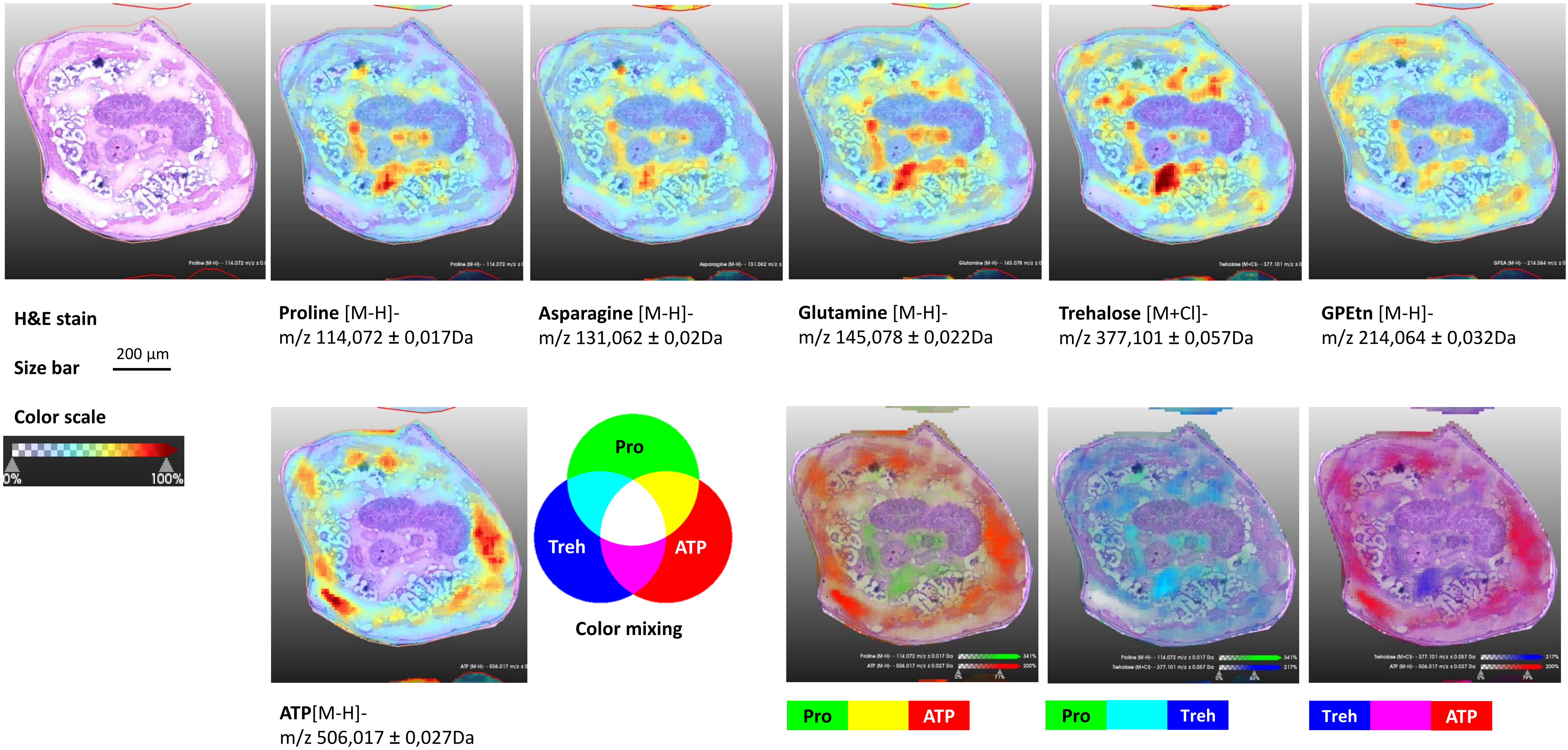

0256_SDA-1A-10

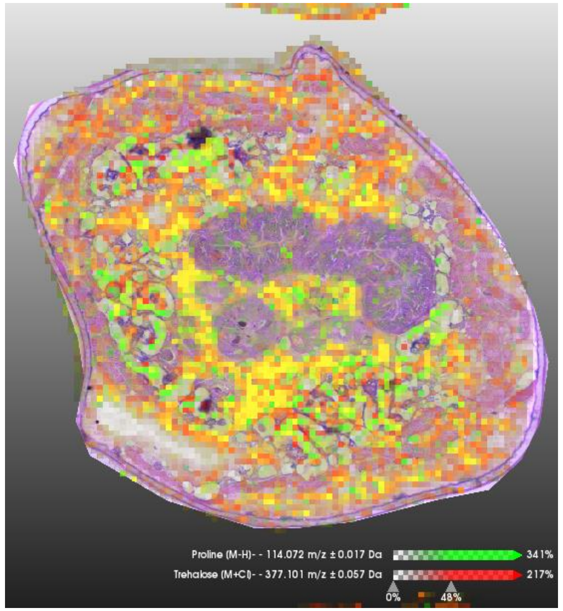


Pro Treh

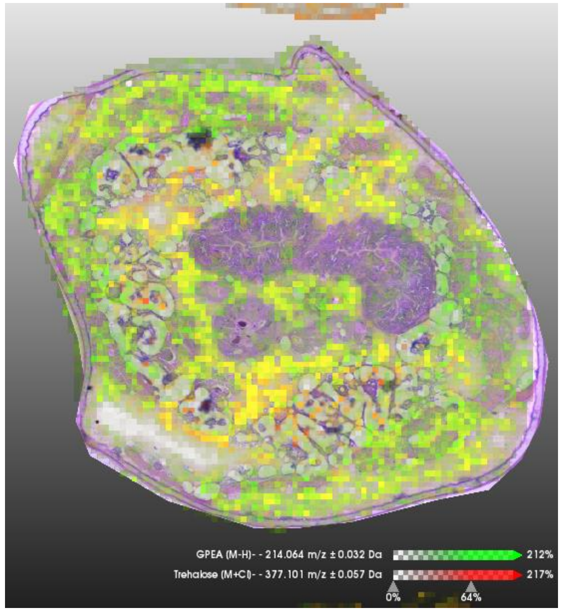


GPE Treh

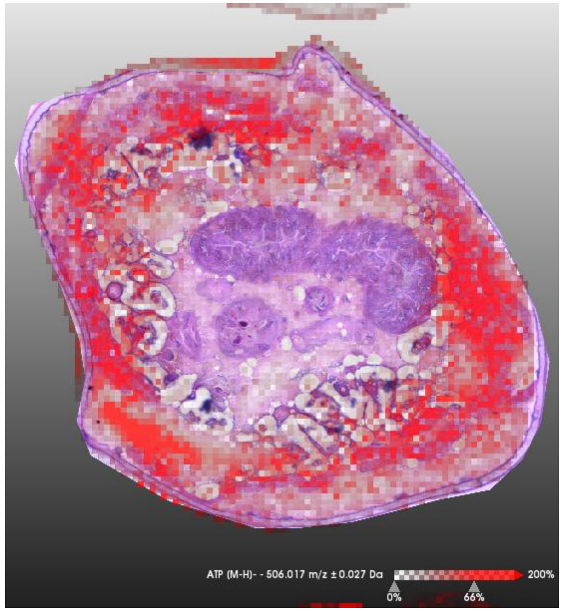


ATP

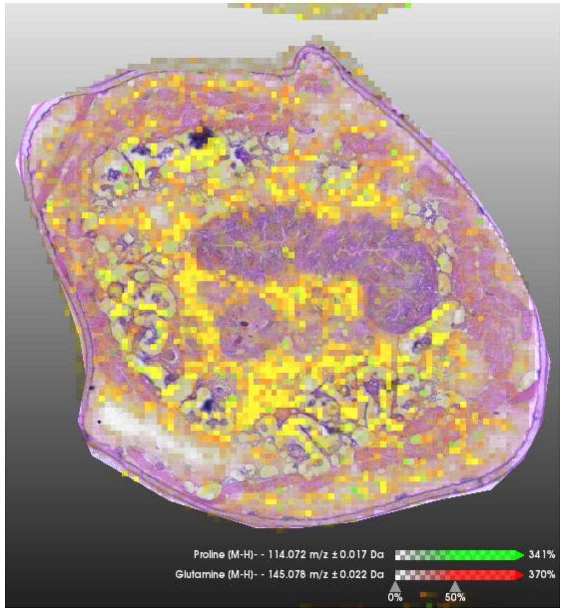


Pro Gln

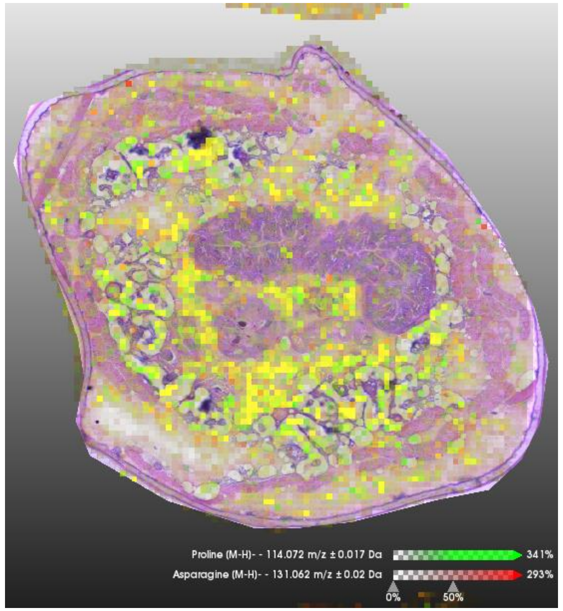


Pro Asn

### 0256_SDA-1A-03

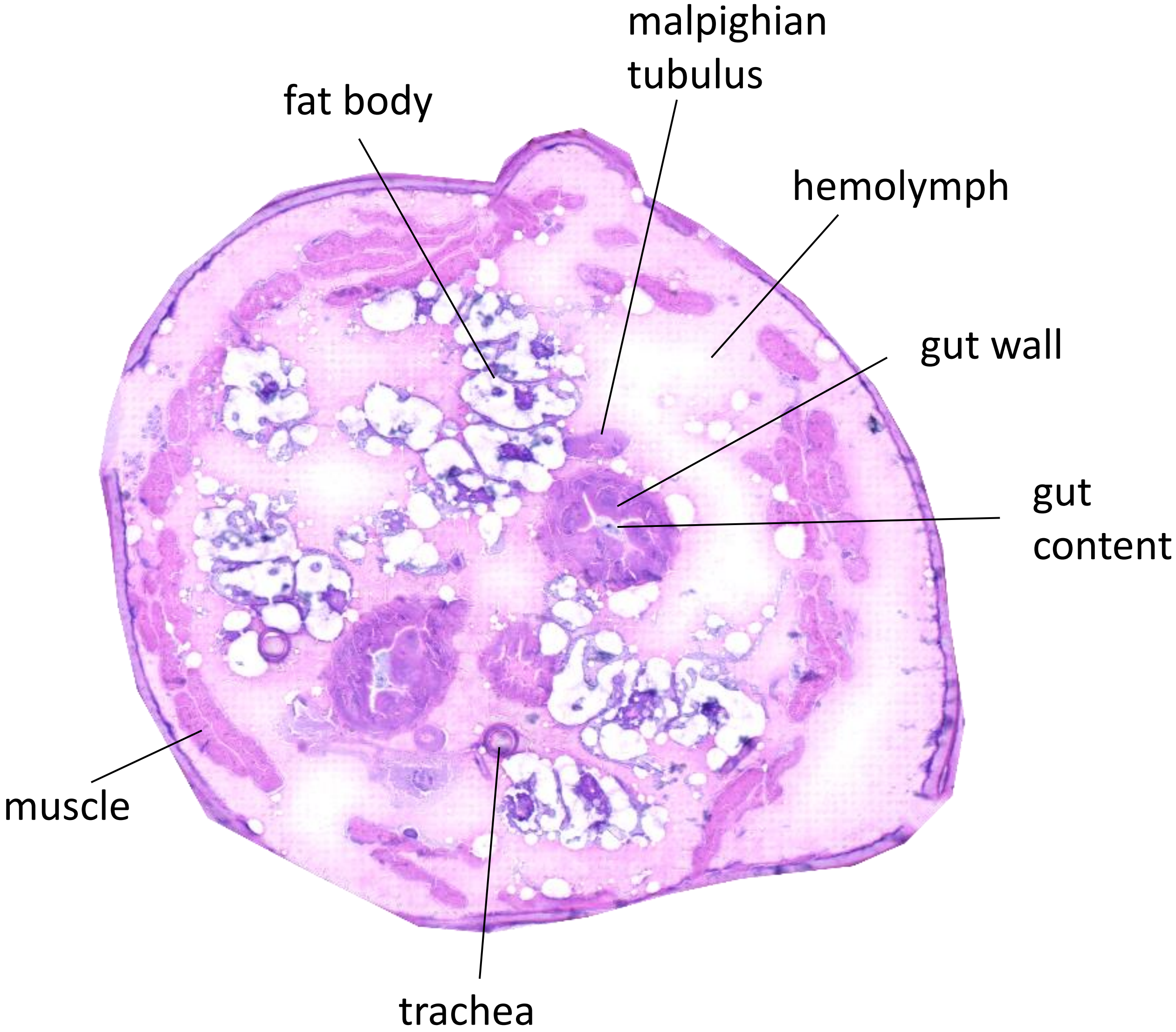


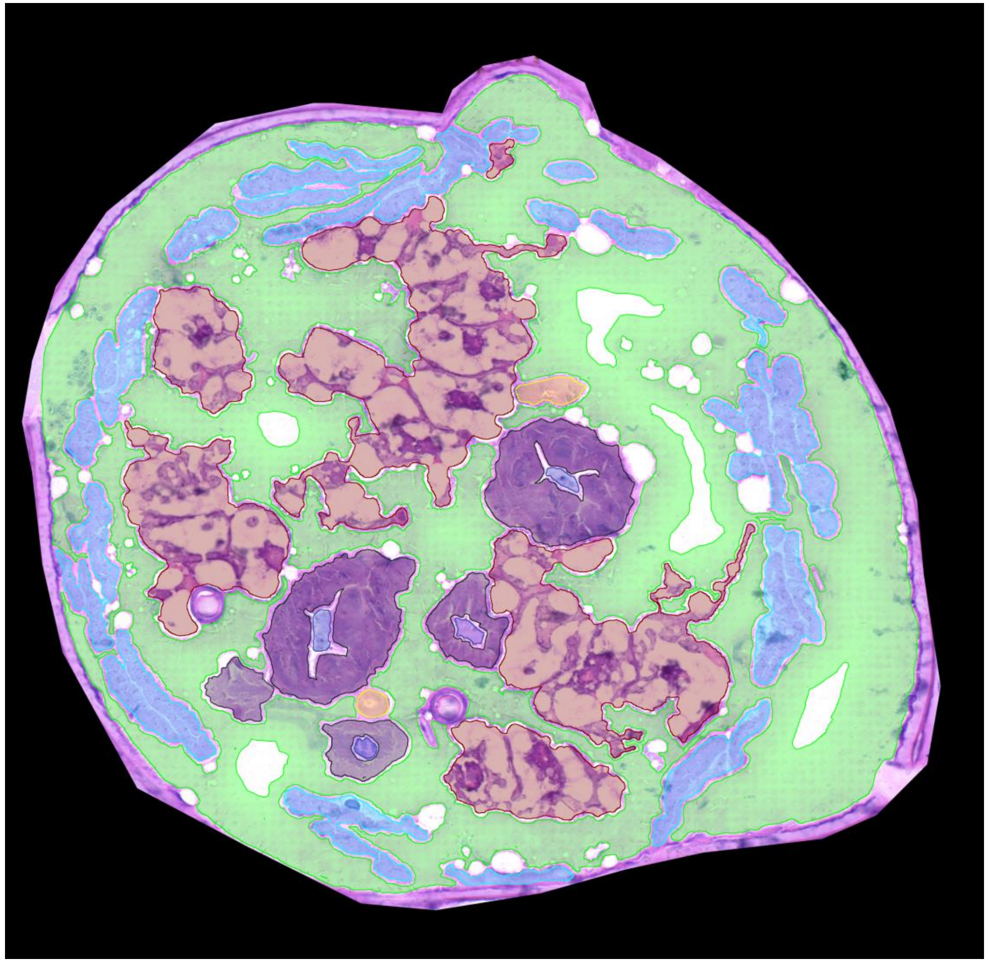

0256_SDA-1A-03

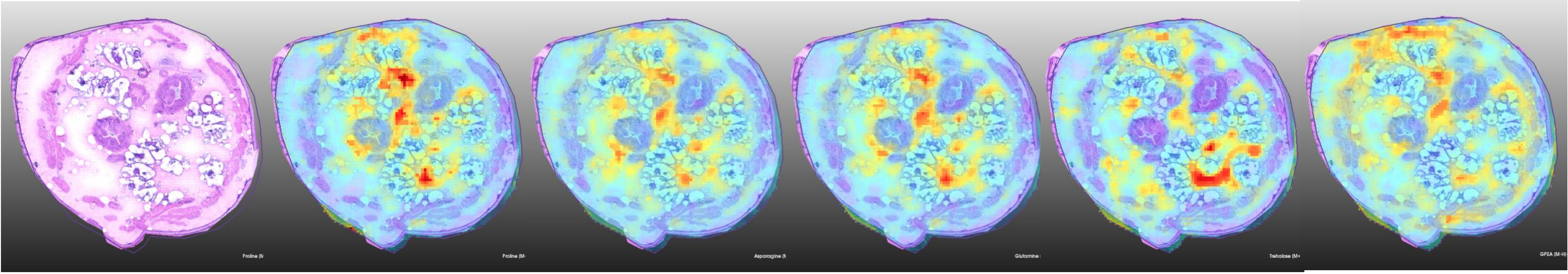


**H&E stain** | **Proline** [M-H]- m/z 114,072 ± 0,017Da | **Asparagine** [M-H]- m/z 131,062 ± 0,02Da | **Glutamine** [M-H]- m/z 145,078 ± 0,022Da | **Trehalose** [M+Cl]- m/z 377,101 ± 0,057Da | **GPEtn** [M-H]- m/z 214,064 ± 0,032Da

**Size bar** 200 µm

**Color scale**

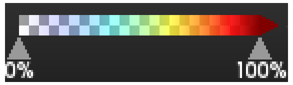


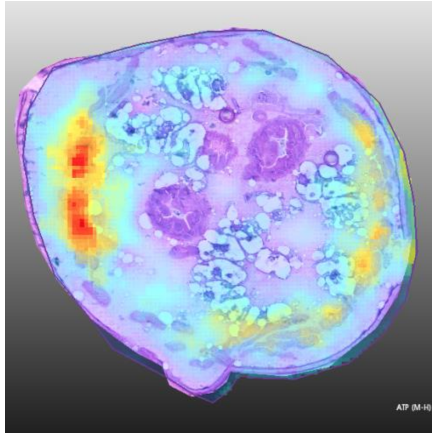


**ATP**[M-H]-
m/z 506,017 ± 0,027Da

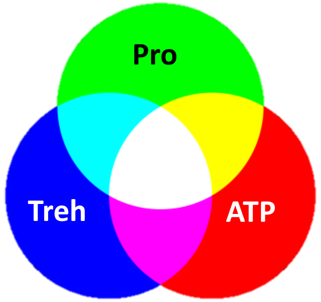


**Color mixing**

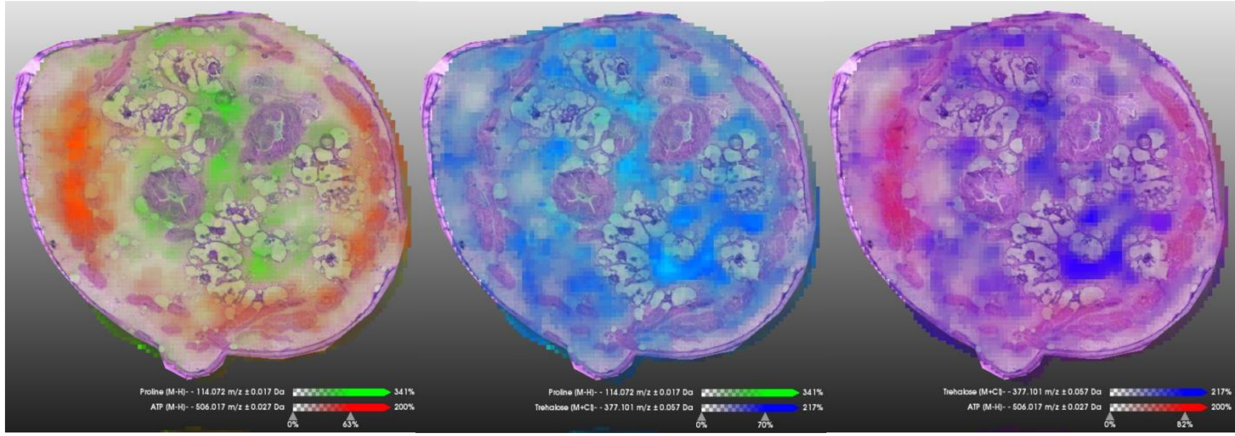

**Pro** **ATP** | **Pro** **Treh** | **Treh** **ATP**

0256_SDA-1A-03

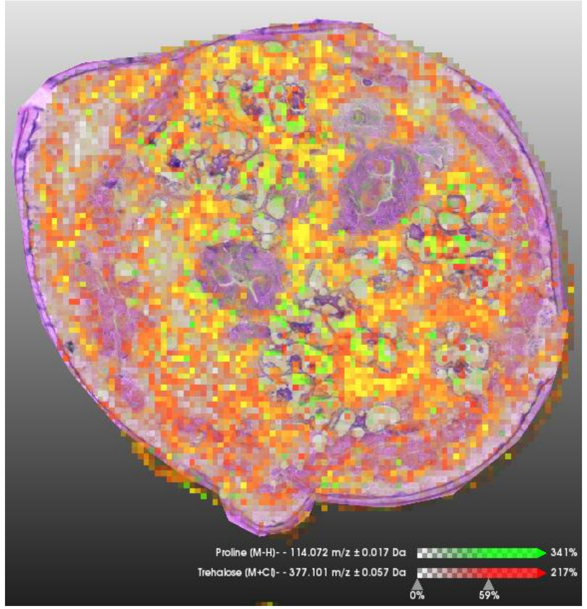


Pro Treh

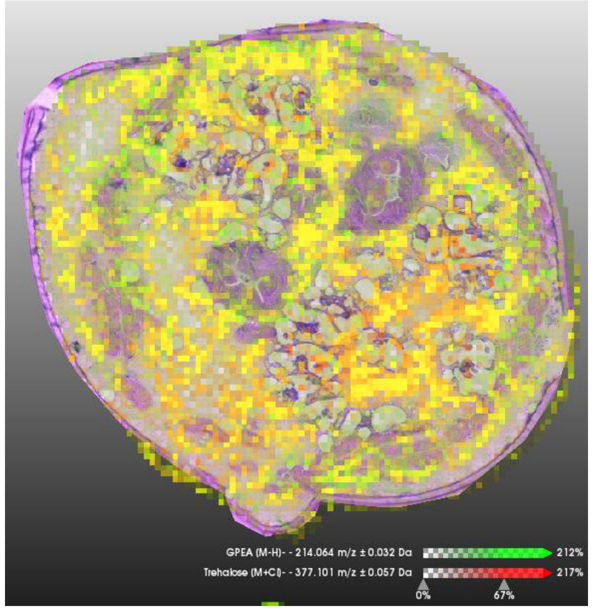


GPE Treh

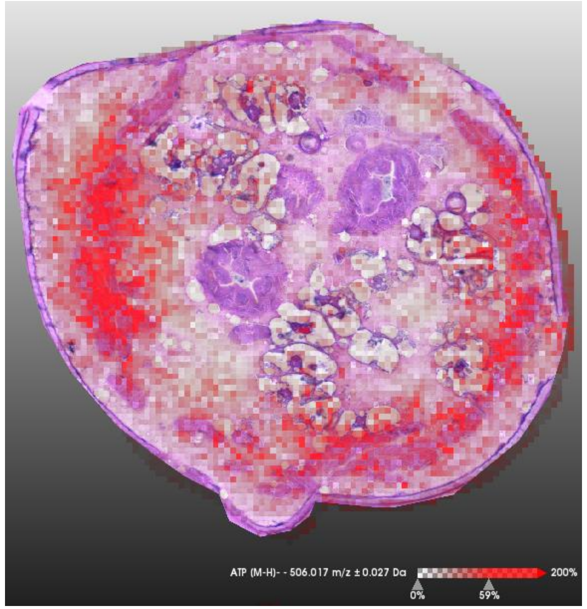


ATP

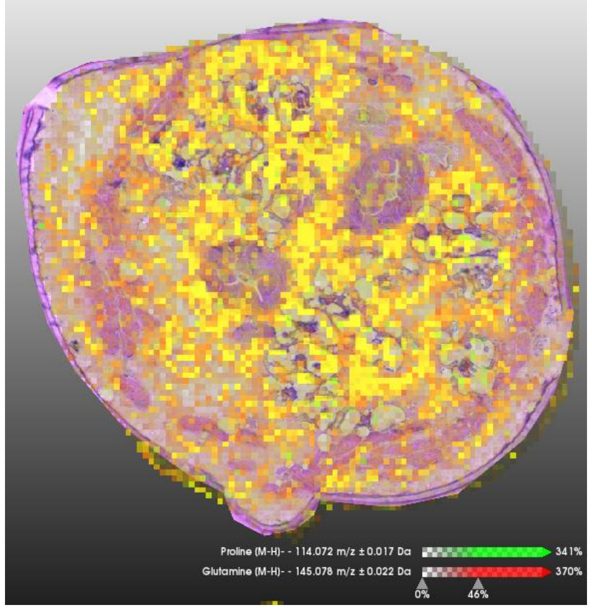


Pro Gln

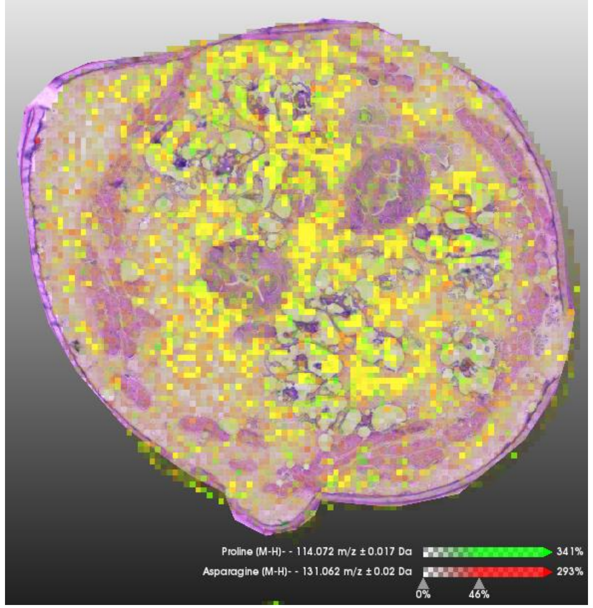


Pro Asn

### 0263_SDA-2B-05

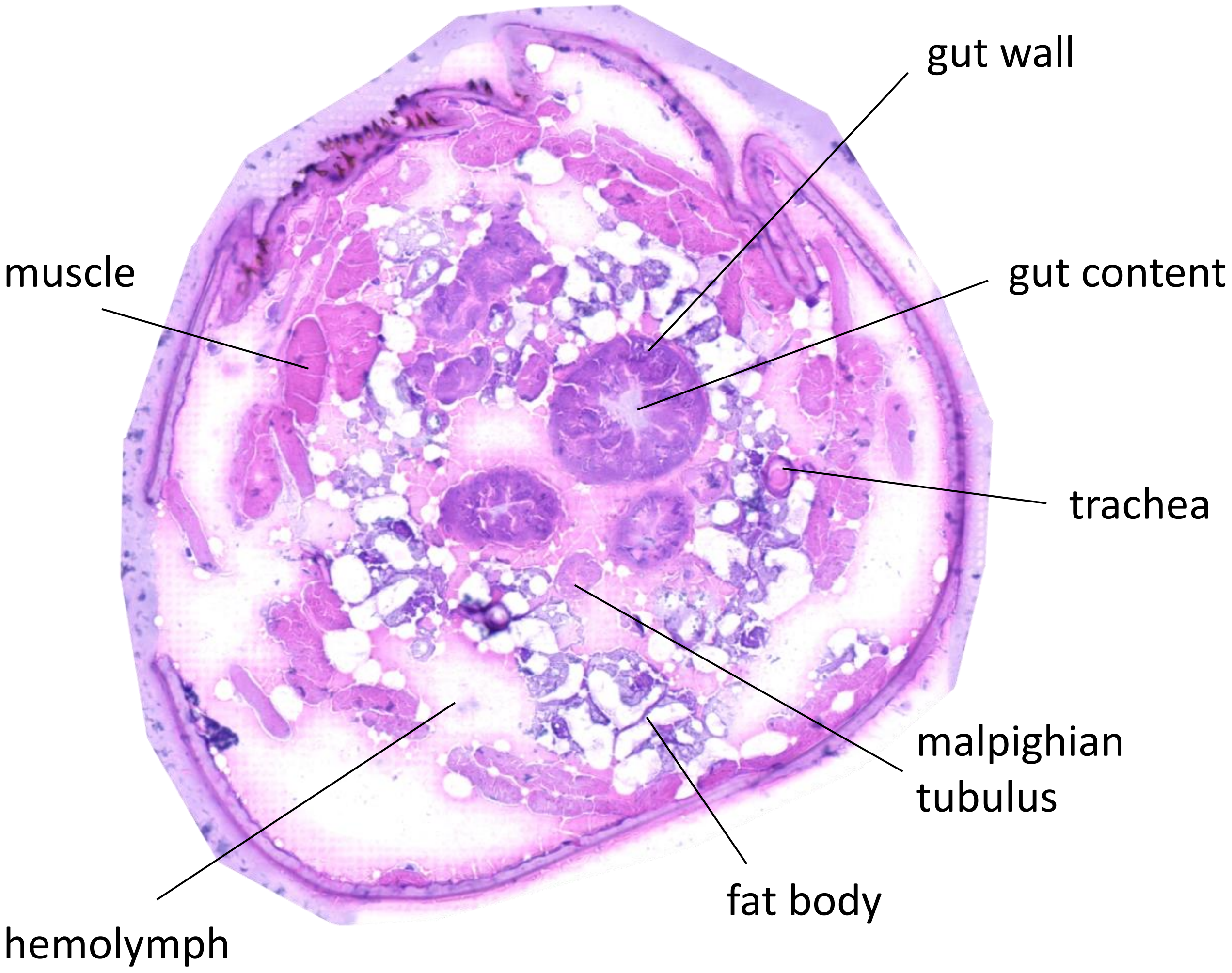


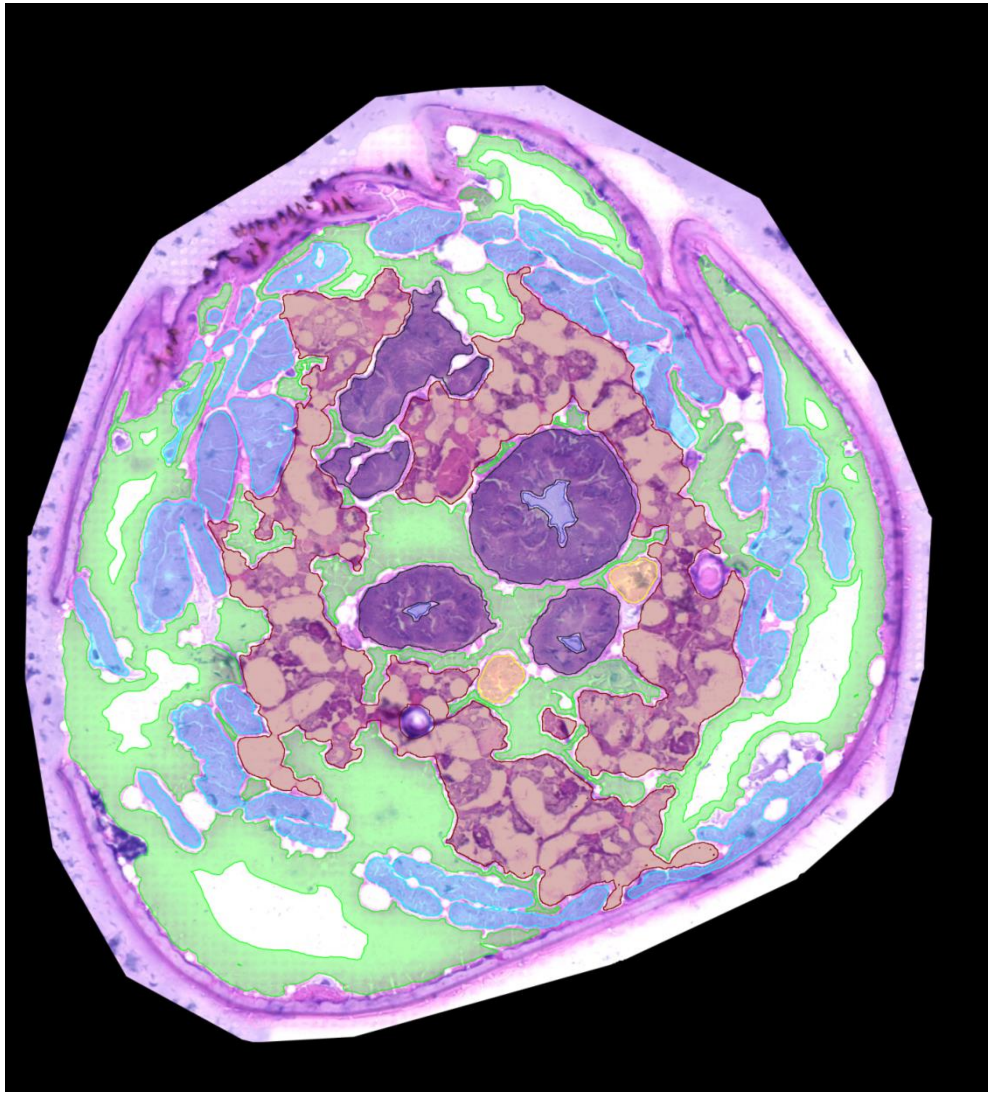

0263_SDA-2B-05

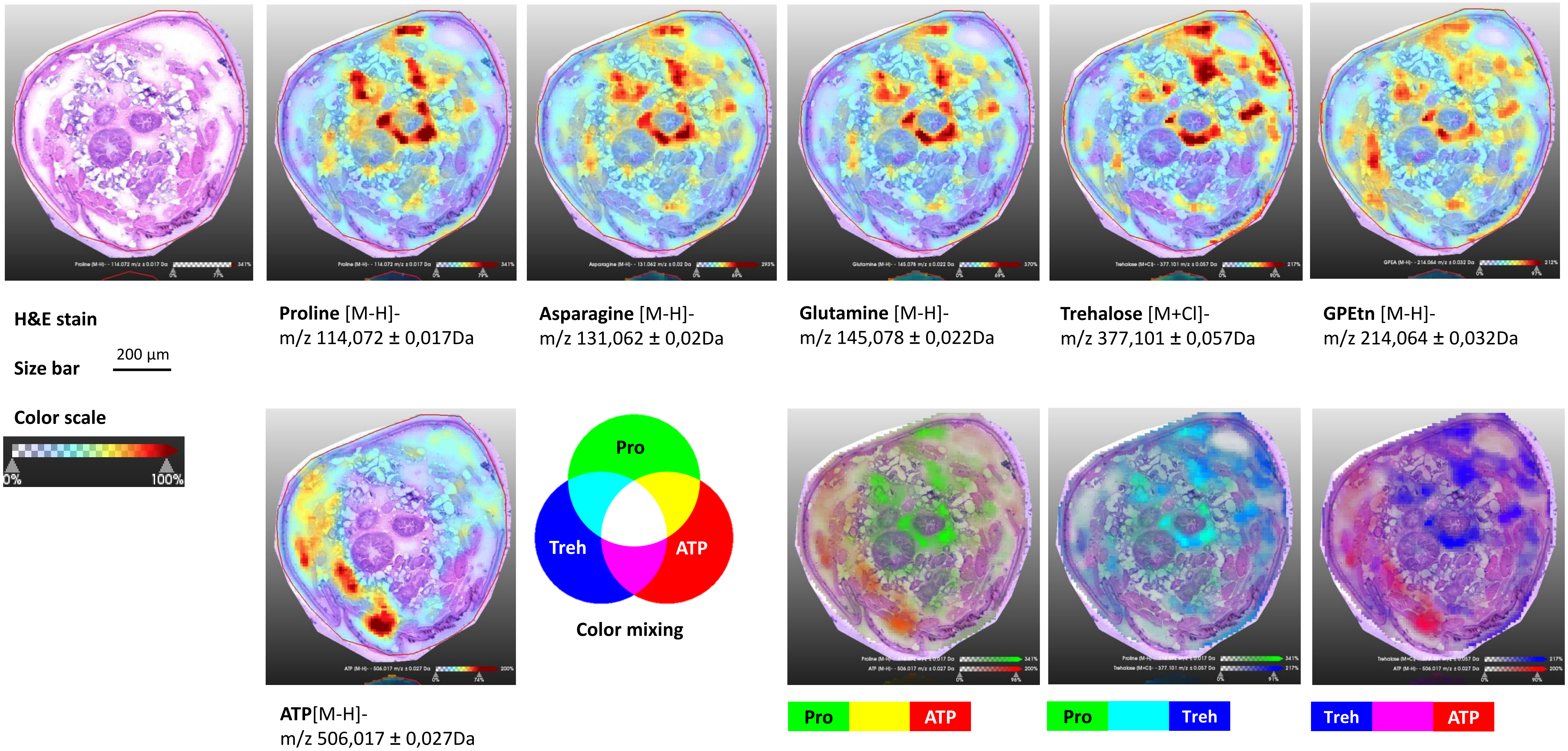

0263_SDA-2B-05

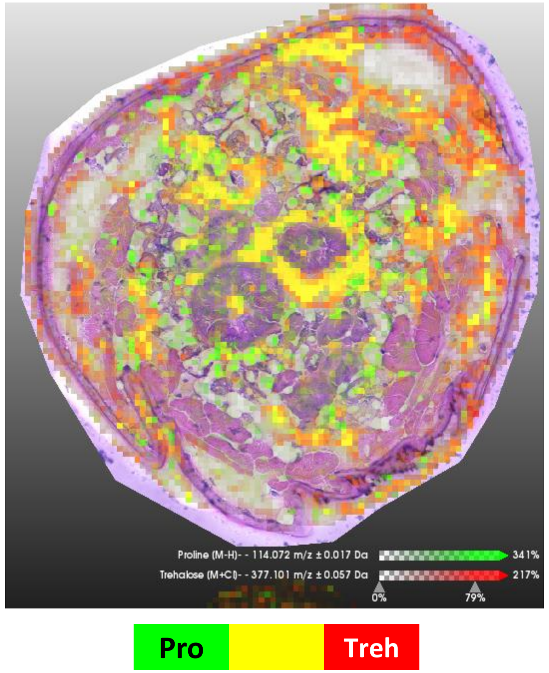


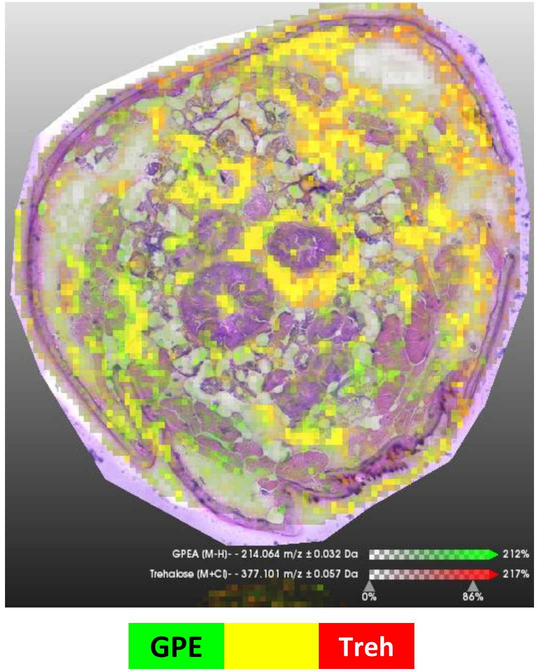


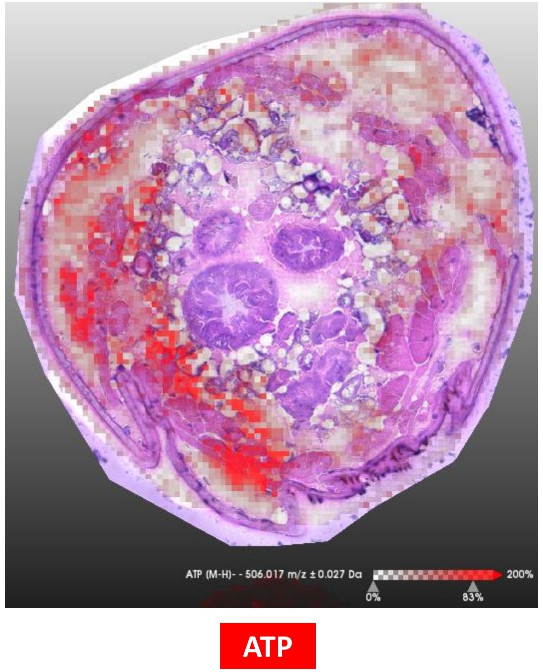


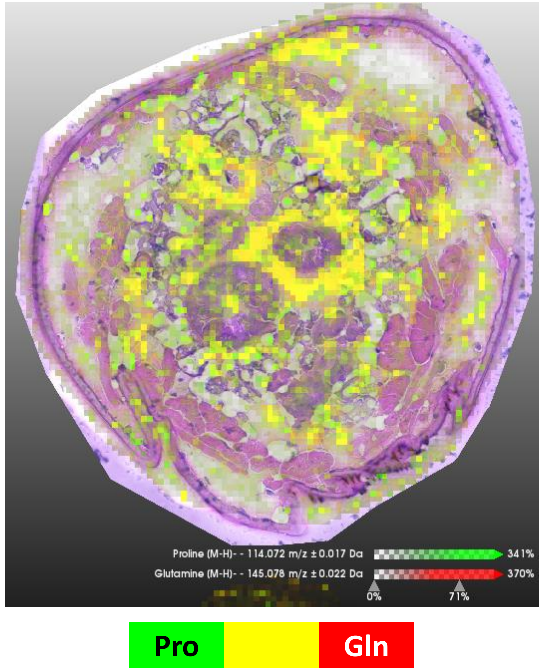


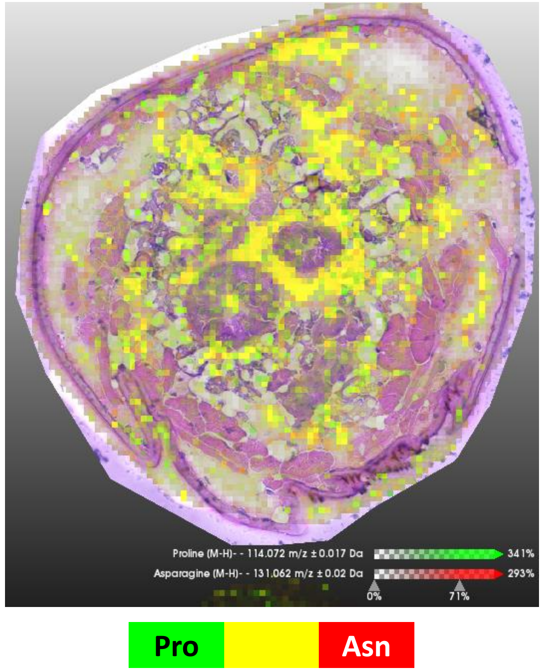

### 0268_SDA-3B-03

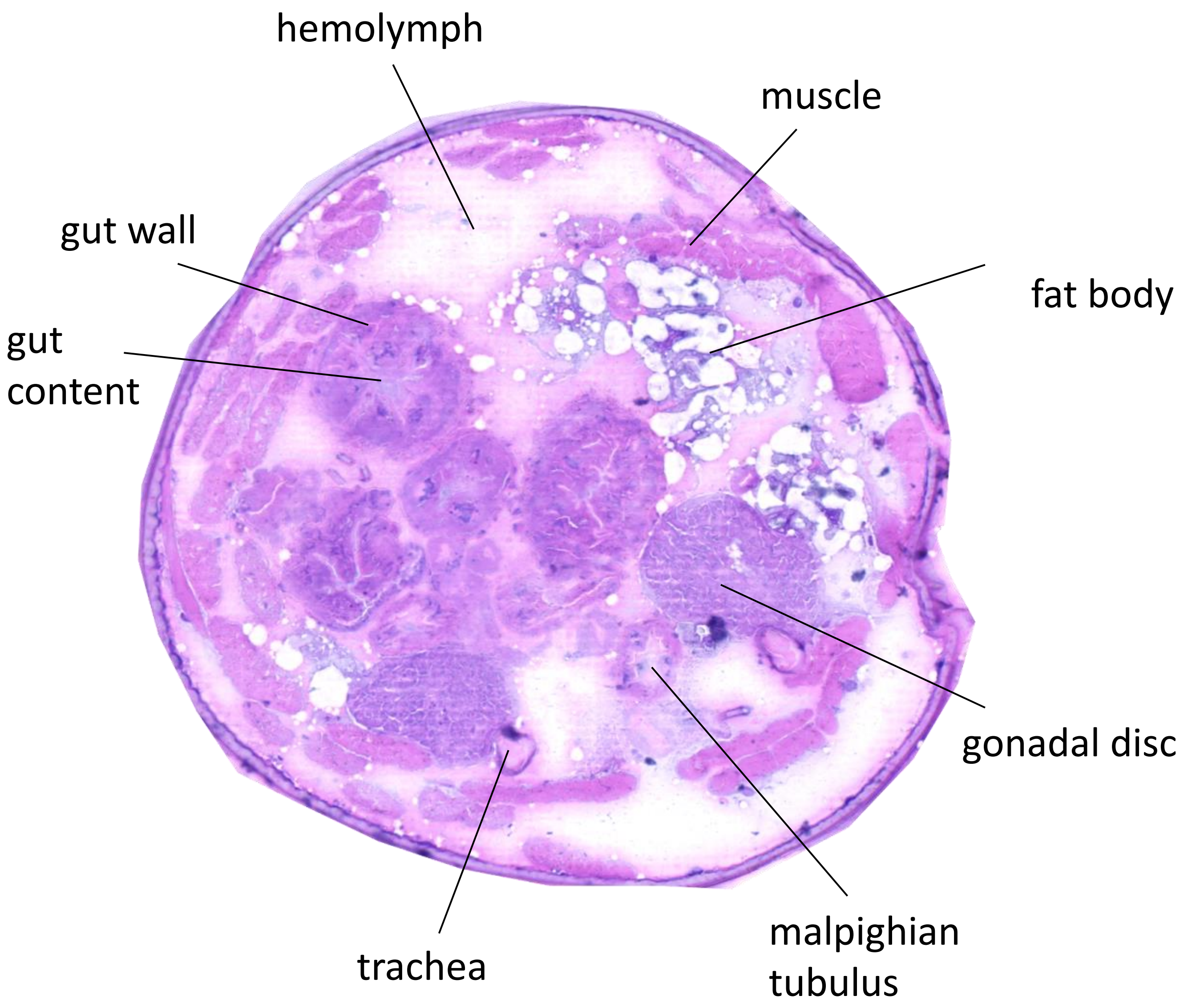


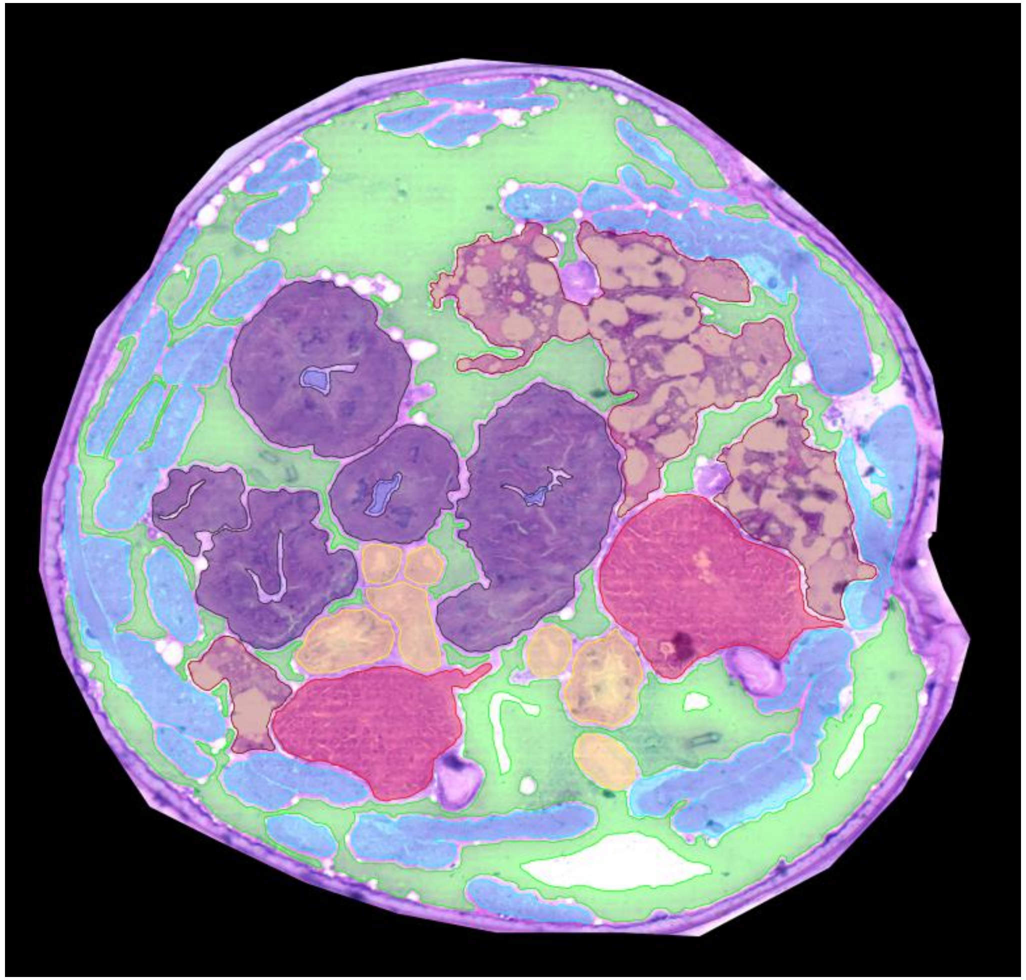

0268_SDA-3B-03

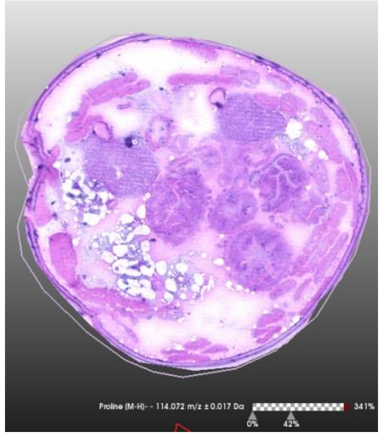

H&E stain

Size bar 200 µm

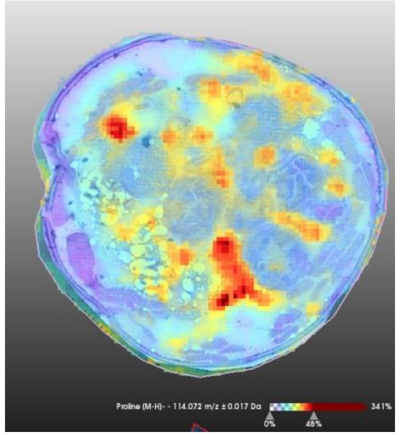

**Proline** [M-H]-
m/z 114,072 ± 0,017Da

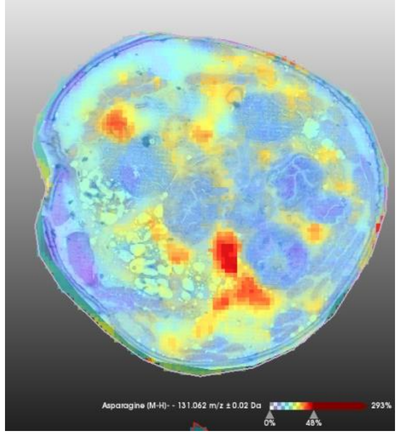

**Asparagine** [M-H]-
m/z 131,062 ± 0,02Da

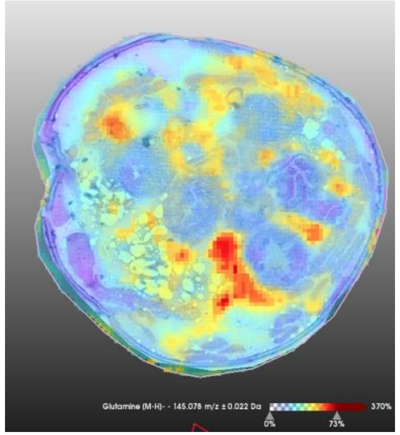

**Glutamine** [M-H]-
m/z 145,078 ± 0,022Da

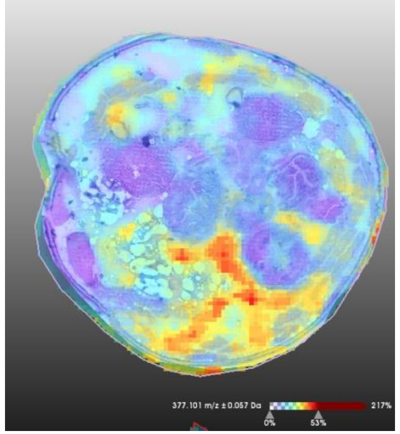

**Trehalose** [M+Cl]-
m/z 377,101 ± 0,057Da

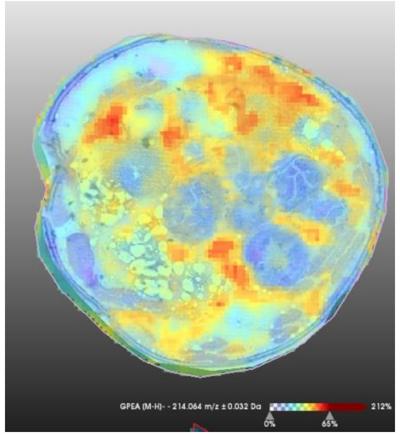

**GPEtn** [M-H]-
m/z 214,064 ± 0,032Da

Color scale

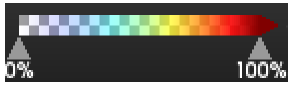


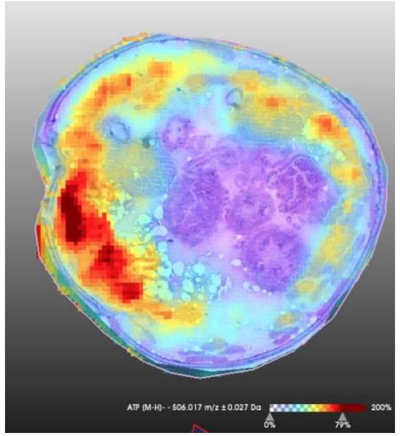

**ATP**[M-H]-
m/z 506,017 ± 0,027Da

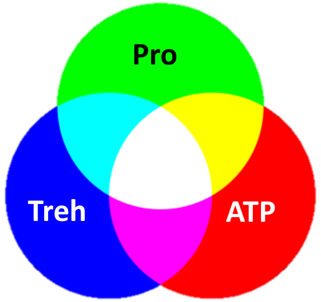


Color mixing

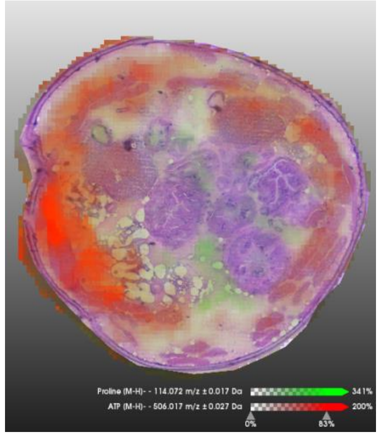

Pro ATP

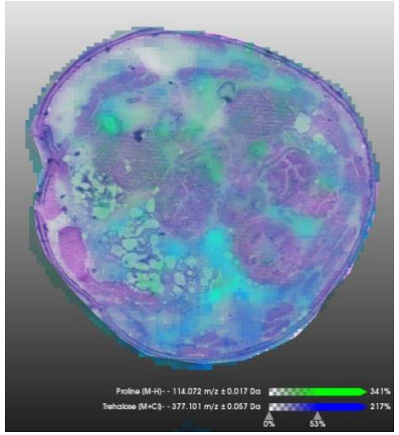

Pro Treh

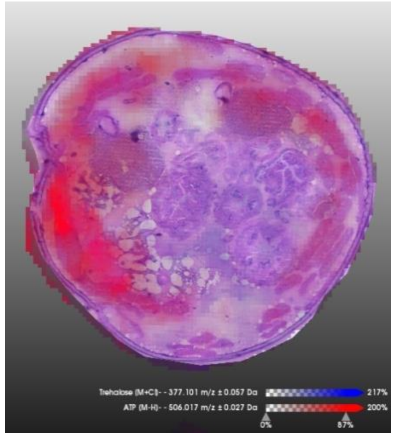

Treh ATP

0268_SDA-3B-03

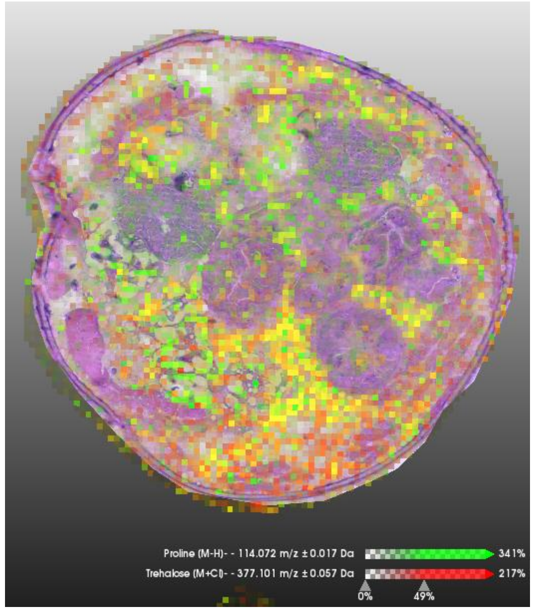


Pro | Treh

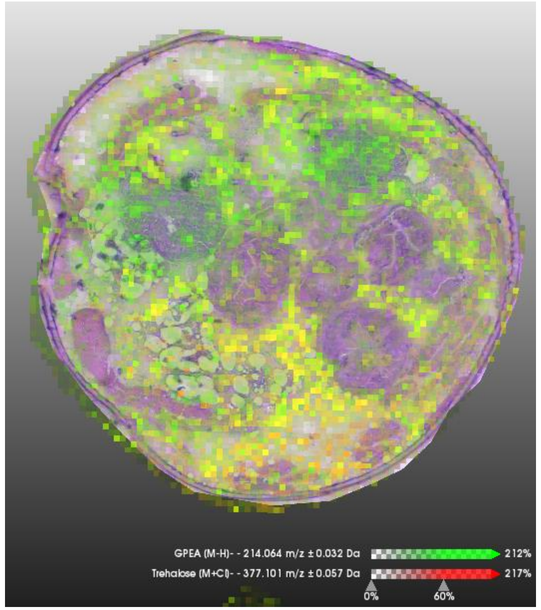


GPE | Treh

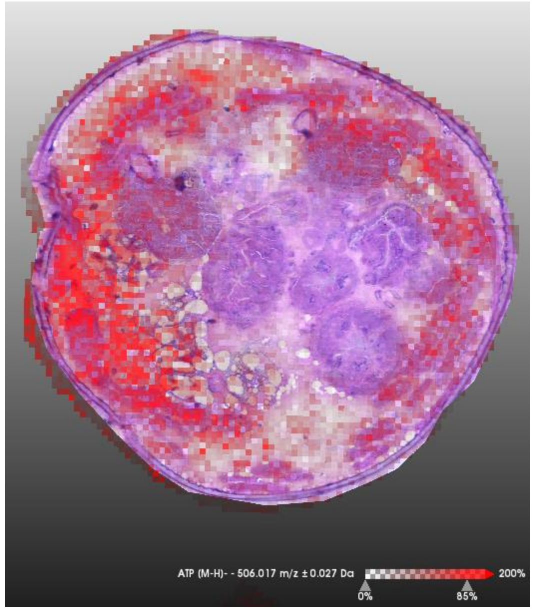


ATP

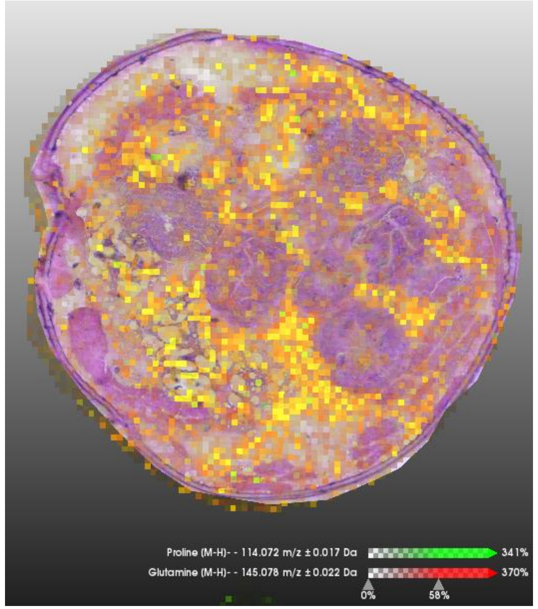


Pro | Gln

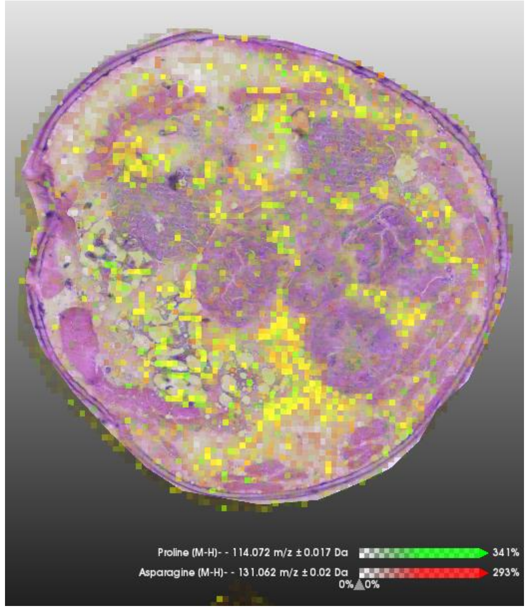


Pro | Asn

*Chymomyza costata*, 3rd instar larvae, 3-week-old

**phenotypic/acclimation variant SDA-frozen**

after slow freezing to -30°C

transversal sections through the middle part of larva

### Legend

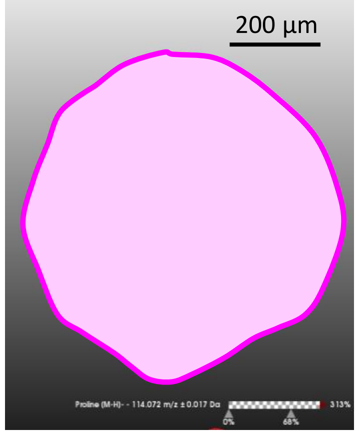


**H&E stain**

**Size bar** 200 µm

**Color scale**

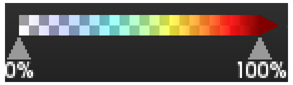


For each section, the signal intensities of five putative CPs (proline, asparagine, glutamine, trehalose, and GPE) plus ATP are shown using arbitrary color scale overlaid over H&E stained transversal section. The same scale bar (200 µm applies to all sections). Medium denoising (SCiLS tool) of signal intensities among neighboring pixels of 10x10 µm was applied.

**Proline** [M-H]- m/z 114,072 ± 0,017Da

**Asparagine** [M-H]- m/z 131,062 ± 0,02Da

**Glutamine** [M-H]- m/z 145,078 ± 0,022Da

**Trehalose** [M+Cl]- m/z 377,101 ± 0,057Da

**GPE** [M-H]- m/z 214,064 ± 0,032Da

**ATP**[M-H]- m/z 506,017 ± 0,027Da

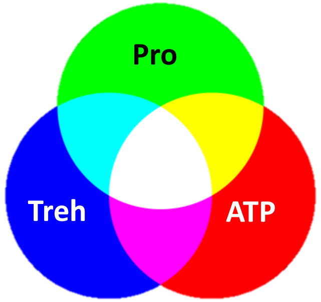


**Color mixing**

Next, signals of three select compounds (proline, trehalose, and ATP) are shown in color mixing mode.

**Pro** **ATP**

**Pro** **Treh**

**Treh** **ATP**

0258_LD-3A-06

Next, the raw signals (without denoising)
of putative CPs are shown in color mixing mode.

Pro Treh

GPE Treh

For comparison, the raw signal of ATP is shown
(no denoising)

ATP

Legend

Pro Gln

Pro Asn

### 0330B_SDA_A2_frozen-06

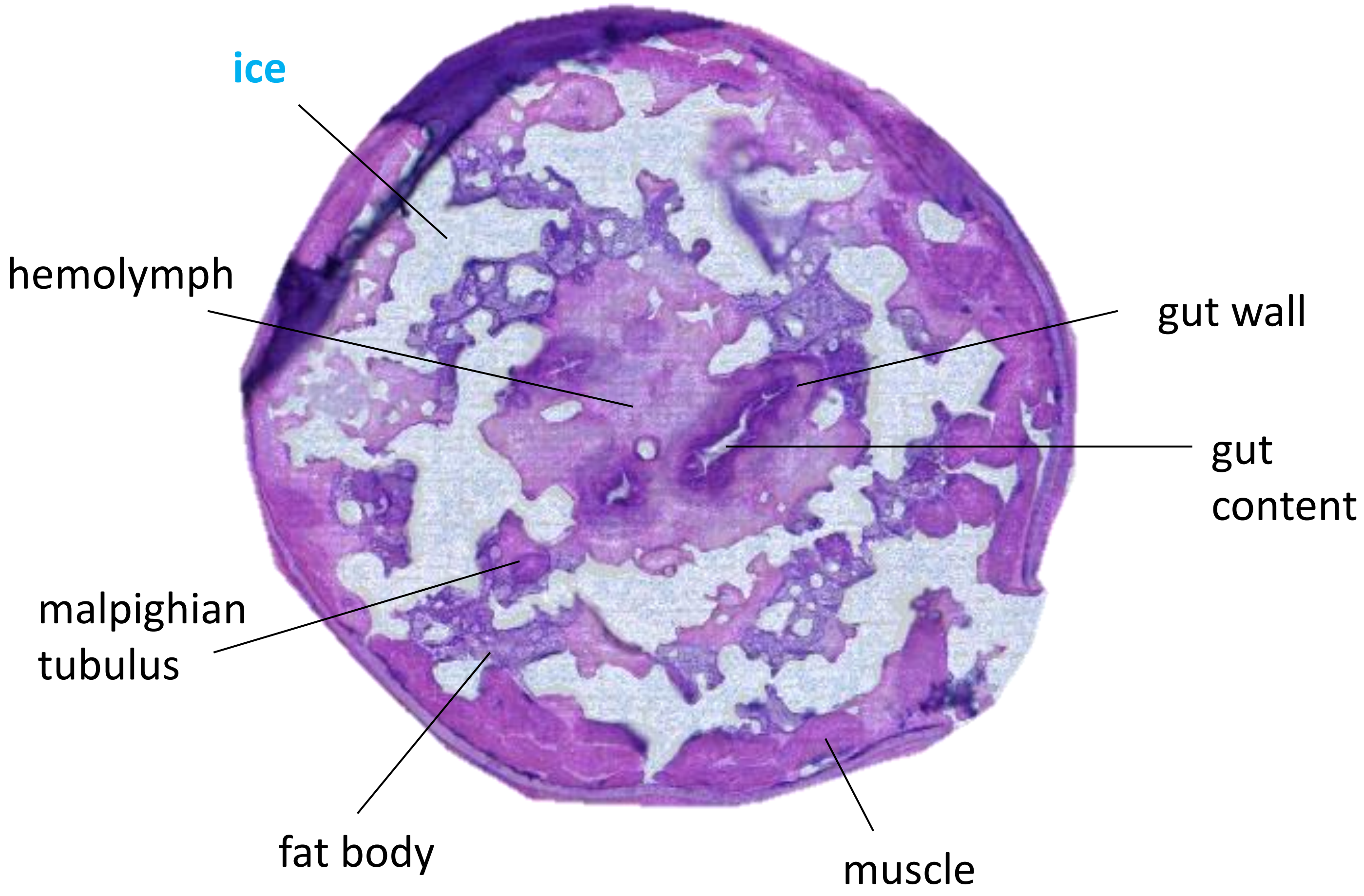


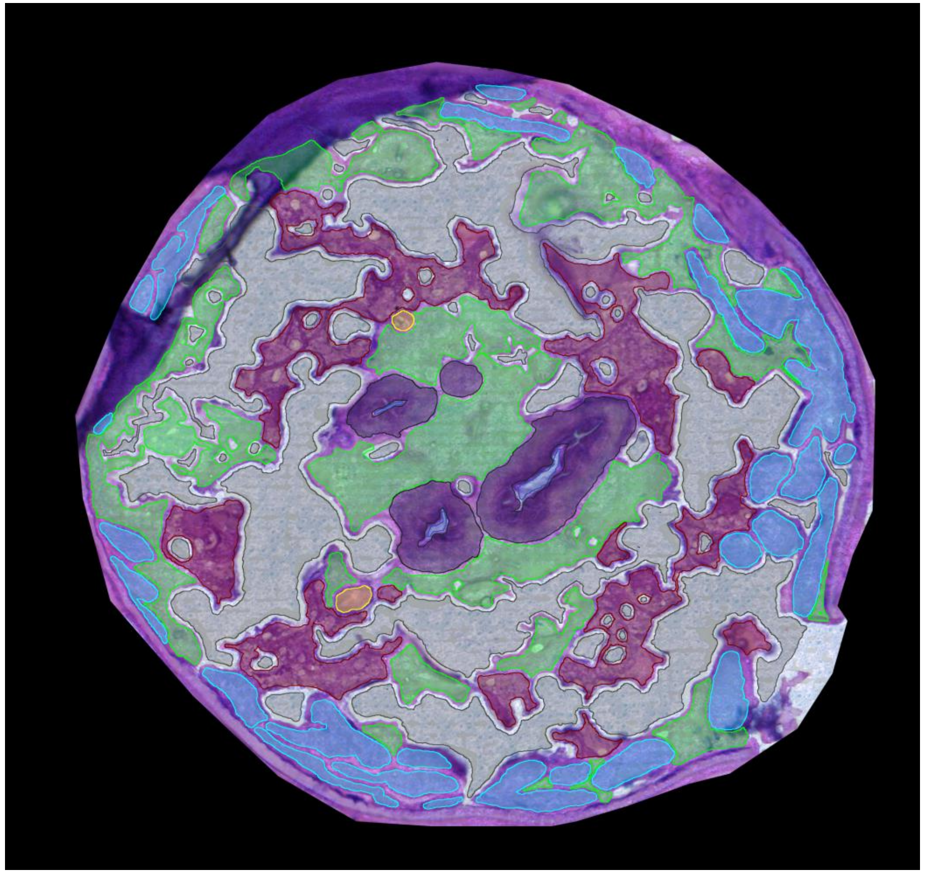

0330B_SDA_A2_Lyo-06

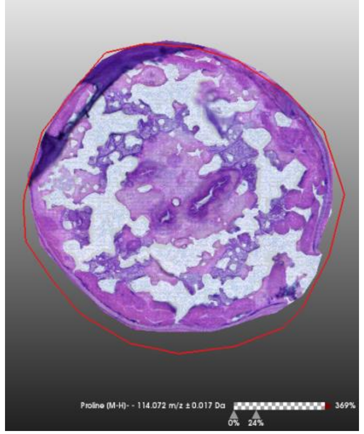

**H&E stain**

**Size bar** 200 µm

**Color scale**

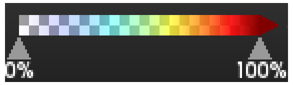


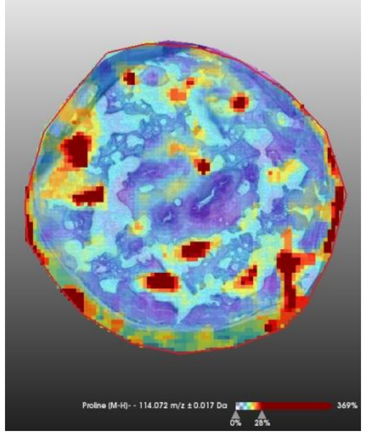

**Proline** [M-H]-
m/z 114,072 ± 0,017Da

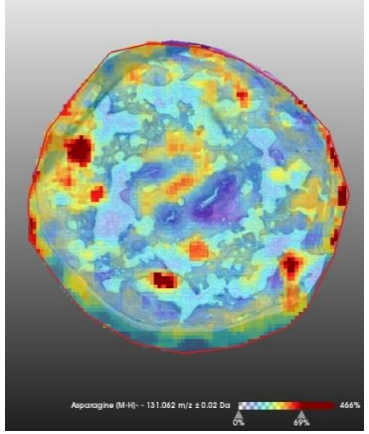

**Asparagine** [M-H]-
m/z 131,062 ± 0,02Da

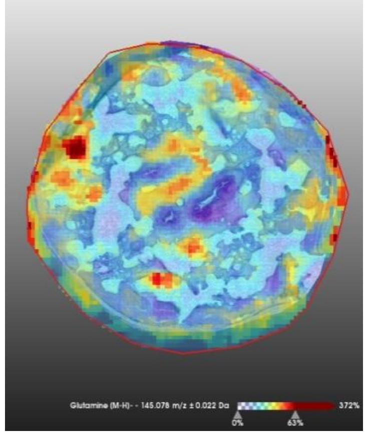

**Glutamine** [M-H]-
m/z 145,078 ± 0,022Da

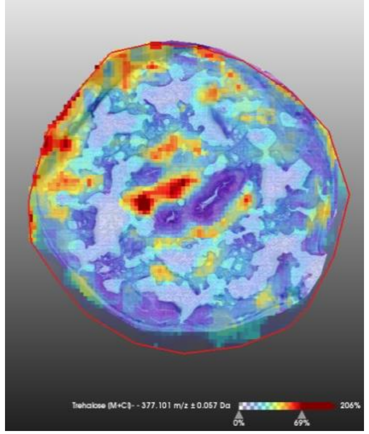

**Trehalose** [M+Cl]-
m/z 377,101 ± 0,057Da

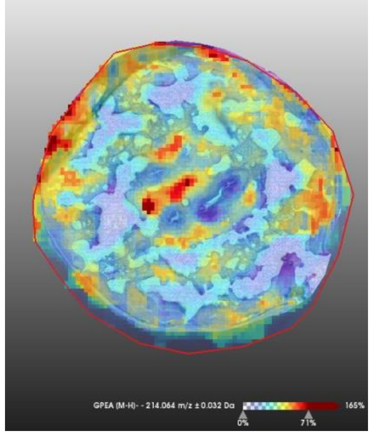

**GPEtn** [M-H]-
m/z 214,064 ± 0,032Da

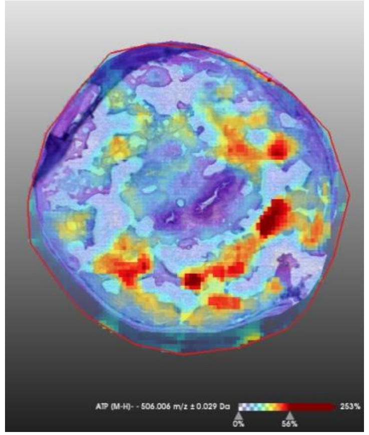

**ATP**[M-H]-
m/z 506,017 ± 0,027Da

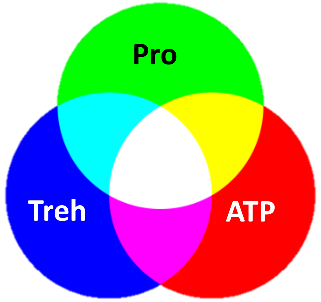


**Color mixing**

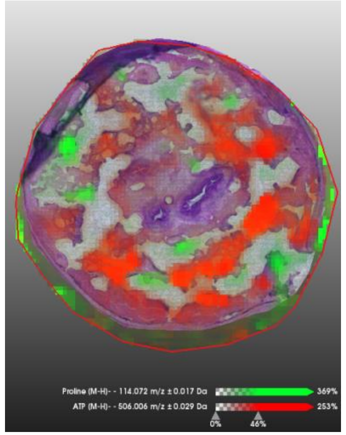

Pro | | ATP

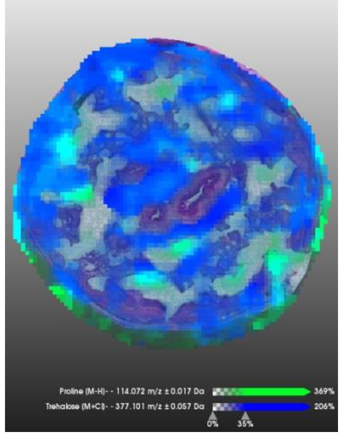

Pro | | Treh

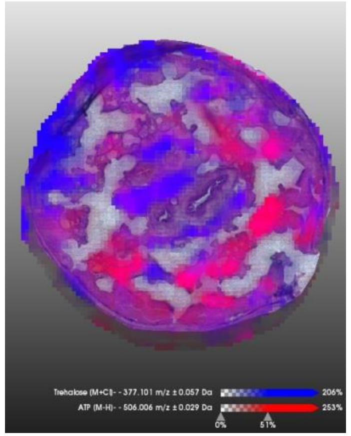

Treh | | ATP

0330B_SDA_A2_Lyo-06

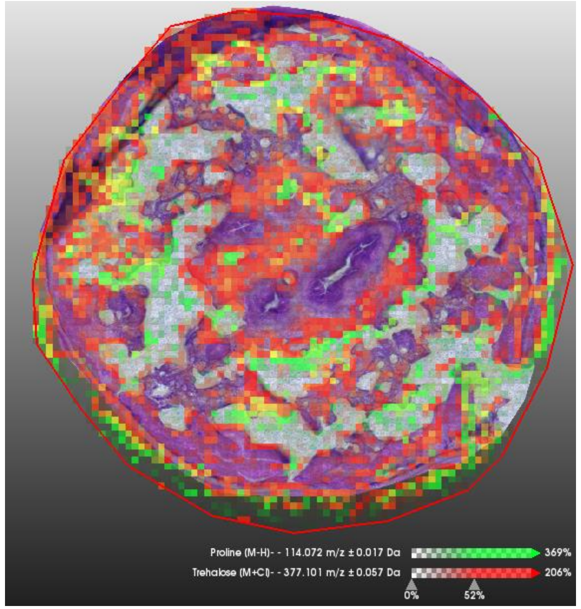


Pro Treh

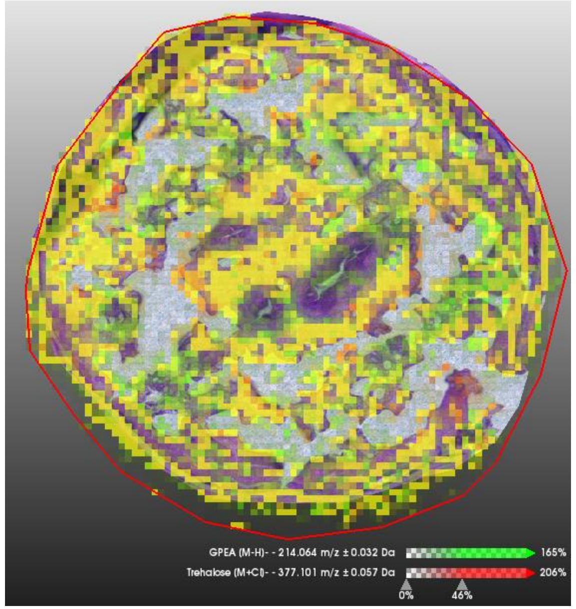


GPE Treh

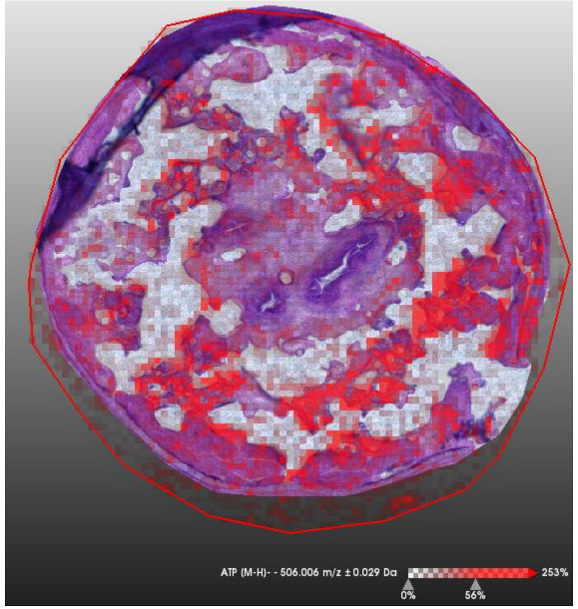


ATP

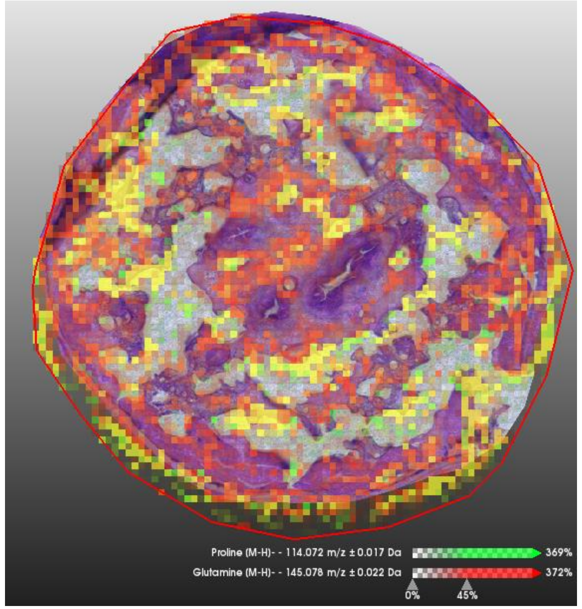


Pro Gln

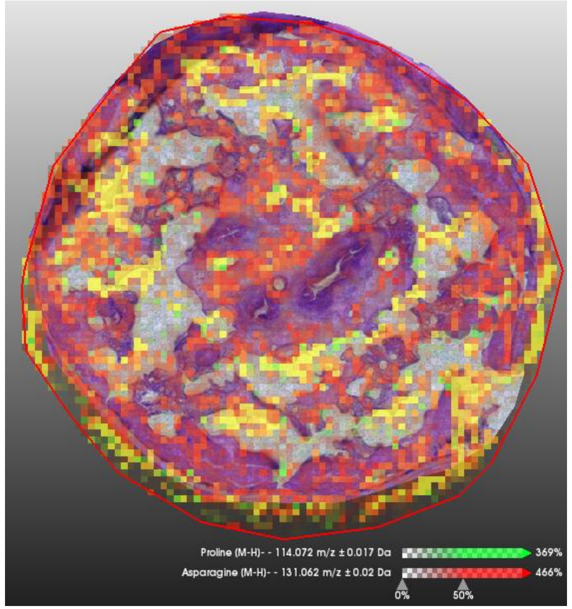


Pro Asn

### 0330B_SDA_A2_frozen-10

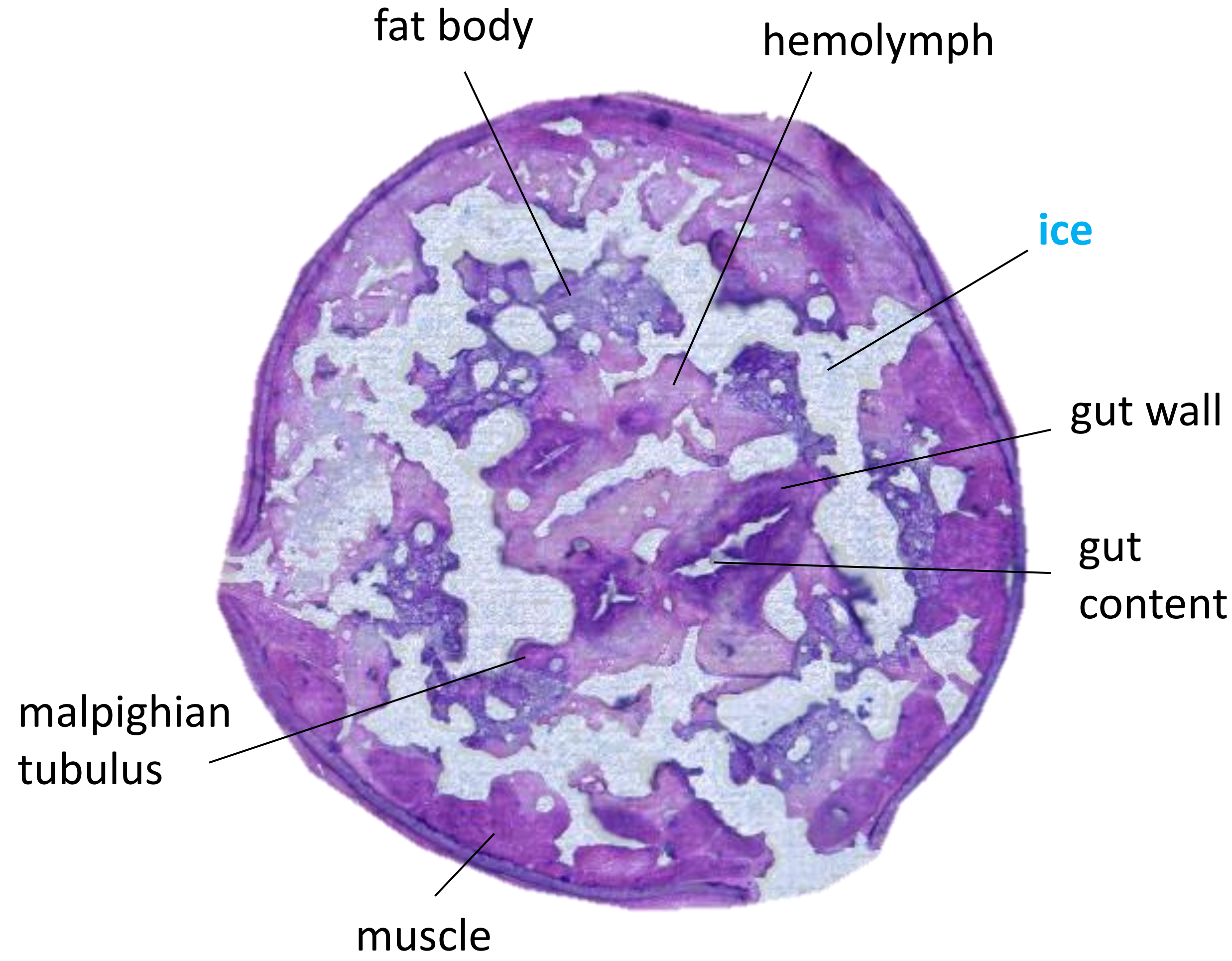


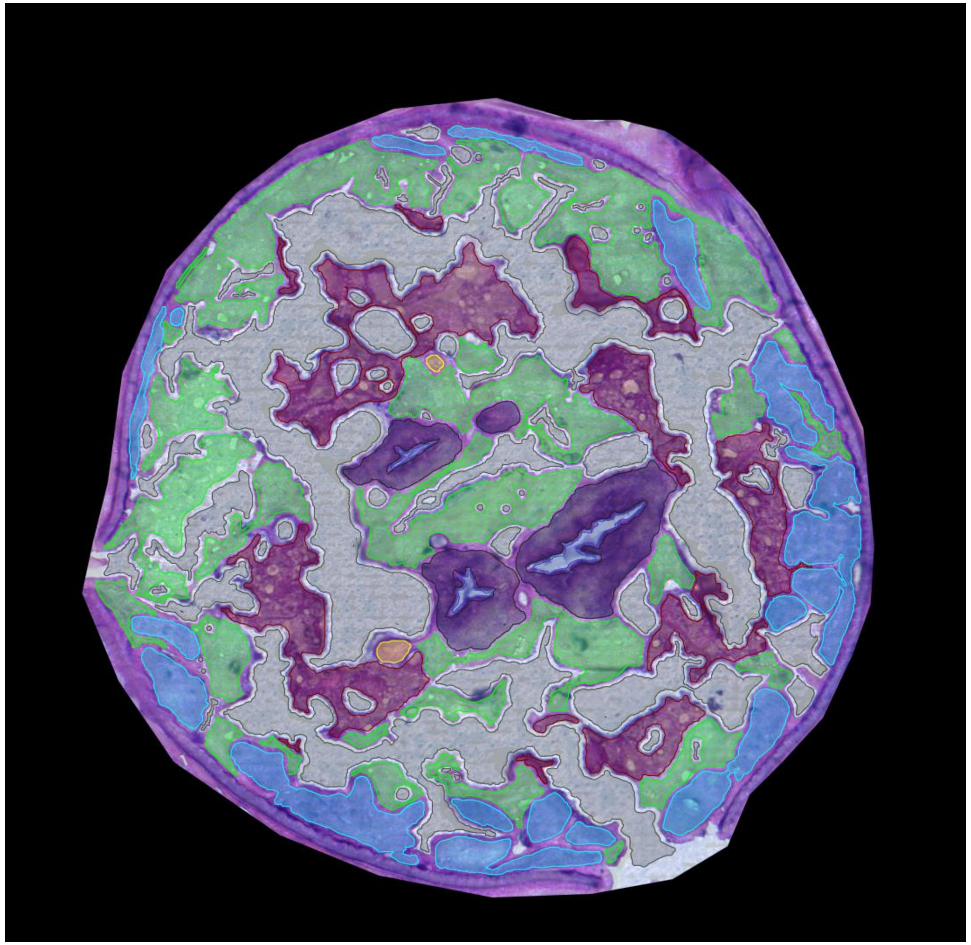

0330B_SDA_A2_Lyo-10

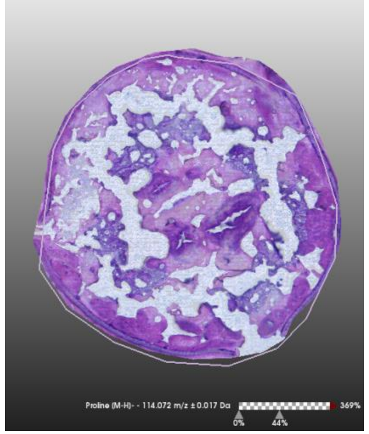

**H&E stain**

**Size bar** 200 µm

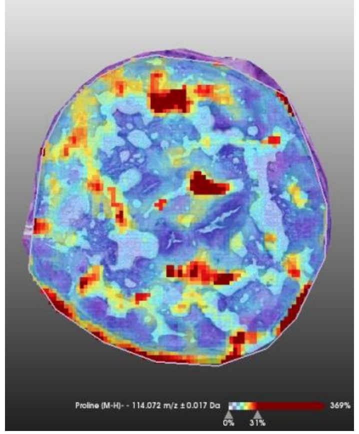

**Proline** [M-H]-
m/z 114,072 ± 0,017Da

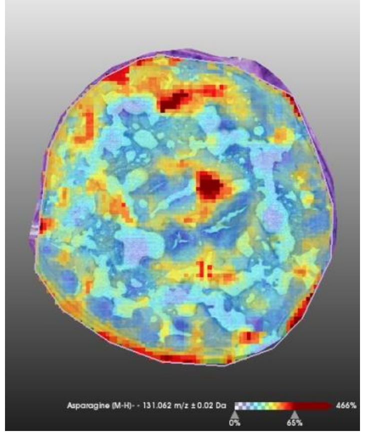

**Asparagine** [M-H]-
m/z 131,062 ± 0,02Da

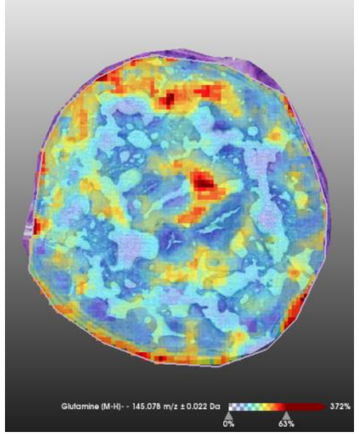

**Glutamine** [M-H]-
m/z 145,078 ± 0,022Da

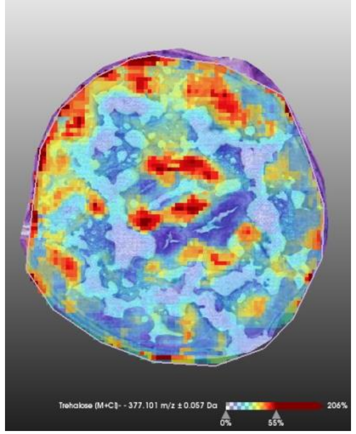

**Trehalose** [M+Cl]-
m/z 377,101 ± 0,057Da

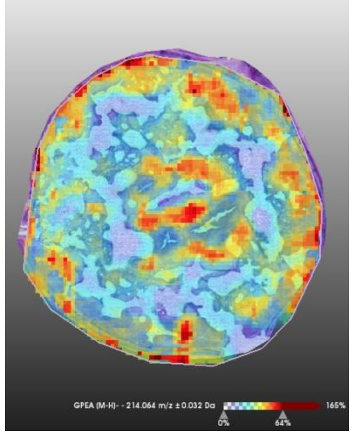

**GPEtn** [M-H]-
m/z 214,064 ± 0,032Da

**Color scale**

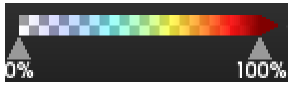


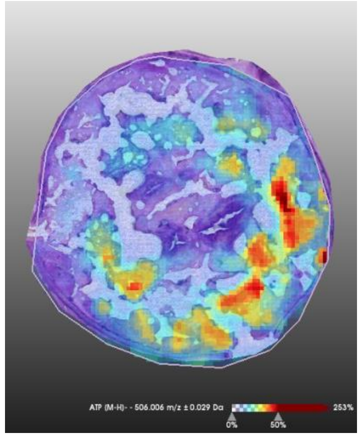

**ATP**[M-H]-
m/z 506,017 ± 0,027Da

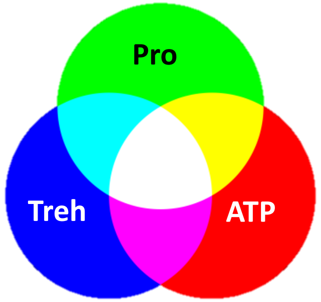


**Color mixing**

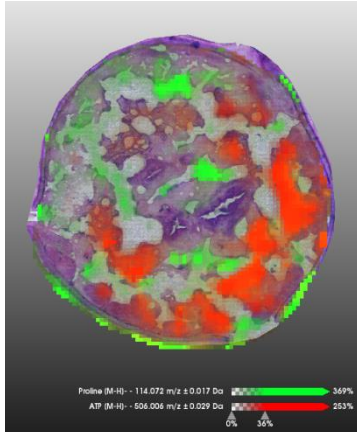

**Pro** | | **ATP**

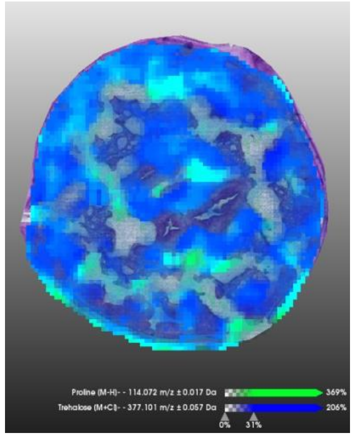

**Pro** | | **Treh**

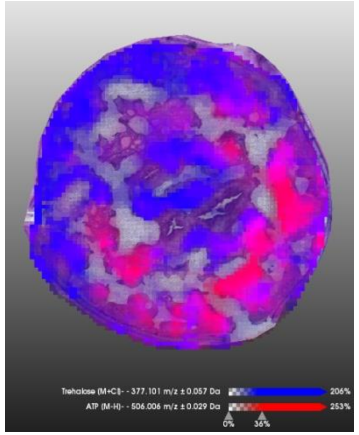

**Treh** | | **ATP**

0330B_SDA_A2_Lyo-10

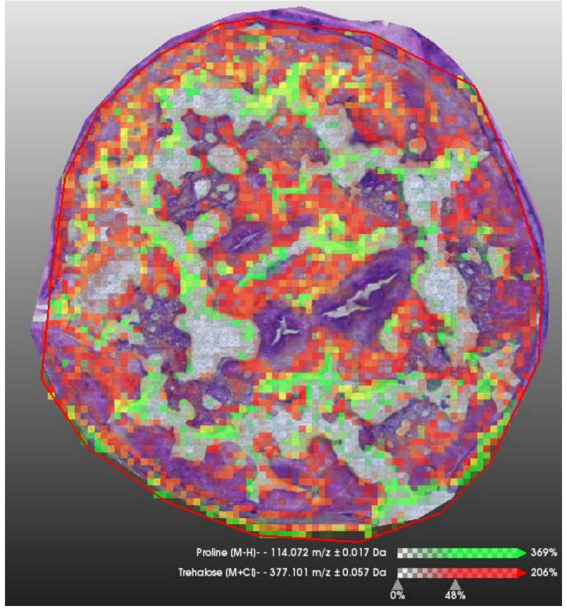


Pro Treh

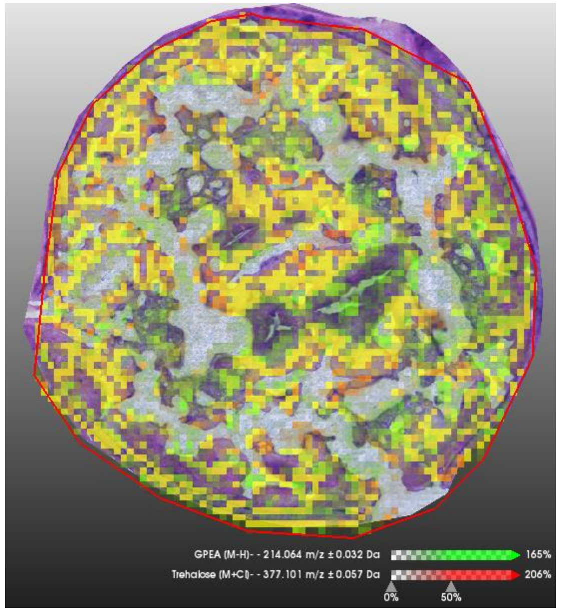


GPE Treh

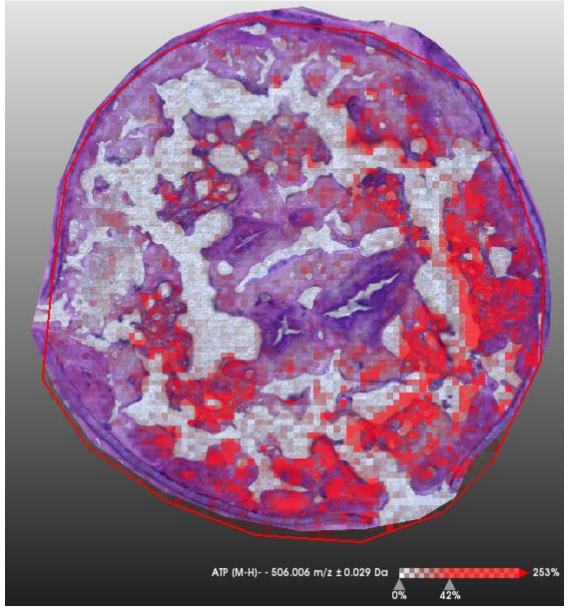


ATP

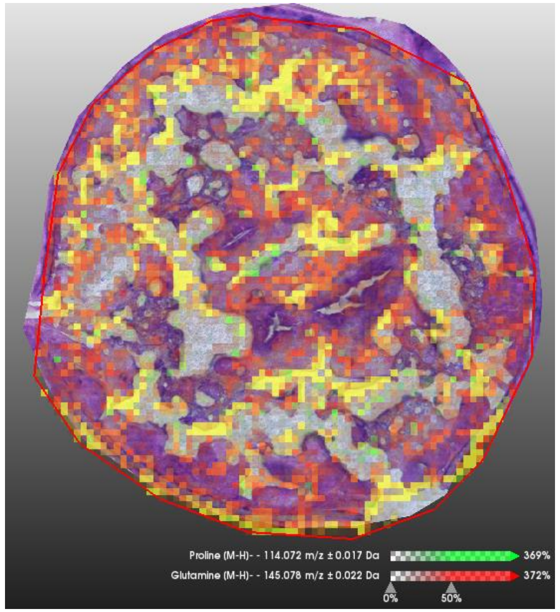


Pro Gln

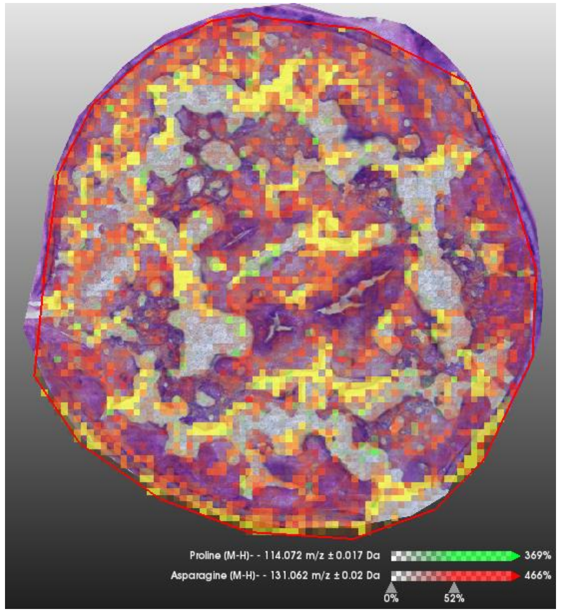


Pro Asn

### 0330B_SDA_A2_frozen-15

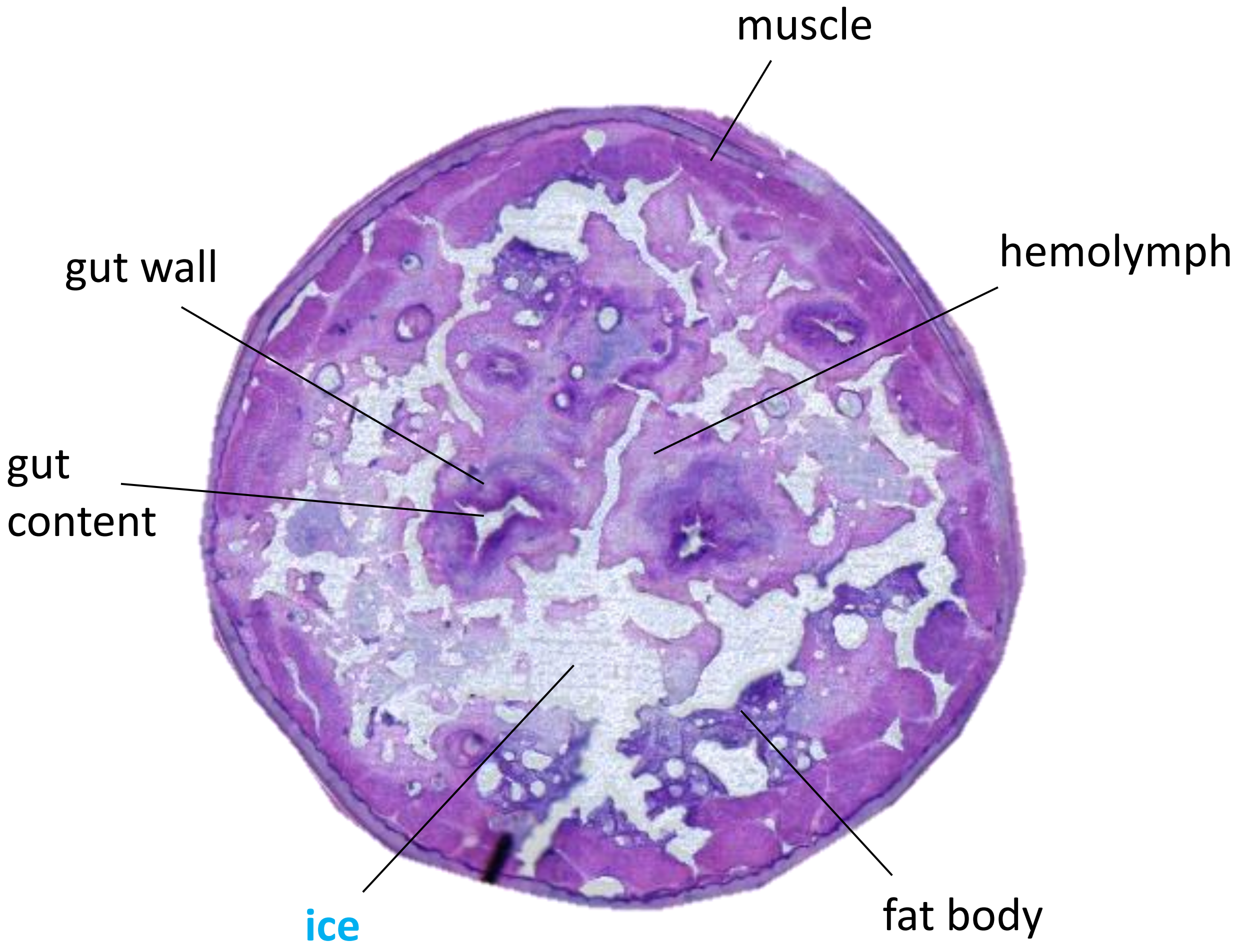


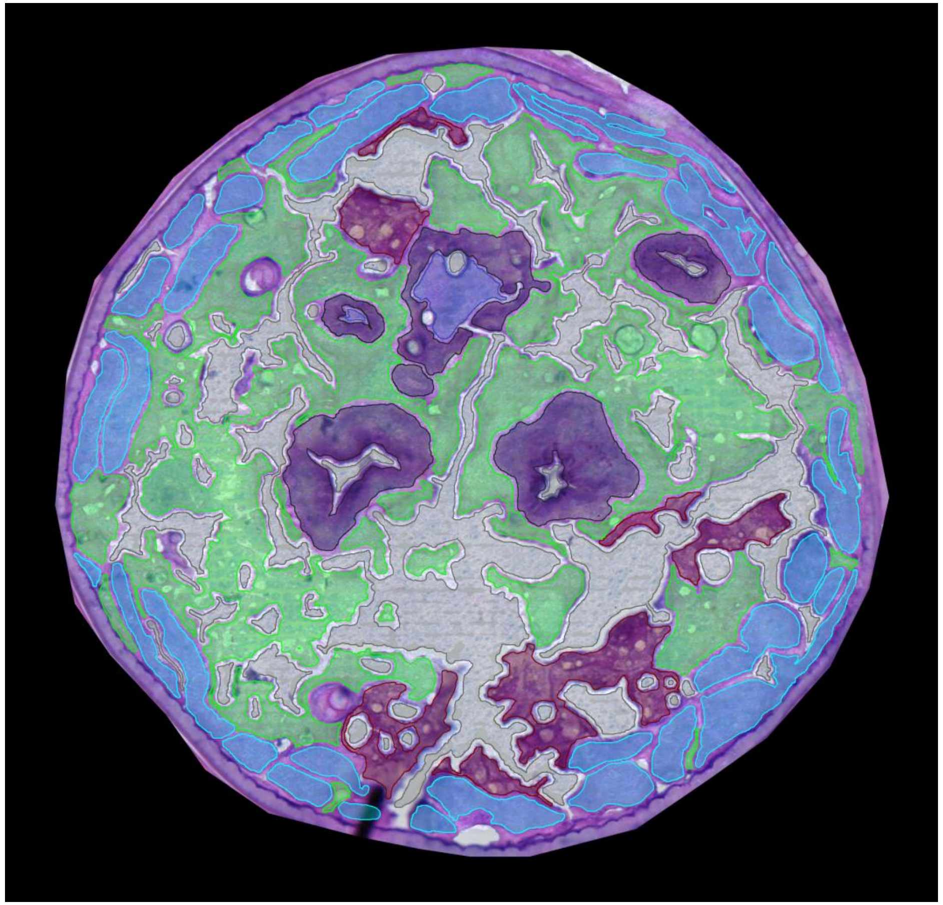

0330B_SDA_A2_Lyo-15

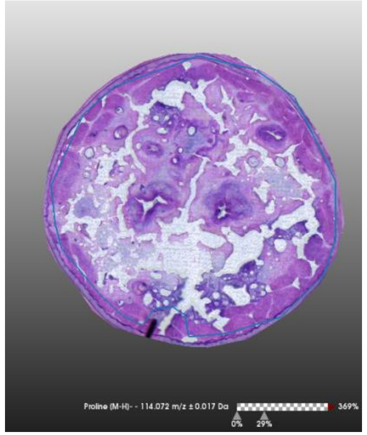

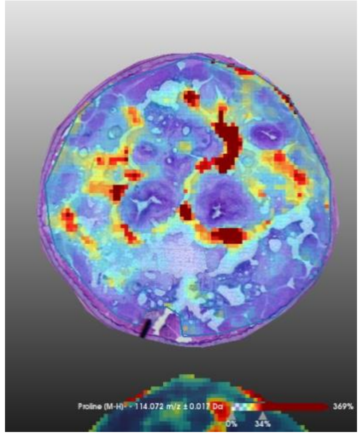

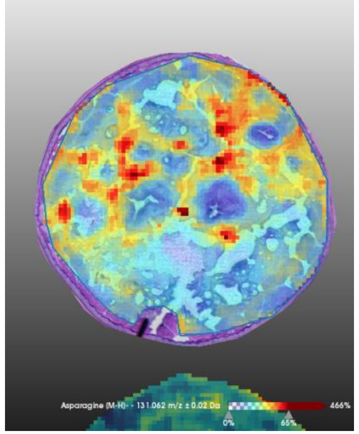

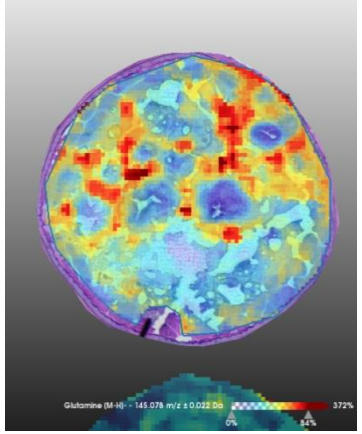

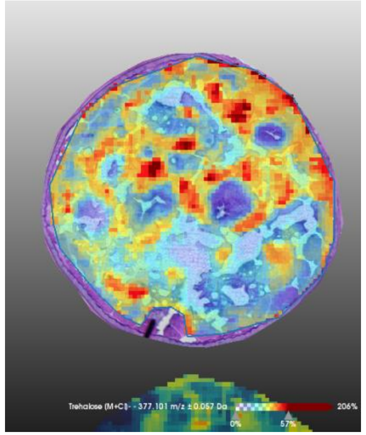

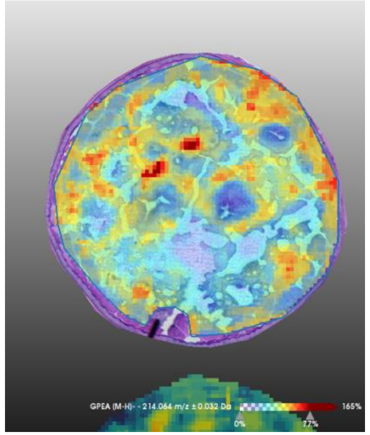

**H&E stain**

**Size bar** 200 µm

**Proline** [M-H]-
m/z 114,072 ± 0,017Da

**Asparagine** [M-H]-
m/z 131,062 ± 0,02Da

**Glutamine** [M-H]-
m/z 145,078 ± 0,022Da

**Trehalose** [M+Cl]-
m/z 377,101 ± 0,057Da

**GPEtn** [M-H]-
m/z 214,064 ± 0,032Da

**Color scale**

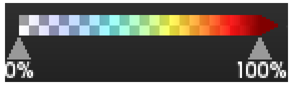


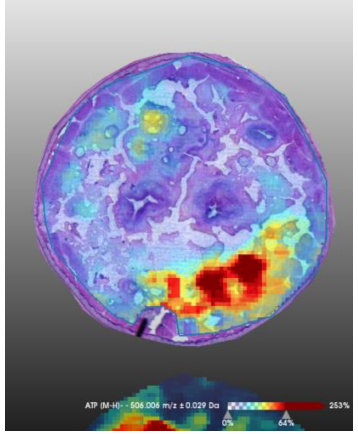

**ATP**[M-H]-
m/z 506,017 ± 0,027Da

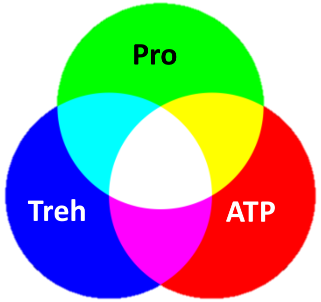


**Color mixing**

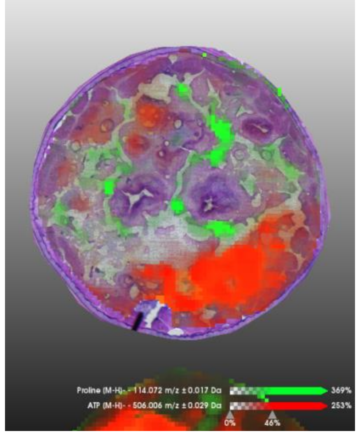

**Pro** **ATP**

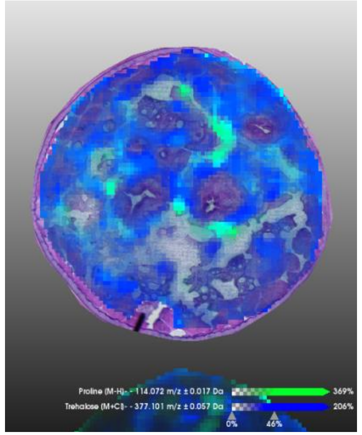

**Pro** **Treh**

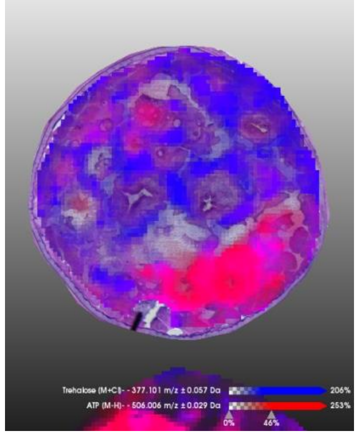

**Treh** **ATP**

0330B_SDA_A2_Lyo-15

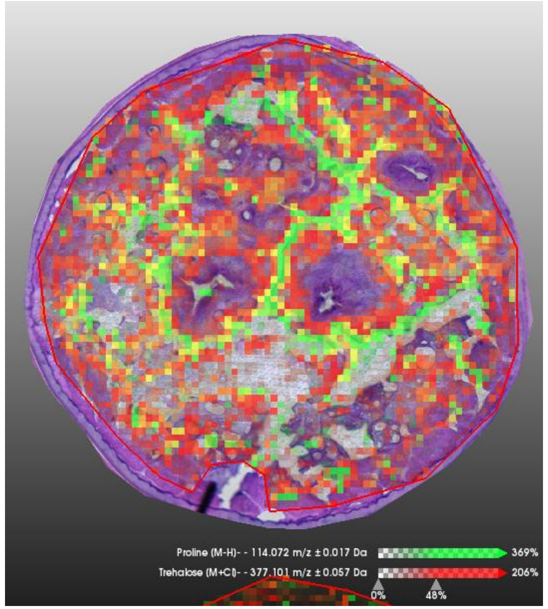


Pro Treh

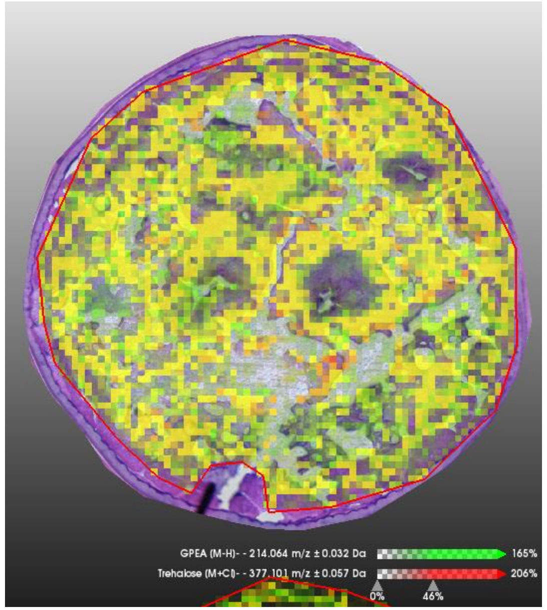


GPE Treh

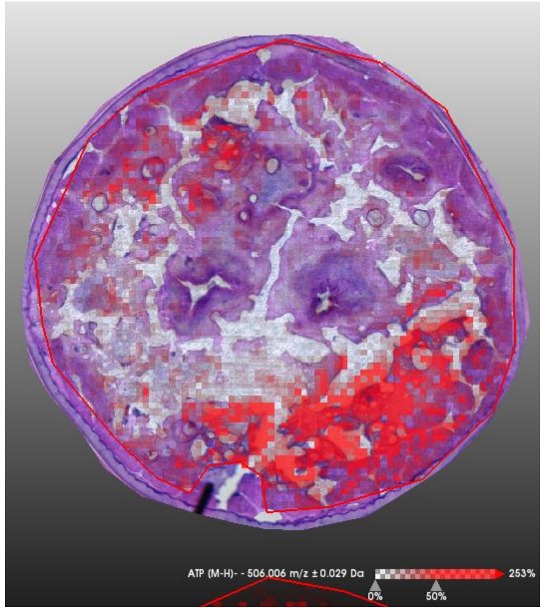


ATP

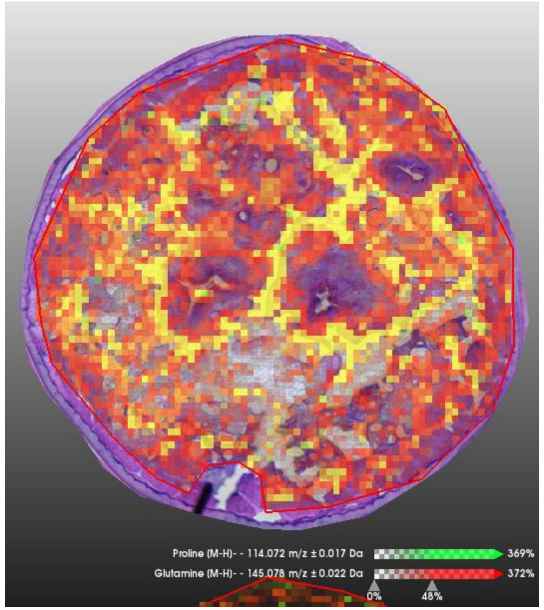


Pro Gln

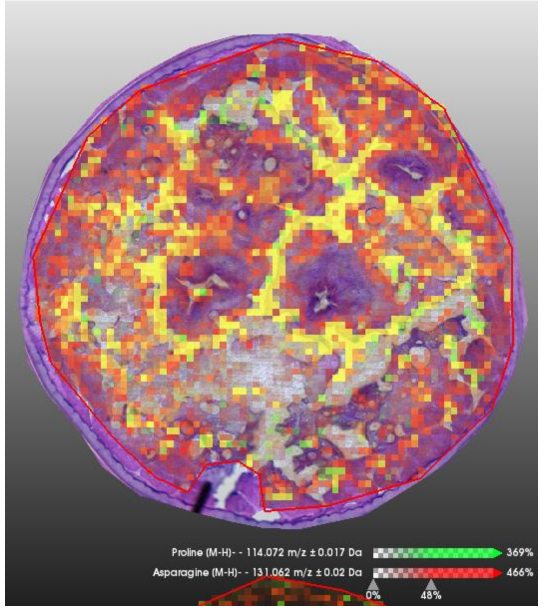


Pro Asn

### 0331B_SDA_B2_frozen-06

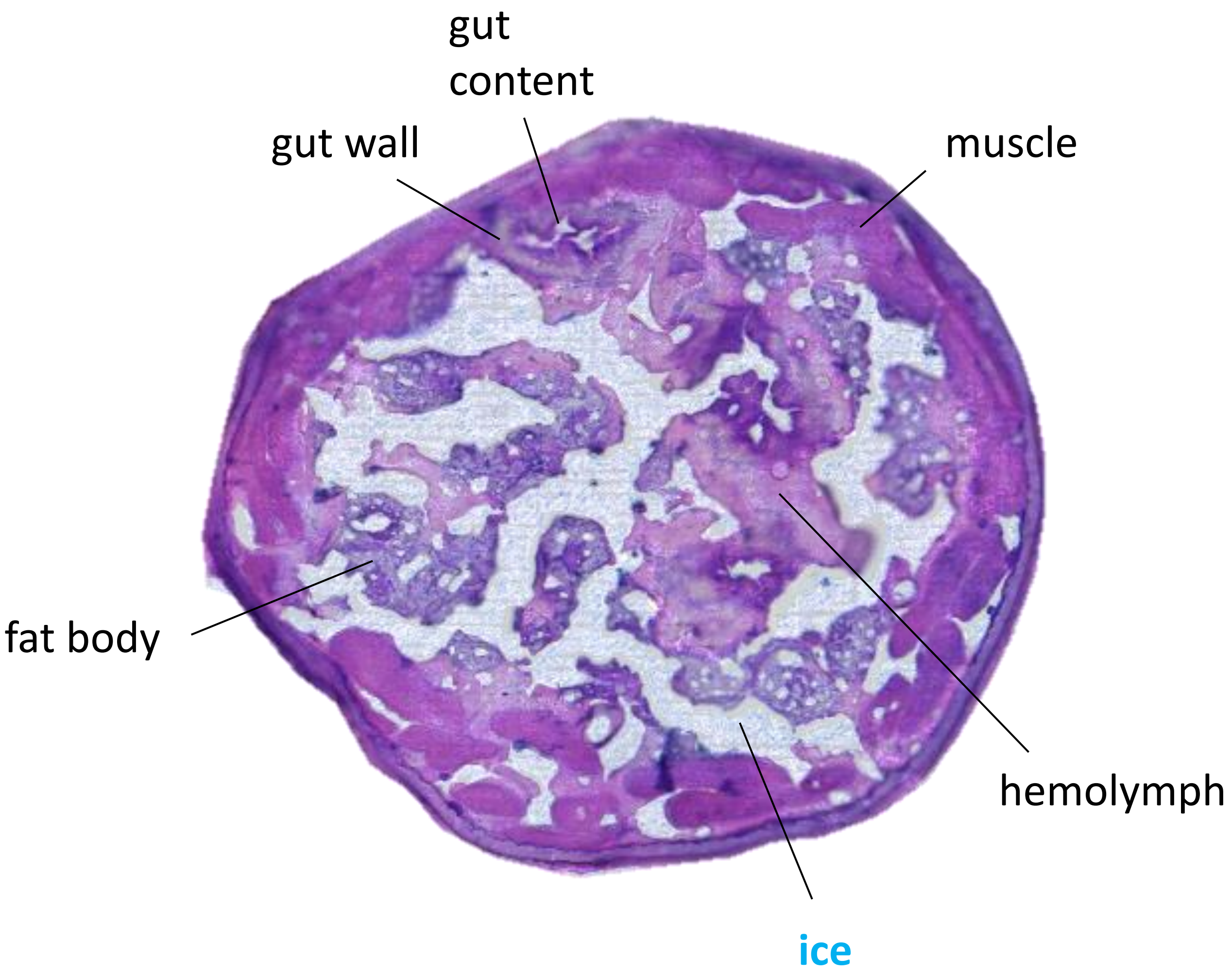


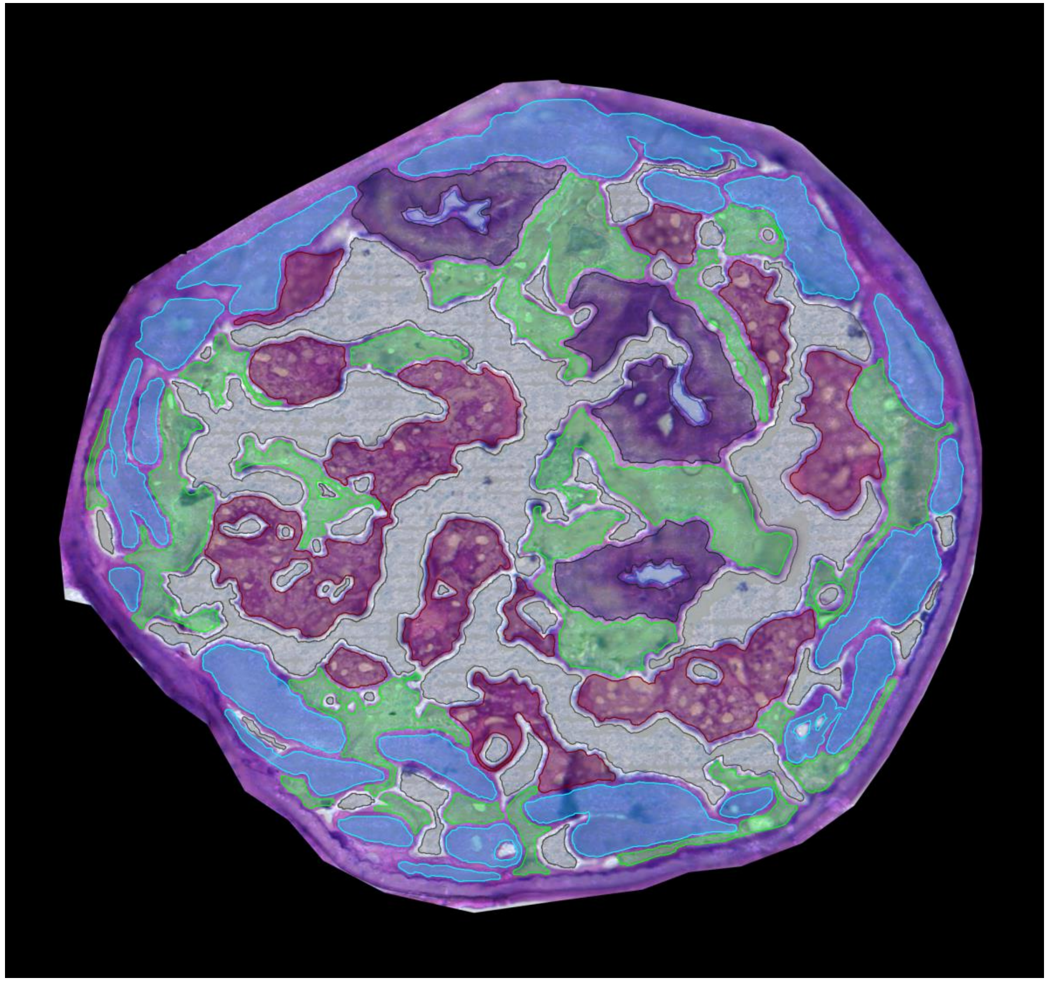

0331B_SDA_B2_Lyo-06

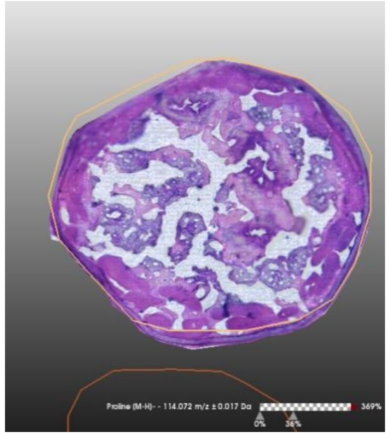

**H&E stain**

**Size bar** 200 µm

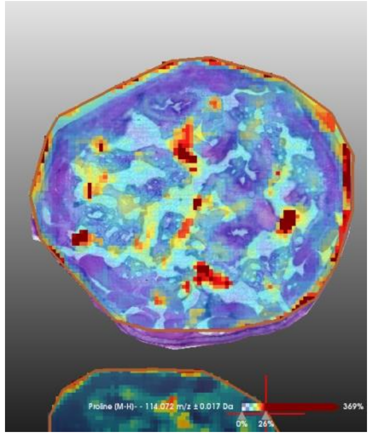

**Proline** [M-H]-
m/z 114,072 ± 0,017Da

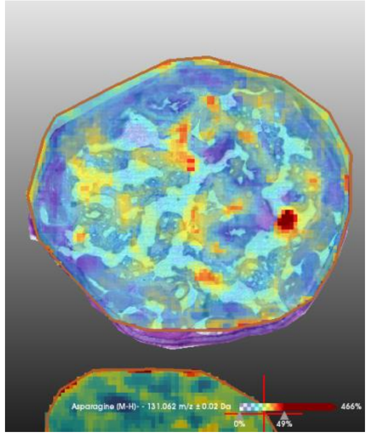

**Asparagine** [M-H]-
m/z 131,062 ± 0,02Da

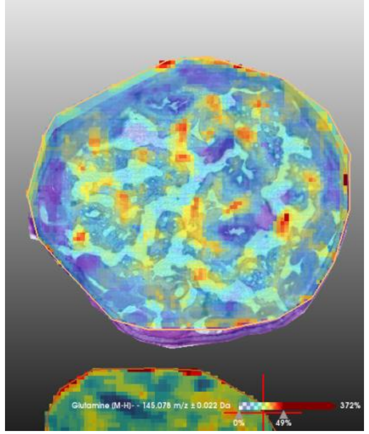

**Glutamine** [M-H]-
m/z 145,078 ± 0,022Da

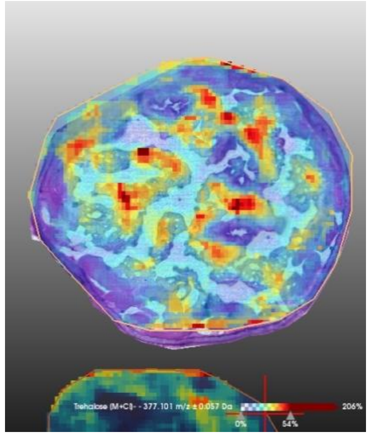

**Trehalose** [M+Cl]-
m/z 377,101 ± 0,057Da

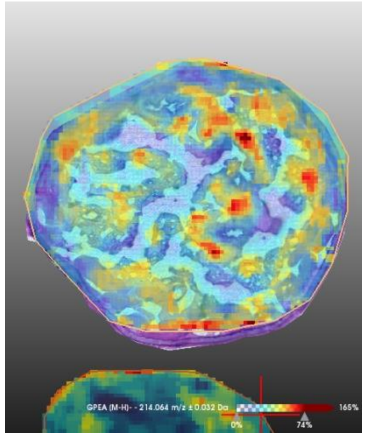

**GPEtn** [M-H]-
m/z 214,064 ± 0,032Da

**Color scale**

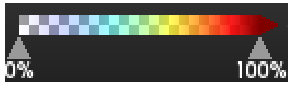


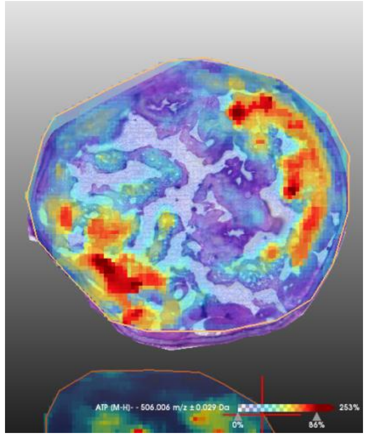

**ATP**[M-H]-
m/z 506,017 ± 0,027Da

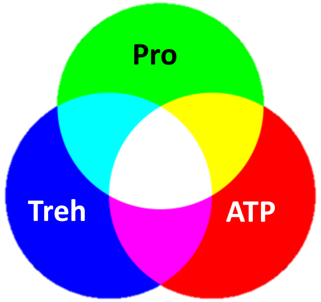


**Color mixing**

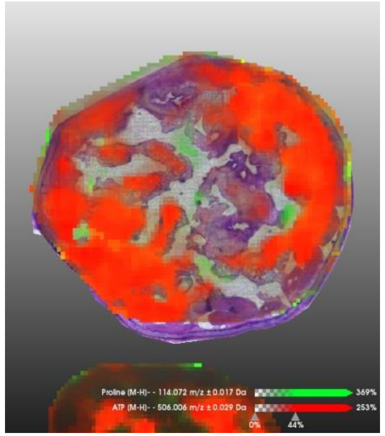

**Pro** **ATP**

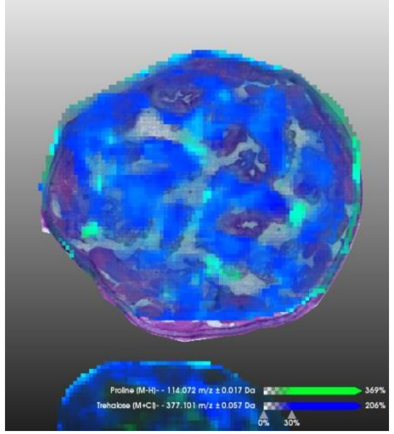

**Pro** **Treh**

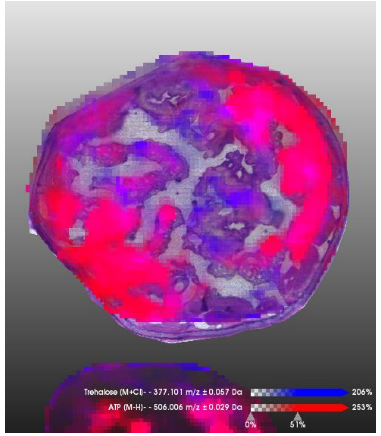

**Treh** **ATP**

0331B_SDA_B2_Lyo-06

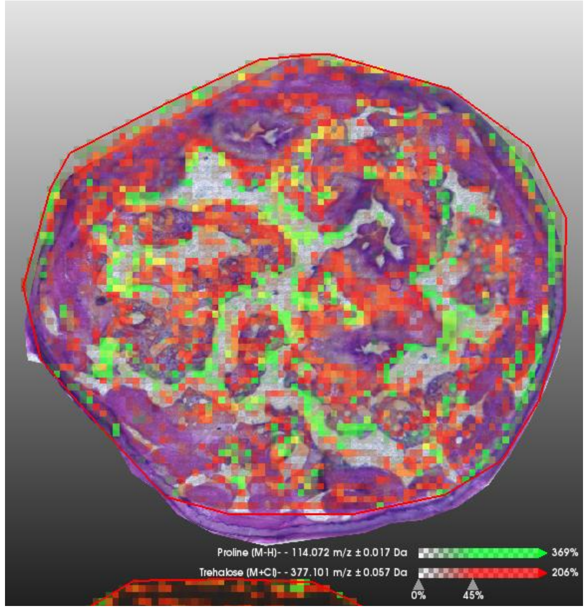


Pro Treh

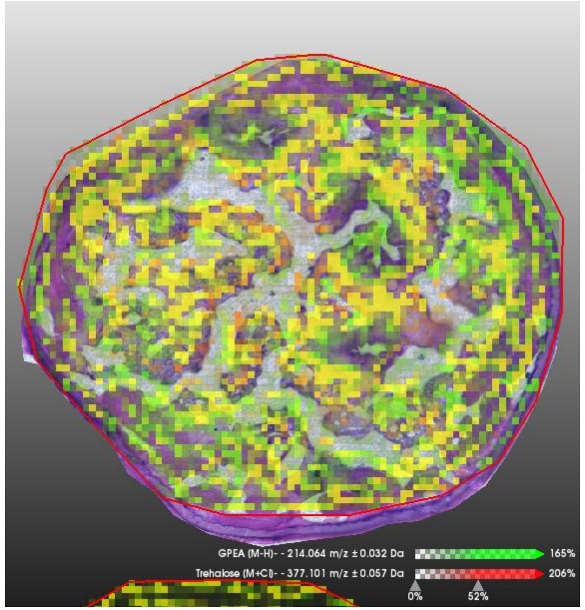


GPE Treh

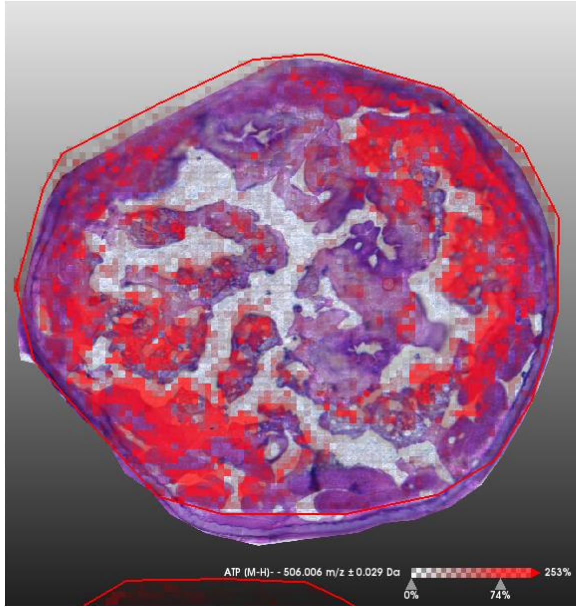


ATP

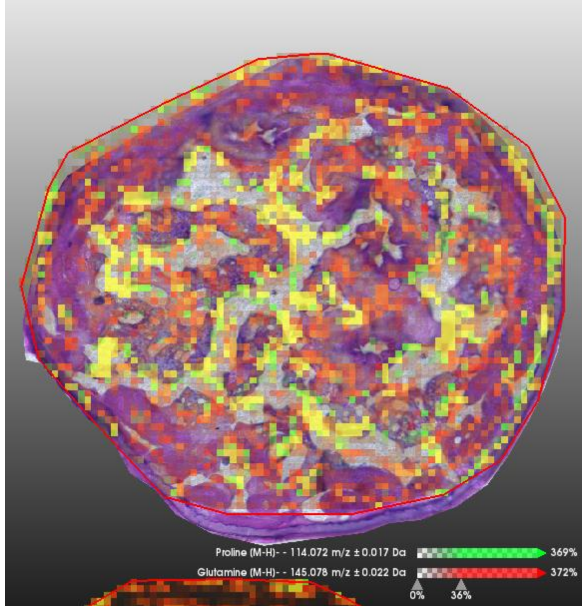


Pro Gln

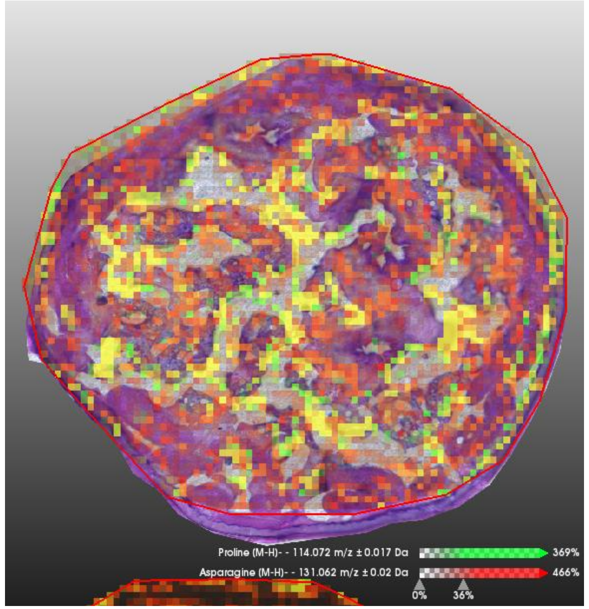


Pro Asn

### 0331B_SDA_B2_frozen-08

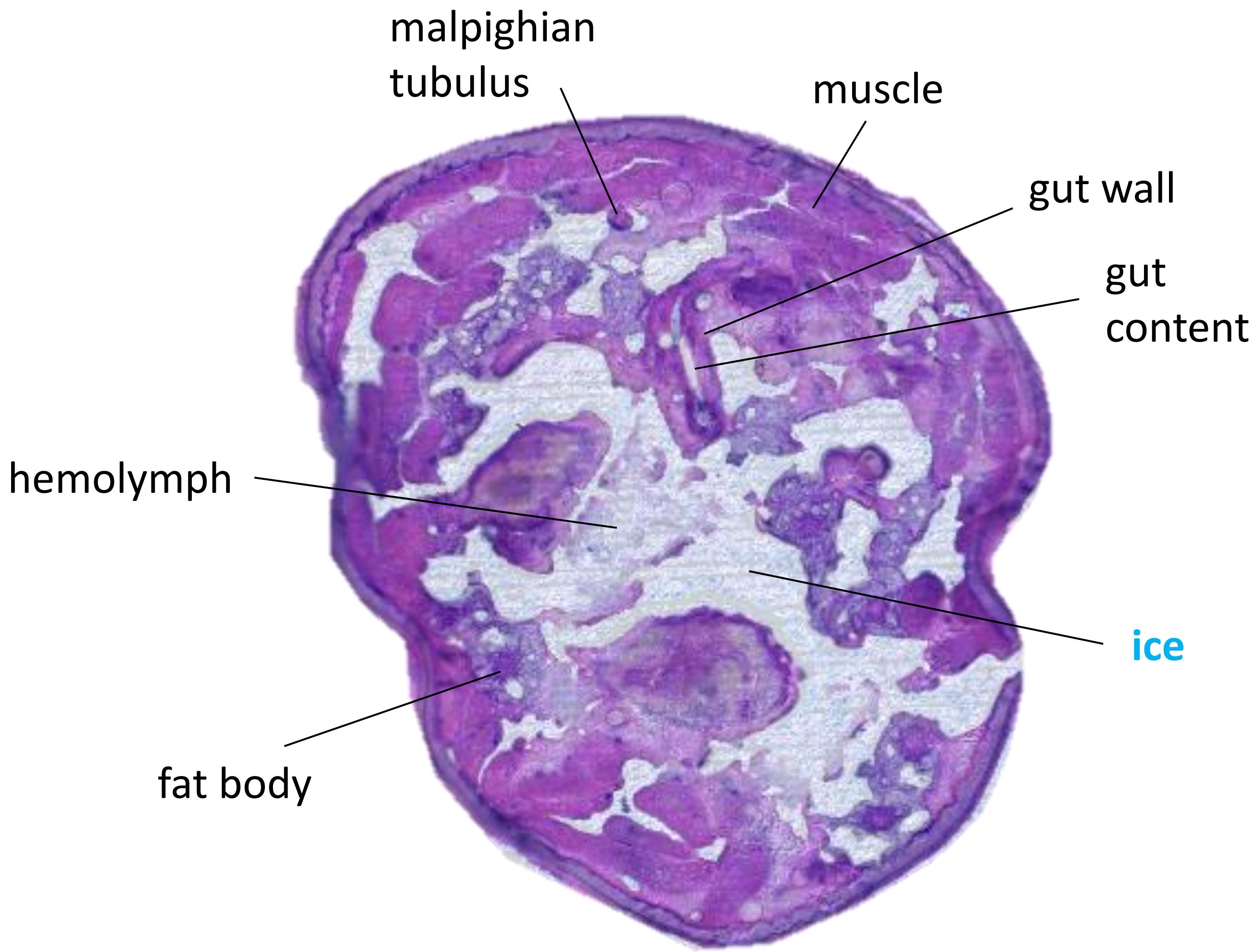


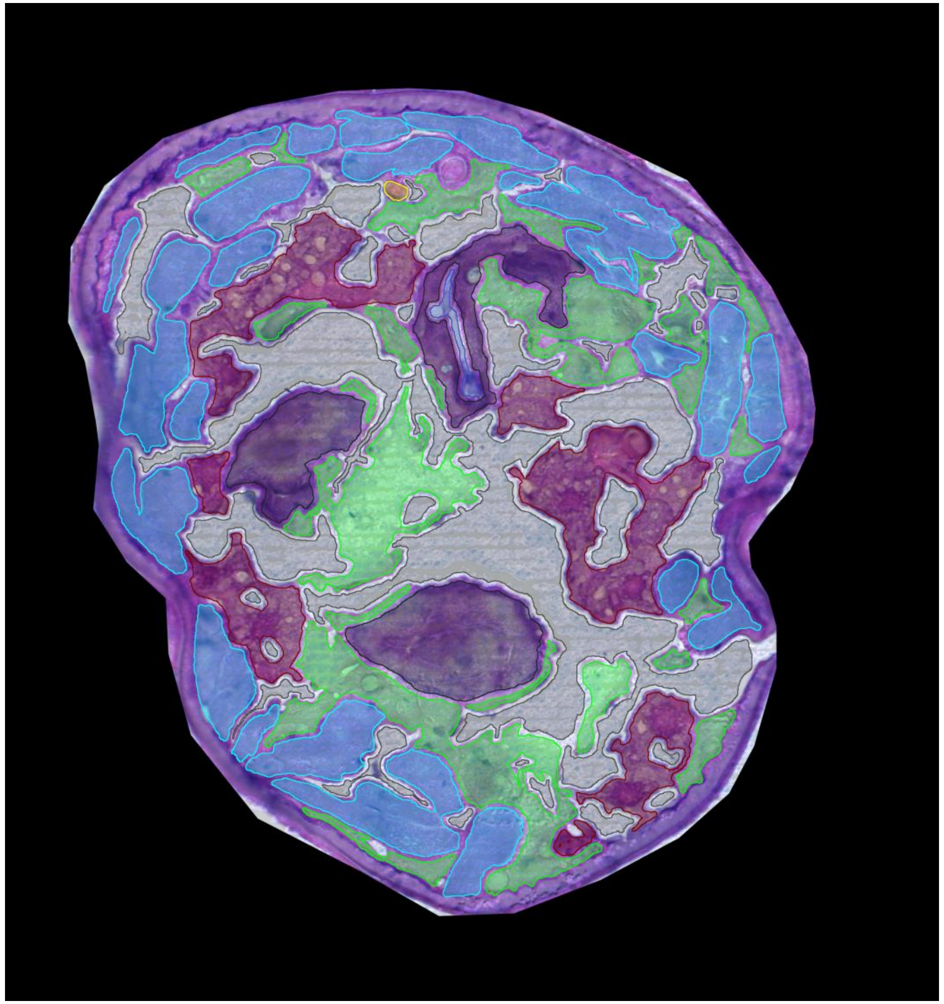

0331B_SDA_B2_Lyo-08

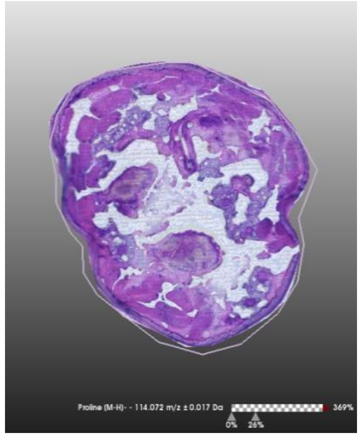

**H&E stain**

**Size bar** 200 µm

**Color scale**

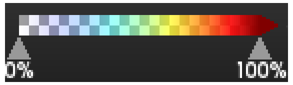


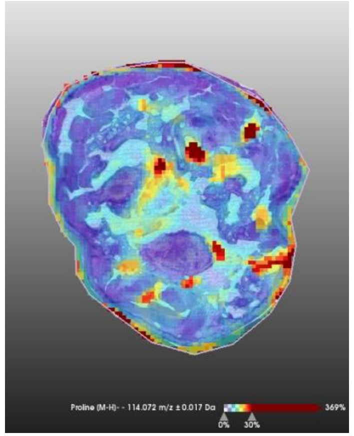

**Proline** [M-H]-
m/z 114,072 ± 0,017Da

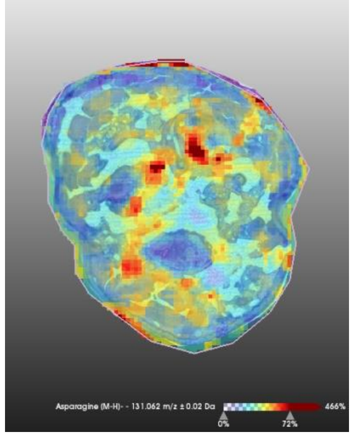

**Asparagine** [M-H]-
m/z 131,062 ± 0,02Da

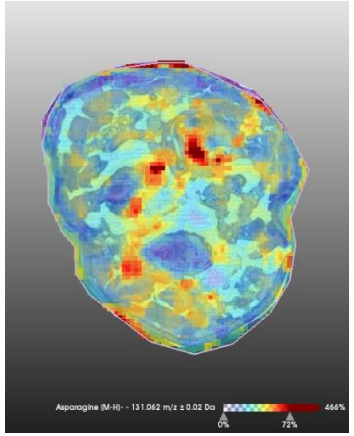

**Glutamine** [M-H]-
m/z 145,078 ± 0,022Da

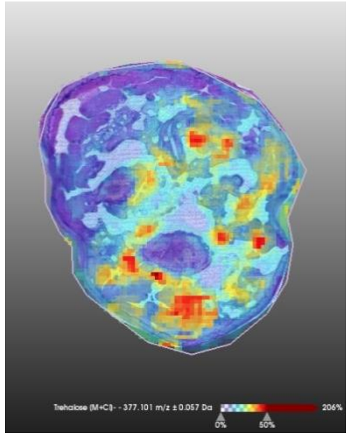

**Trehalose** [M+Cl]-
m/z 377,101 ± 0,057Da

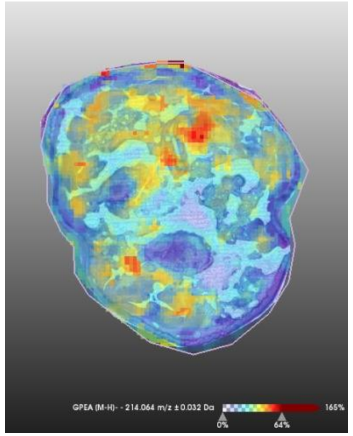

**GPEtn** [M-H]-
m/z 214,064 ± 0,032Da

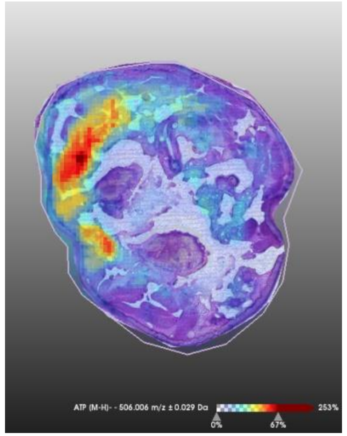

**ATP**[M-H]-
m/z 506,017 ± 0,027Da

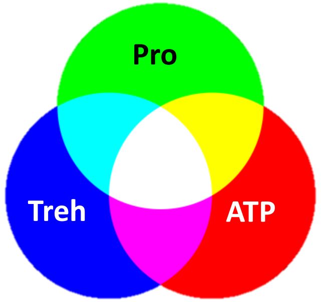


**Color mixing**

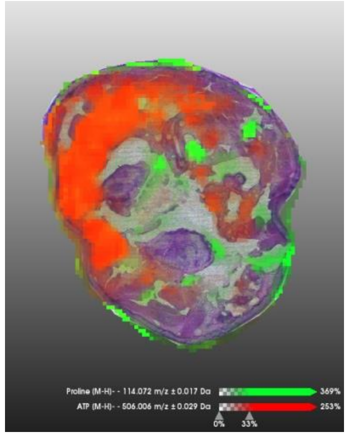

**Pro** **ATP**

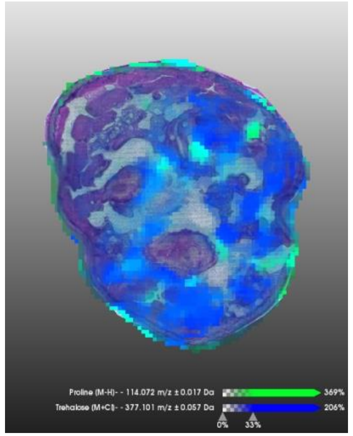

**Pro** **Treh**

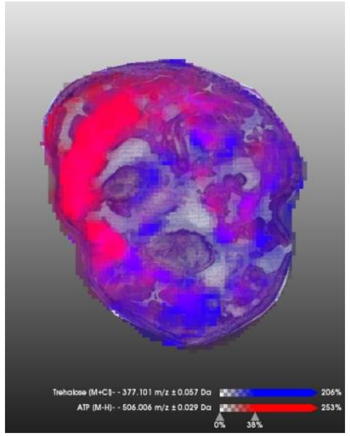

**Treh** **ATP**

0331B_SDA_B2_Lyo-08

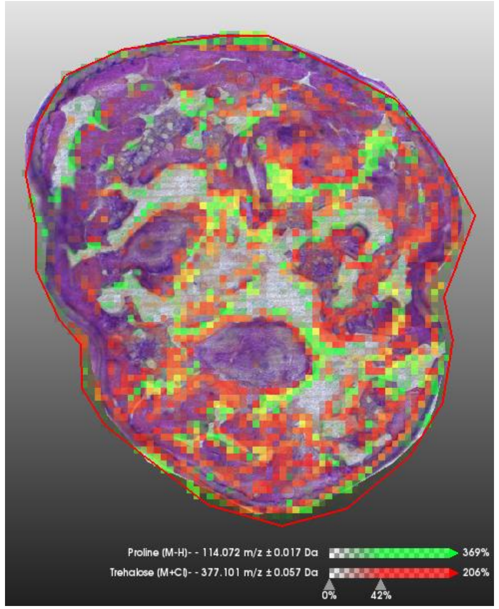


Pro Treh

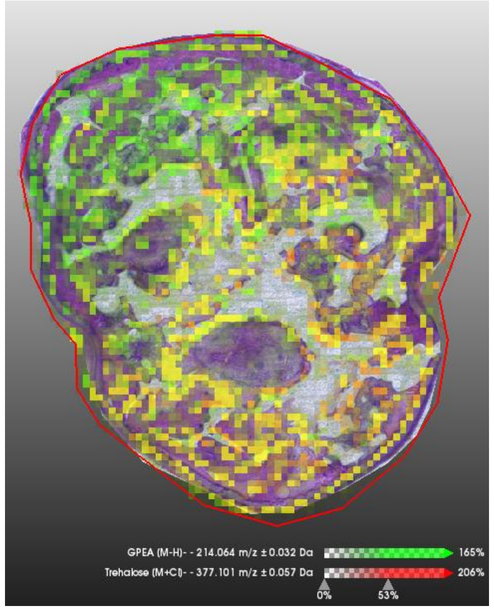


GPE Treh

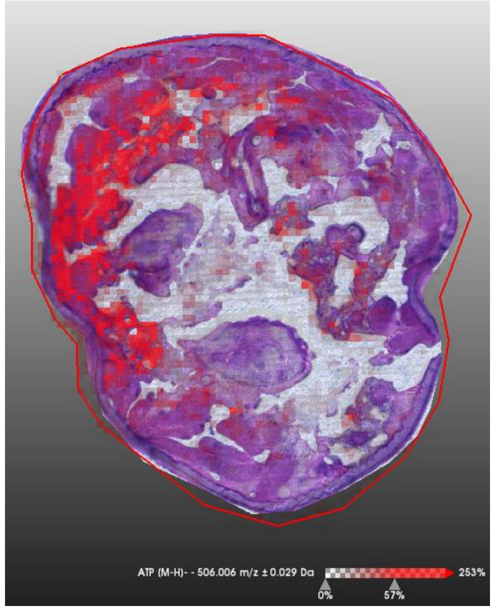


ATP

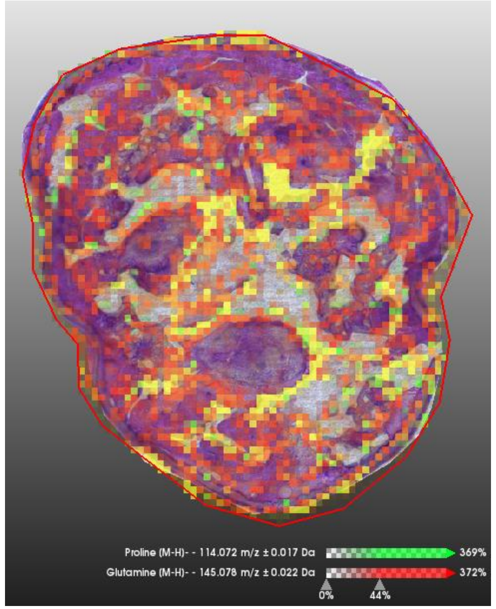


Pro Gln

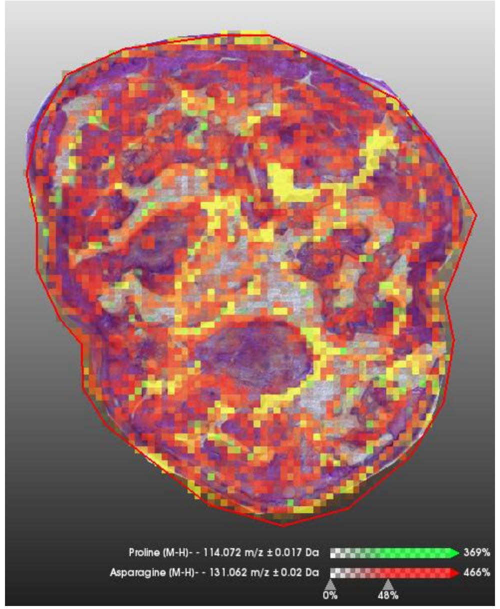


Pro Asn

### 0331B_SDA_B2_frozen-13

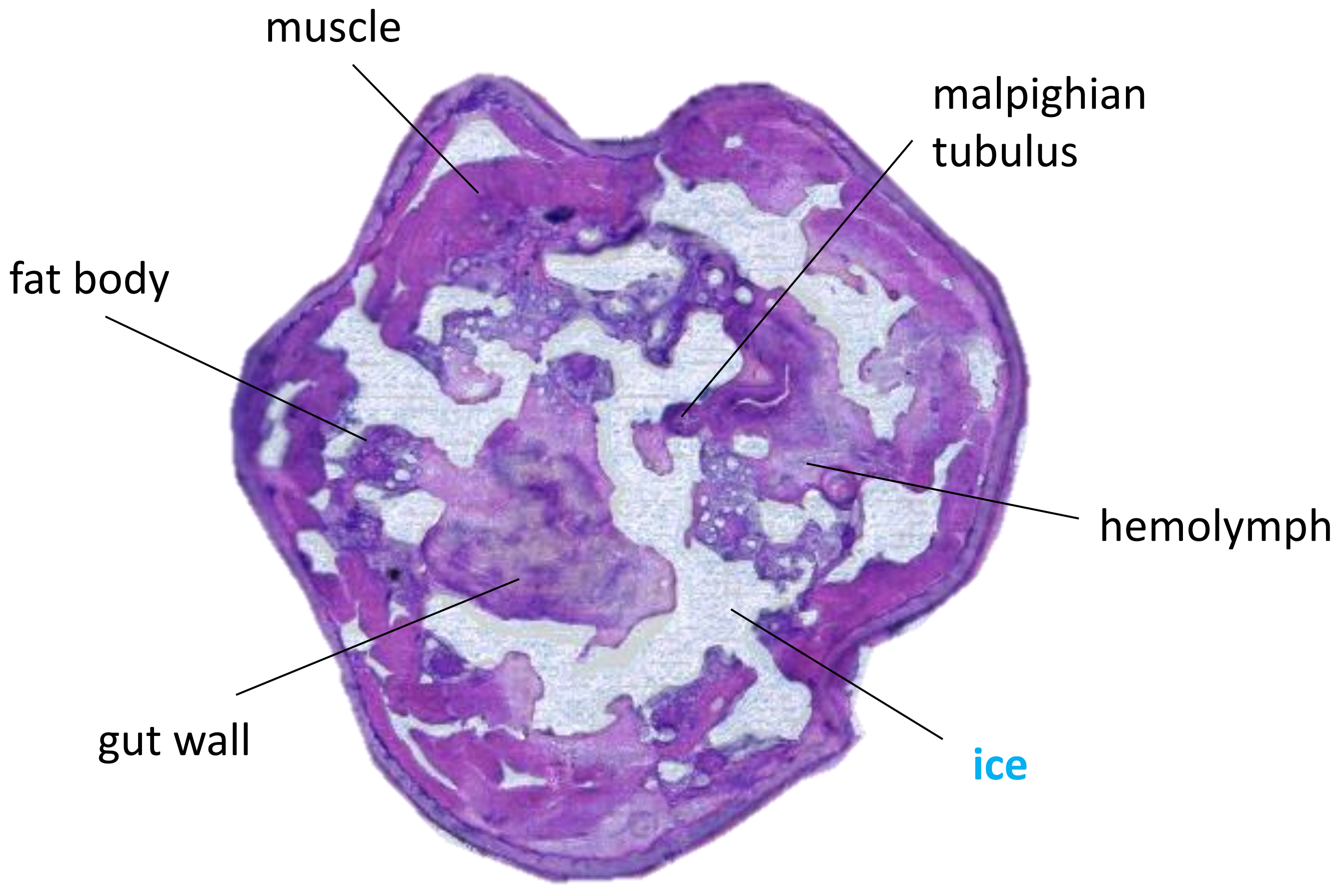


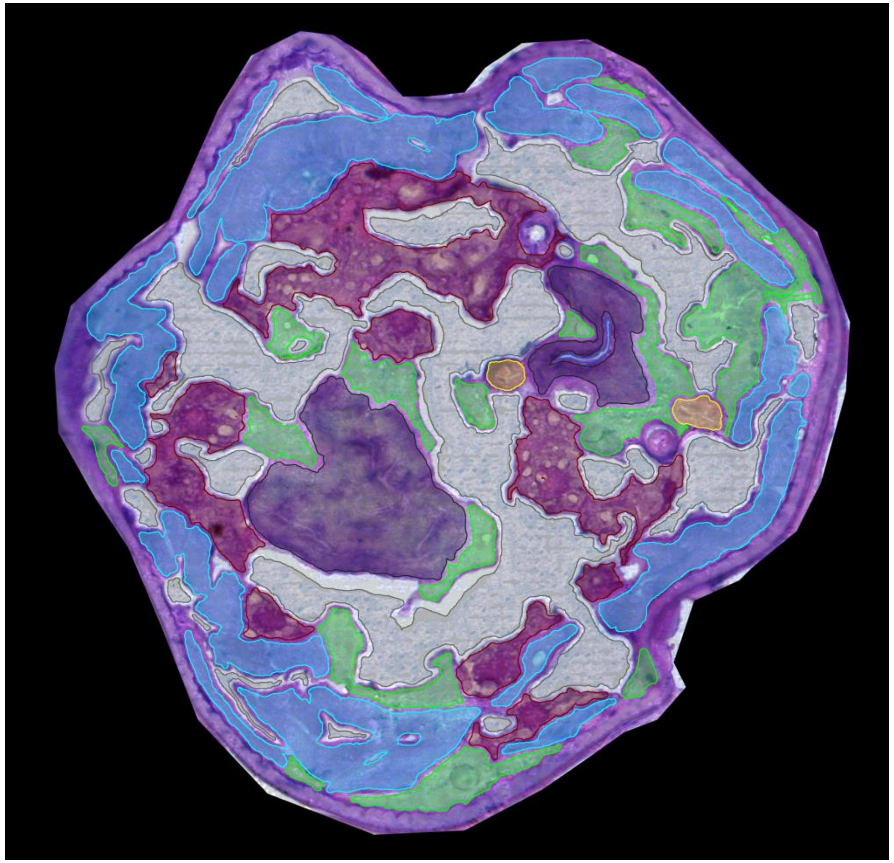

0331B_SDA_B2_Lyo-13

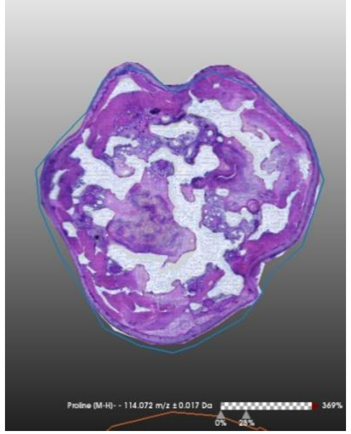

**H&E stain**

**Size bar** 200 µm

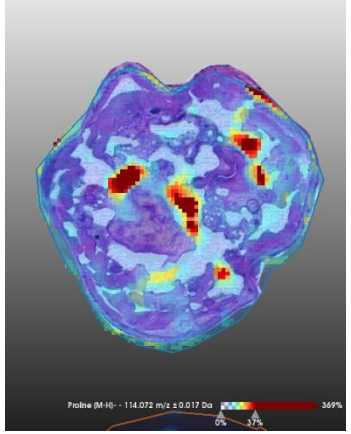

**Proline** [M-H]-
m/z 114,072 ± 0,017Da

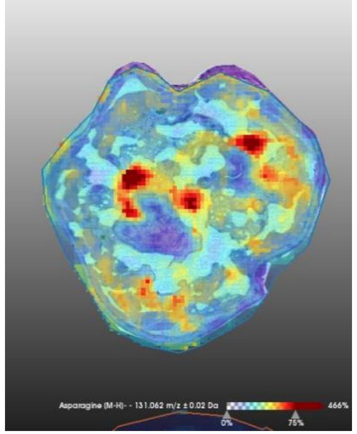

**Asparagine** [M-H]-
m/z 131,062 ± 0,02Da

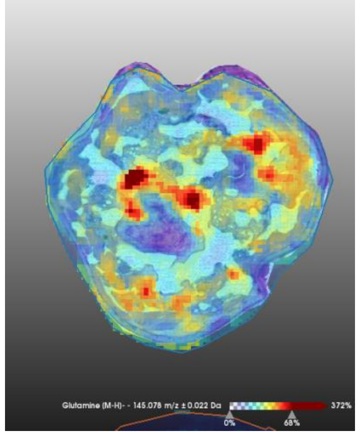

**Glutamine** [M-H]-
m/z 145,078 ± 0,022Da

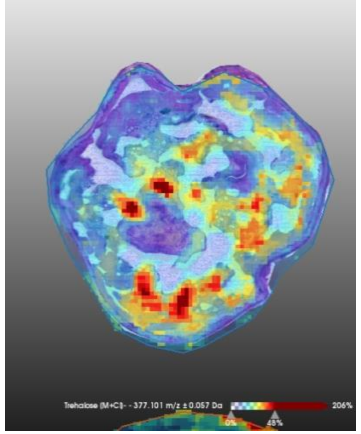

**Trehalose** [M+Cl]-
m/z 377,101 ± 0,057Da

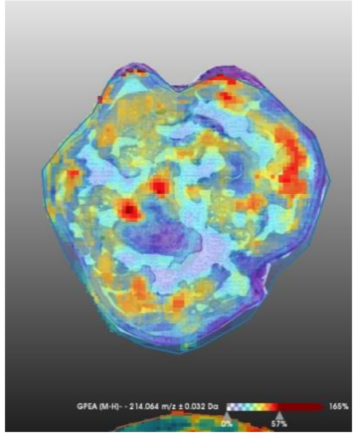

**GPEtn** [M-H]-
m/z 214,064 ± 0,032Da

**Color scale**

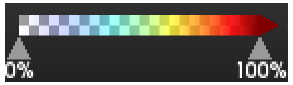


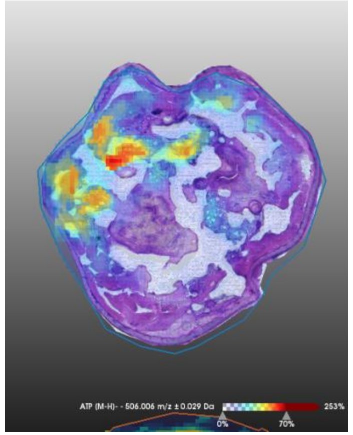

**ATP**[M-H]-
m/z 506,017 ± 0,027Da

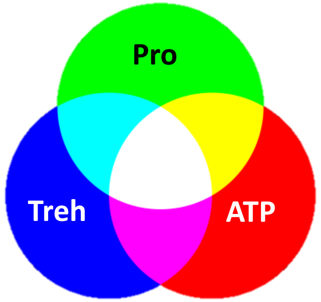


**Color mixing**

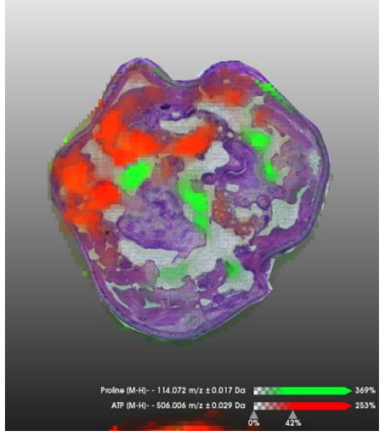

Pro | ATP

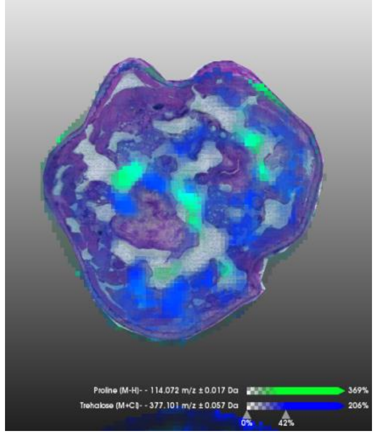

Pro | Treh

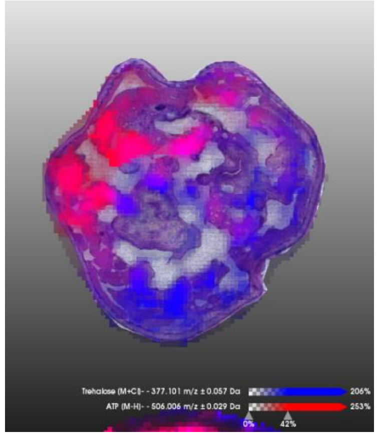

Treh | ATP

0331B_SDA_B2_Lyo-13

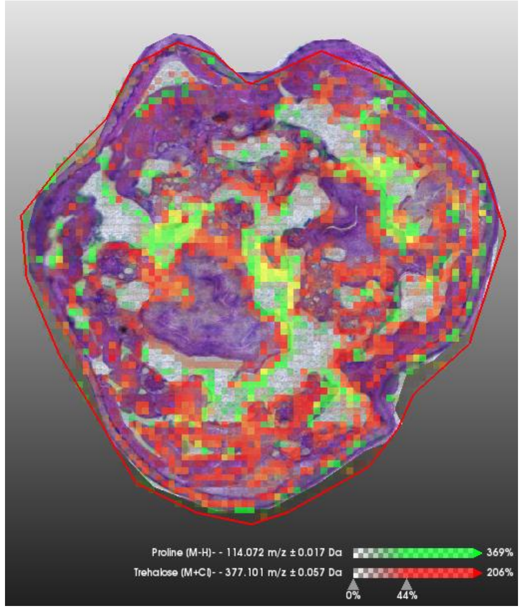


Pro | Treh

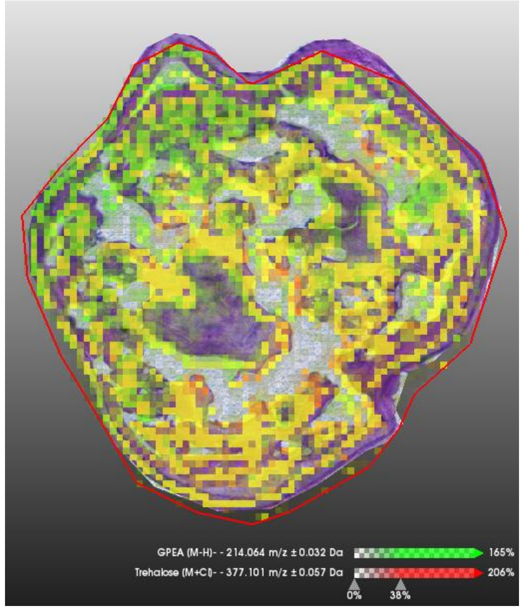


GPE | Treh

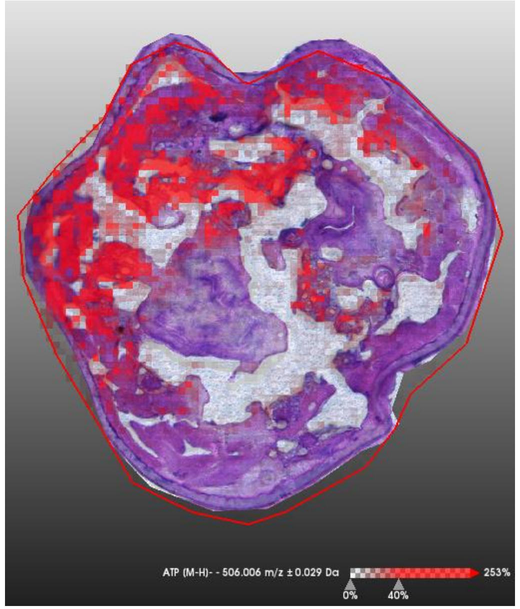


ATP

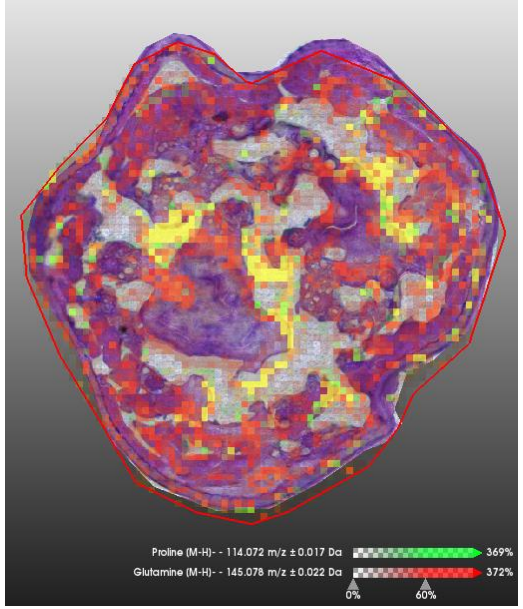


Pro | Gln

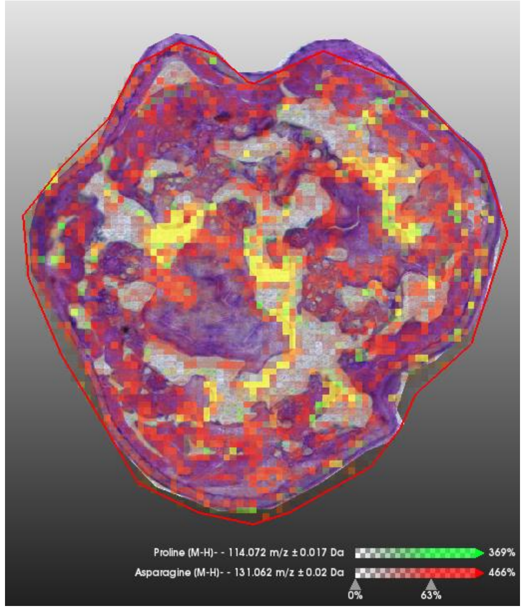


Pro | Asn
