## Supplementary Figures for "A mixture of innate cryoprotectants is key for freeze tolerance and cryopreservation of a drosophilid fly larva"

| | |
|---|---|
| **Supplementary Figure A1** | Phenotypic/acclimation variants and freeze tolerance bioassay. |
| **Supplementary Figure A2** | Preparation of SDA-frozen larvae for MALDI-TOF/MS analysis. |
| **Supplementary Figure A3** | Metabolomic profiles in *C. costata* tissues. |
| **Supplementary Figure A4** | Sizes of metabolite pools in *C. costata* tissues. |
| **Supplementary Figure A5** | MALDI-MSI of Gln, Asn and GPE on a longitudinal section of an SDA larva. |
| **Supplementary Figure A6** | Relative quantification of MALDI-TOF signals: method and limitations. |
| **Supplementary Figure A7** | Relative quantification of MALDI-TOF signals in LD larvae. |
| **Supplementary Figure A8** | Relative quantification of MALDI-TOF signals in SDA larvae. |
| **Supplementary Figure A9** | Relative quantification of MALDI-TOF signals in SDA-frozen larvae. |
| **Supplementary Figure A10** | Analysis of DSC heating curves of trehalose solution (338 $mmol.kg^{-1}$) in water. |
| **Supplementary Figure A11** | Analysis of DSC heating curves of proline solution (978 $mmol.kg^{-1}$) in water. |
| **Supplementary Figure A12** | Analysis of DSC heating curves of glutamine solution (172 $mmol.kg^{-1}$) in water. |
| **Supplementary Figure A13** | Validation of trehalose m/z peak identity using $^{13}C6$ experiment. |
| **Supplementary Figure A14** | Validation of proline m/z peak identity using $^{13}C6$ experiment. |

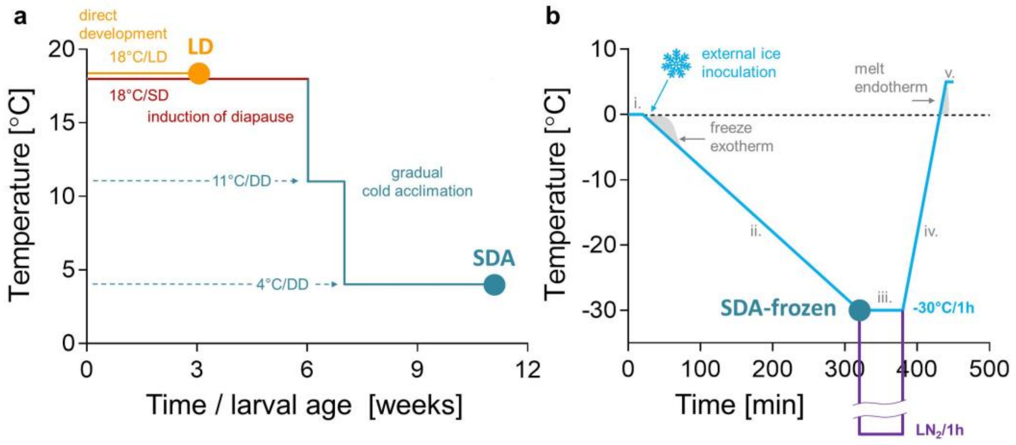


**Figure A1: Phenotypic/acclimation variants and freeze tolerance bioassay.**

**(a)** Acclimation: *Chymomyza costata* flies were reared from eggs (time 0) to 3rd larval instars (time 3 weeks) at constant 18°C under one of two photoperiods: long-day (LD, 16 h light:8 h dark) or short day (SD, 12 h light:12 h dark). The LD conditions promote direct development from larva to pupa. The LD larvae are freeze sensitive and do not survive cryopreservation in $LN_2$. The SD conditions induce entry into developmental arrest (diapause). Larvae reach their final weight aSDAer 6 weeks and are then transferred to constant darkness (DD) and low temperatures of 11°C followed by 4°C. Cold acclimation induces acquisition of freeze tolerance (phenotype SDA) including the ability to survive in $LN_2$. **(b)** Optimal protocols for freezing and $LN_2$ cryopreservation according to (Rozsypal et al., 2018). The protocol consisted of five steps set in a Ministat 240 programmable cryostat (Huber, Offenburg, Germany). The addition of a small ice crystal at the beginning of step (ii.) induces inoculative internal freezing in larvae. The SDA-frozen larvae underwent slow pre-freezing to -30°C associated with freeze dehydration of their tissues.

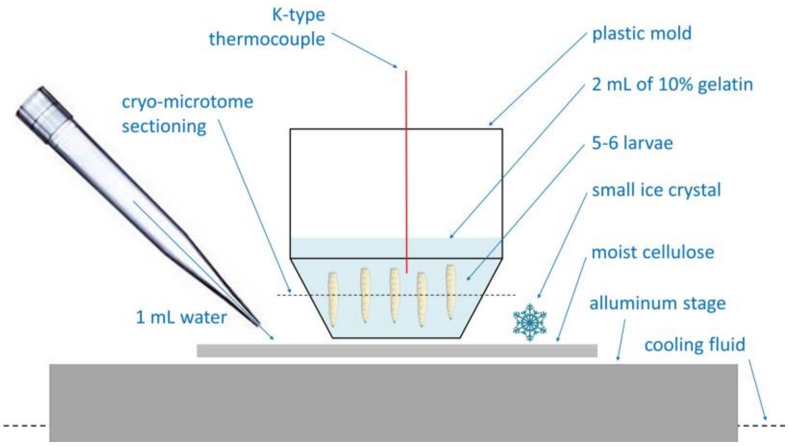


**Figure A2: Preparation of SDA-frozen larvae for MALDI-TOF/MS analysis.**

SDA larvae were embedded in gelatin in a plastic mold a top a layer of moist cellulose to which a small ice crystal was added to initiate freezing. The mold had a hole in the bottom (0.5 cm in diameter) to allow ice crystal spreading from the moist cellulose to the gelatin. The whole setup was placed on an aluminum stage pre-cooled to 0°C in the programmable thermostat F32-ME (Julabo, Seelbach, Germany) and the optimal freezing protocol (Fig. A1b) was started. As soon as the temperature inside the gelatin reached -30°C, we immersed the mold into isopentane at -140°C. Using a K-type thermocouple attached to the PicoLog TC-08 datalogger (Pico technology, St. Neots, UK), we verified that the gelatin inside plastic mold was inoculated by external ice crystals at relatively mild subzero temperatures of -1°C to -2°C. We also verified that SDA-frozen larvae survive this treatment and are able to move after re-warming.

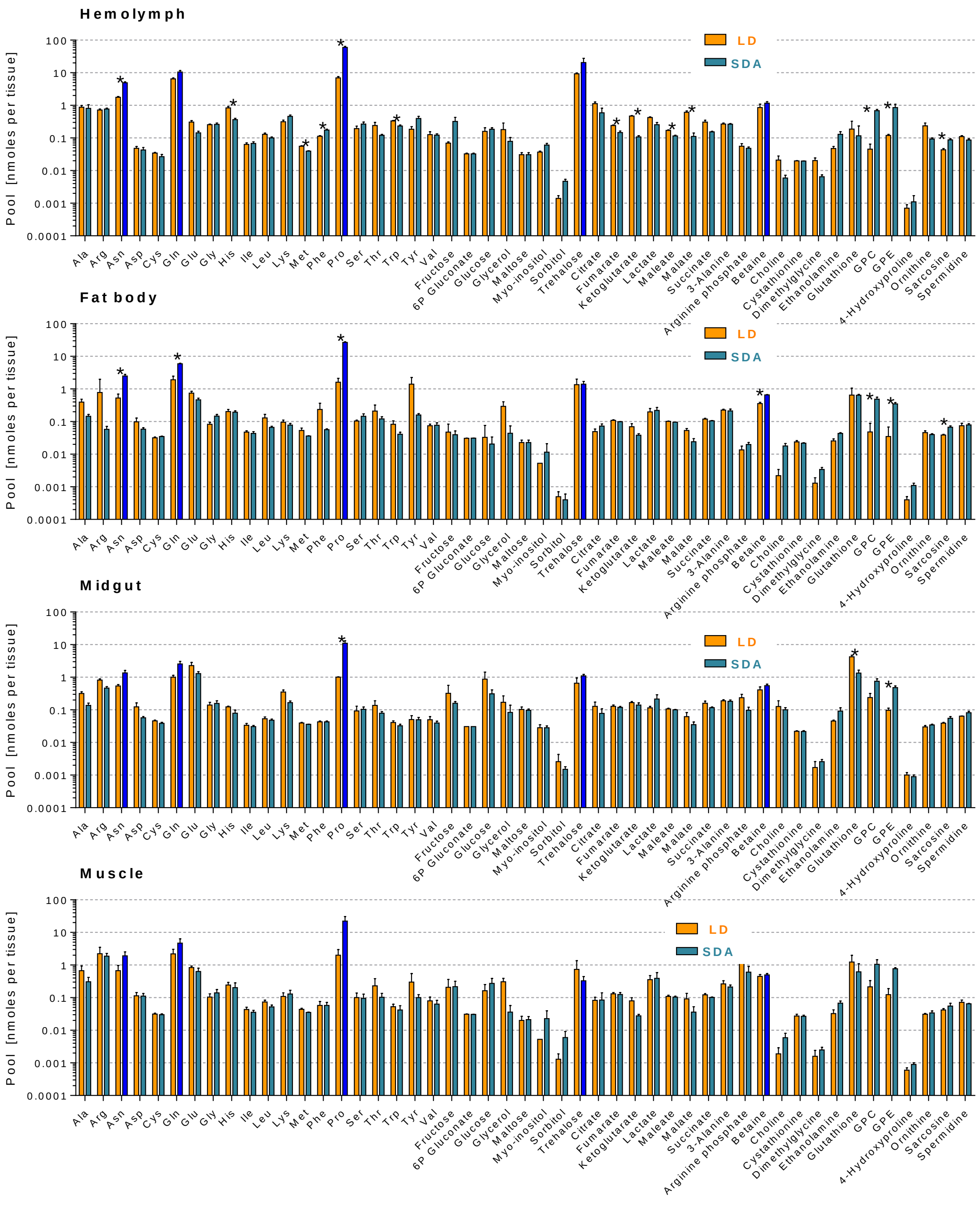


**Supplementary Figure A3: Metabolomic profiles in *C. costata* tissues.**
Forty-nine target metabolites were quantified in absolute terms using four different MS analytical platforms. Total pools in four tissues are shown for larvae of two contrasting phenotypes: LD, the non-diapause, warm-acclimated, freeze-sensitive larvae; SDA, the diapausing, cold-acclimated, extremely freeze-tolerant larvae (see Fig. A1 for further explanation). Each column is a mean ± SD ($n$ = 4 biological replicates, each containing tissues from 30 larvae). Differences between LD and SDA means were tested with unpaired t-tests corrected for multiple

comparison using the Holm-Sidak method with α = 0.01. Asterisks indicate significantly different means. Bright blue columns highlight five metabolites that are most concentrated in the SDA hemolymph.

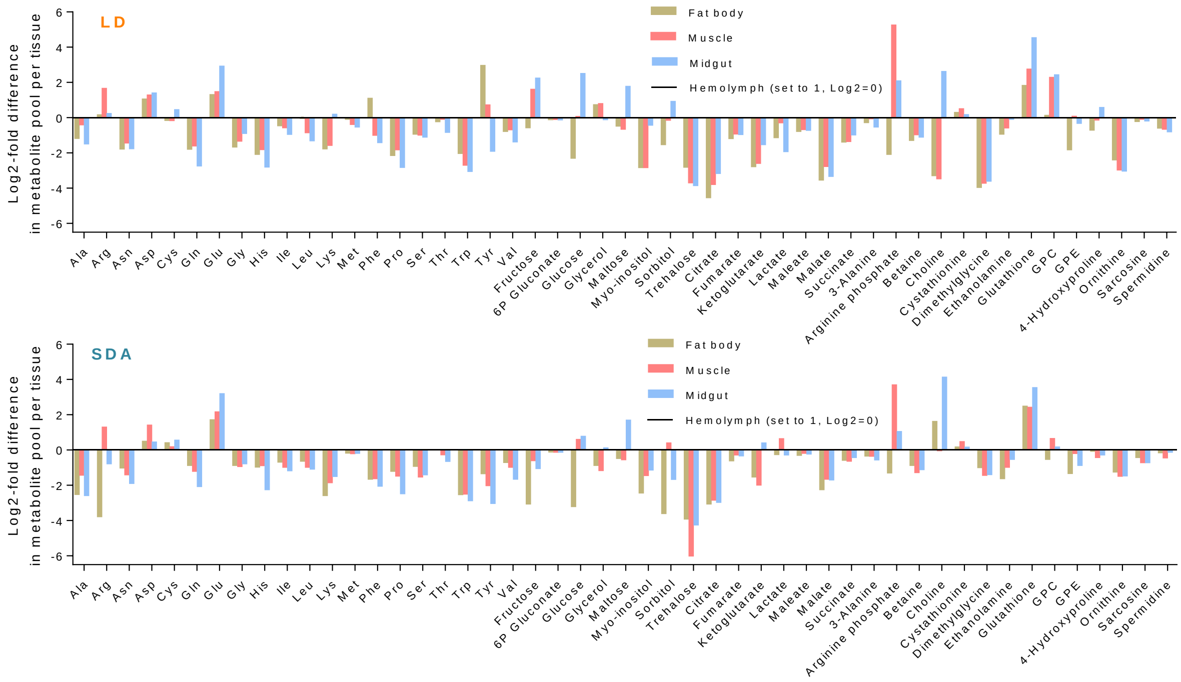


**Supplementary Figure A4: Sizes of metabolite pools in *C. costata* tissues.**
The sizes of metabolite pools (see Supplementary Fig. 1) in freeze-sensitive (LD) and freeze-tolerant (SDA) larvae were compared as $Log_2$-fold differences between tissue and hemolymph pools. The hemolymph pool was set to 1 (i.e. $Log_2$ = 0, baseline). A negative $Log_2$-fold difference means that the tissue pool is smaller than the hemolymph pool (and *vice versa*).

**Notes:**
Insect hemolymph it is known to contain relatively high concentrations of various metabolites (Mullins, 1985), including the insect 'blood sugar' trehalose (Thompson, 2003). Accordingly, the trehalose pool in SDA larval hemolymph was approximately 16 times larger than the pools in the fat body or midgut, and 64 times larger than the pool in muscle. Some metabolites predominated in the tissues rather than in the hemolymph. For example, maltose was found mainly in the midgut, reflecting the digestion of dietary starch to maltose (Applebaum, 1985); arginine and arginine phosphate were specifically enriched in muscle, reflecting the phosphagen role of arginine phosphate in insect muscle (Beis and Newsholme, 1975); aspartate and glutamate were prevalent in all tissues, reflecting their role as central sinks for ammonia groups during the metabolic conversions of amino acids (Champe and Harvey, 1994); glutathione was also prevalent in tissues due to its central role in cellular redox balance (Sies, 1999).

We assume that our MS analysis has probably captured the majority of prominent (most abundant) metabolites. We base our assumption on the following arguments: (i) the total osmolarity of the SDA hemolymph is approximately 700 $mOsmol.kg^{-1}$ (Rozsypal et al., 2018) of which 554 $mmol.L^{-1}$ is explained by our MS analysis of 49 metabolites; (ii) the metal cations of the hemolymph (dominated by $Na^{+}$ and $K^{+}$) occupy approximately 80 $mmol.L^{-1}$ (Olsson et al., 2016; Štětina et al., 2018); (iii) a similar concentration (70 $mmol.L^{-}$) must be reserved for anions (represented by $Cl^{-}$, $HCO_3^{-}$, and proteins, with 10 $mmol.L^{-1}$ excluded to account for the negatively charged metabolites aspartate, glutamate, TCA intermediates, etc.); (iv) a calculation [700 –

554 – (80 + 70) = -4] suggests that no unexplained 'osmotic gap' remains between the sum of the molar concentrations of the 49 metabolites analyzed and the hemolymph osmolality.

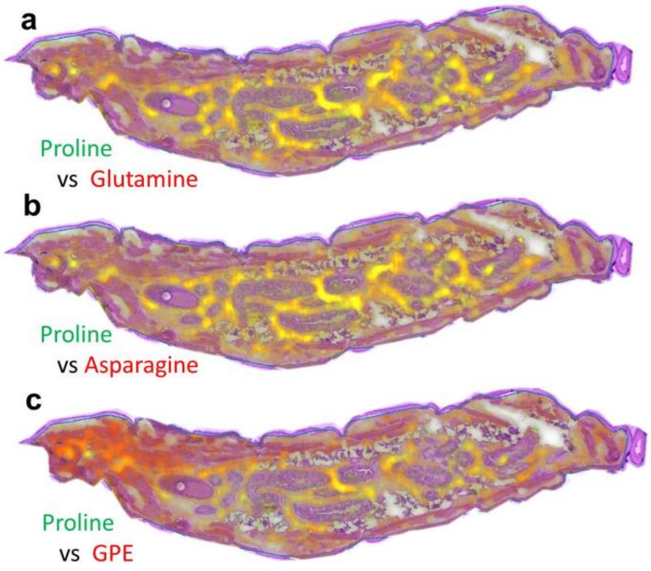


**Supplementary Figure A5: MALDI-MSI of proline colocalized with Gln, Asn, and GPE in a longitudinal section of an SDA larva.**
Colocalizations of select signals shown in color mixing code. Colocalizations were observed in hemolymph for proline (green signal) and three other metabolites: **(a)** glutamine (red signal); **(b)** asparagine (red signal); and **(c)** and GPE (red signal). Medium denoising (SCiLS tool) of signal intensities between adjacent pixels of 10x10 µm was performed.

**Note:**
The amino acids glutamine and proline are predominantly co-localized with proline in the hemolymph (the yellow colour results from the mixture of the green and red signals). The high intensity GPE signal is present in the hemolymph and in muscle tissue.

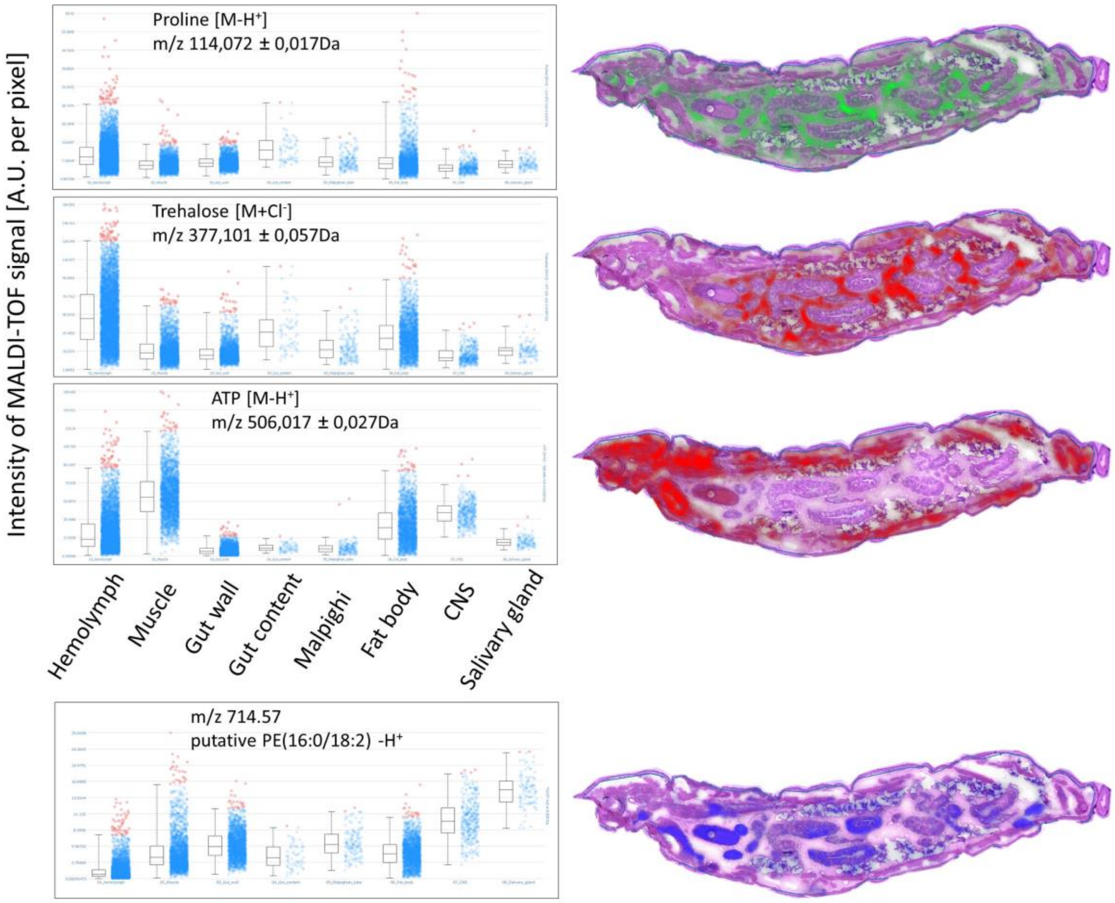


**Supplementary Figure A6: Relative quantification of MALDI-TOF signals: method and limitations.**
Examples of the tissue-specific intensities of the MALDI-TOF signals for three compounds shown in Fig. 1 (proline, trehalose, and ATP). For comparison, we show the signal of a putative phospholipid (m/z 714.57). Each coloured dot in the diagram is the signal intensity per pixel of 10x10 µm (expressed in arbitrary units, A.U.). Boxes show the median, upper, and lower quartiles; whiskers show the 0% and 99% quantiles. Orange data points are those above the 99% quantile.

**Note:**
The signal intensities per pixel vary greatly in all tissues. This is at least partly because the MALDI-MSI analysis of small molecules (smaller than m/z 700) freely dissolved in biological solutions is technically difficult (Fujimura and Miura, 2014). Diffusion of small metabolites within the tissue during matrix application and other steps of sample preparation can limit spatial resolution. In general, some analytes may delocalize from their original positions into adjacent tissues (Fujimura and Miura, 2014). An example of this problem is shown in analysis of the ATP signal, whose highest intensity is expected in tissues. Although our analysis localizes the ATP mainly in muscle, fat body, and CNS tissues, a relatively high signal intensity is also detected in hemolymph. As ATP leaks from the tissues into the hemolymph during sample preparation, the same can happen in reverse with hemolymph-specific metabolites (such as trehalose). For this reason, the MALDI-MSI is most successfully used for larger molecules such as proteins, peptides, and lipids, which are less mobile (Tuthill II et al., 2020). The analysis of the compound with m/z 714.57 (probably phosphatidylethanolamine) is such an example of a larger and less mobile molecule allowing better tissue resolution.

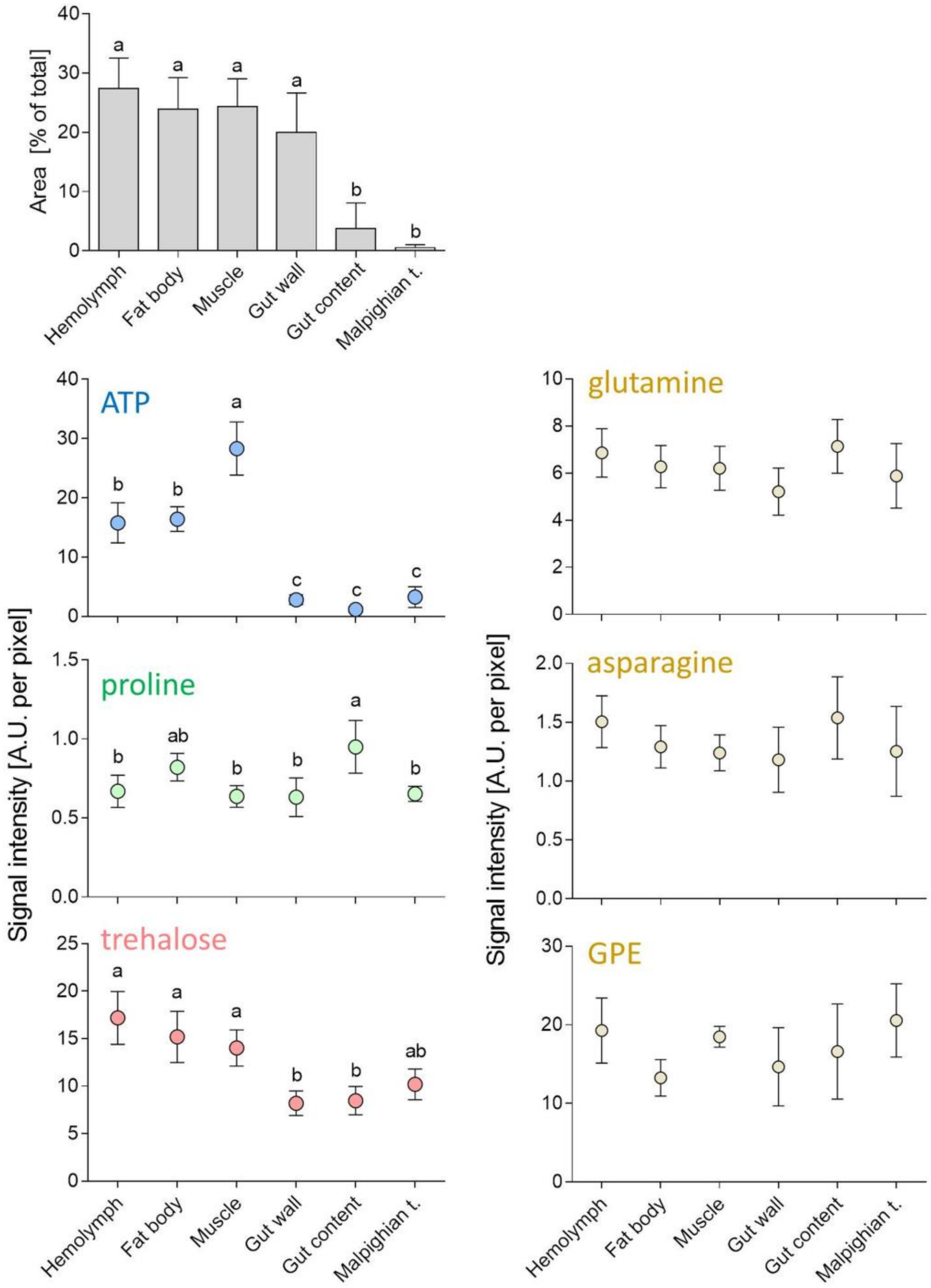


**Supplementary Figure A7: Relative quantification of MALDI-TOF signals in LD larvae.**
Results of the MALDI-TOF signal intensity analysis on transversal sections through the middle part of the larva (five larvae were sectioned). The upper leSDA figure shows the relative proportions of the different tissues on five sections (each column is a mean ± S.D. % of total area). Shown are the signal intensities of five selected putative CPs plus ATP. Each point is a mean ± S.D of five tissue-median intensities (for explanation, see Fig. A6). The relative intensities are expressed in arbitrary units, A.U. Differences between means were statistically analyzed using a one-way ANOVA followed by a Bonferroni's multiple comparison test. Means flanked by different letters (within each subpanel) are statistically different. GPE, glycerophosphoethanolamine. For details, see Appendice containing all MALDI-MSI images.

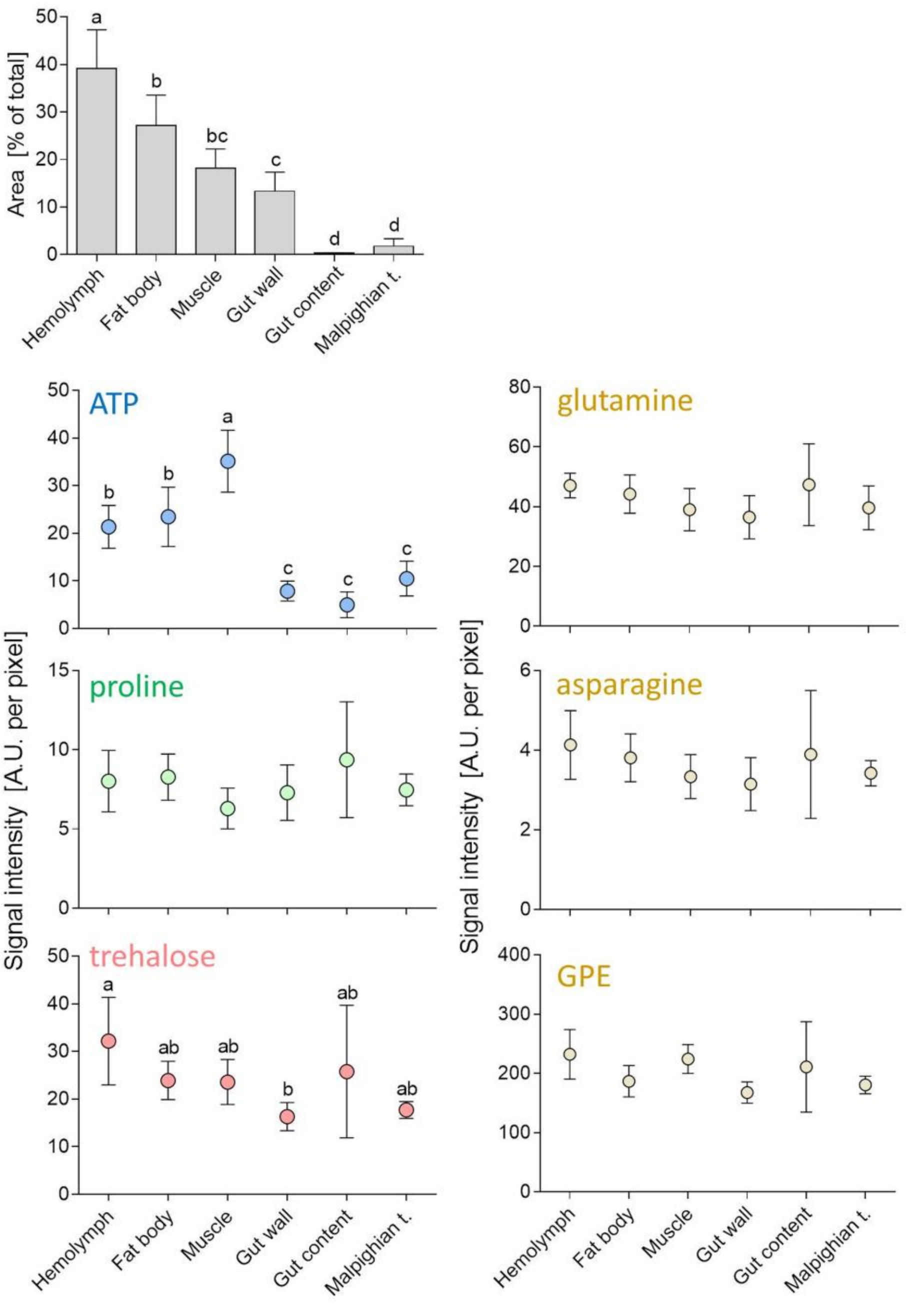


**Supplementary Figure A8: Relative quantification of MALDI-TOF signals in SDA larvae.**
All descriptions as in Fig. A7.
For details, see Appendice containing all MALDI-MSI images.

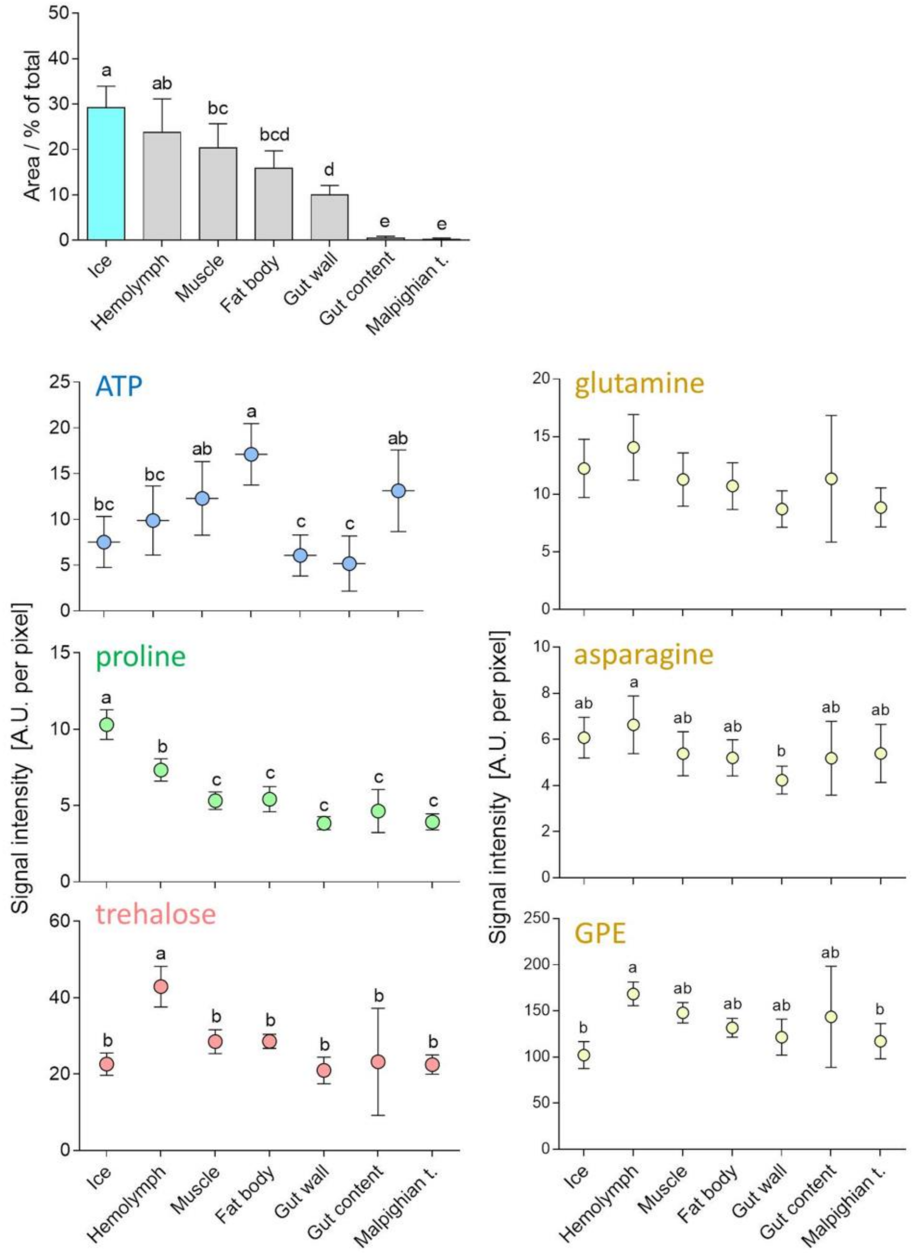


**Supplementary Figure A9: Relative quantification of MALDI-TOF signals in SDA-frozen larvae.**
All descriptions as in Supplementary Fig. A7 except that six larvae were sectioned in the SDA-frozen group. For details, see Appendice containing all MALDI-MSI images.

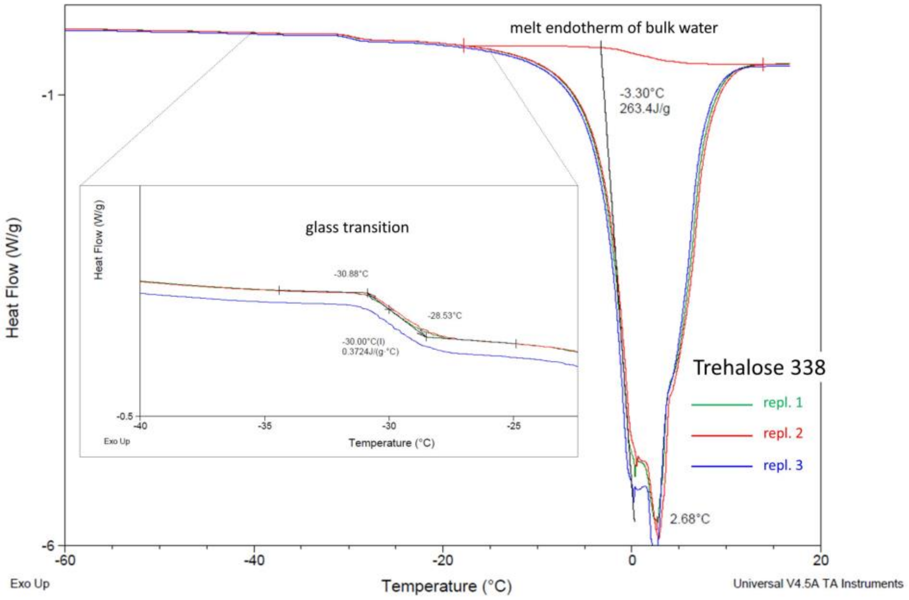


**Supplementary Figure A10: Analysis of DSC heating curves of trehalose solution (338 mmol.kg$^{-1}$) in water.**
Three technical replicates of the same solution are shown in different colours. Replicate 1 (green line) is quantified using TA Universal Analysis 2000 soSDAware as an example. The complete dataset of DSC analysis results can be found in Table A2.

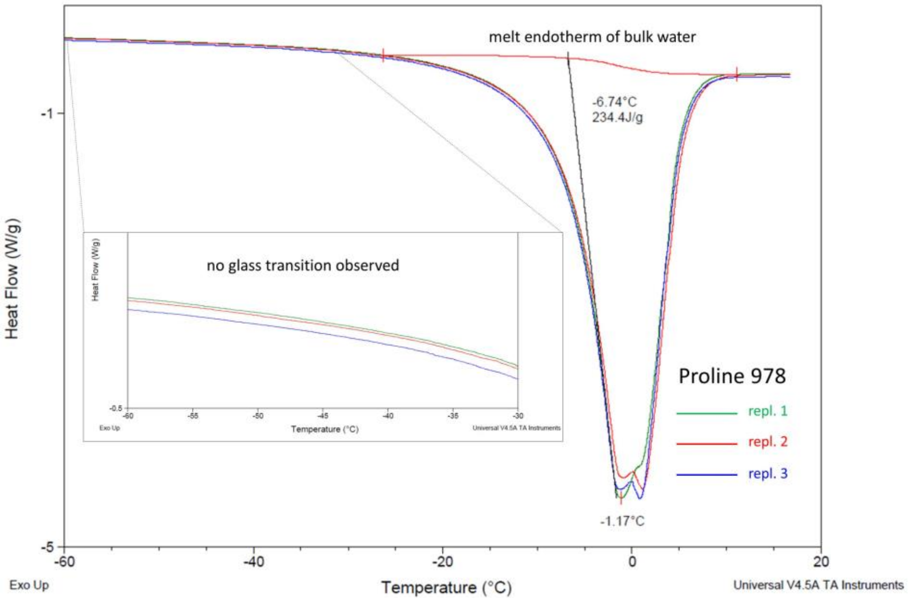


**Supplementary Figure A11: Analysis of DSC heating curves of proline solution (978 mmol.kg$^{-1}$) in water.**
Three technical replicates of the same solution are shown in different colours. Replicate 1 (green line) is quantified as an example. The complete dataset of DSC analysis results can be found in Table A2.

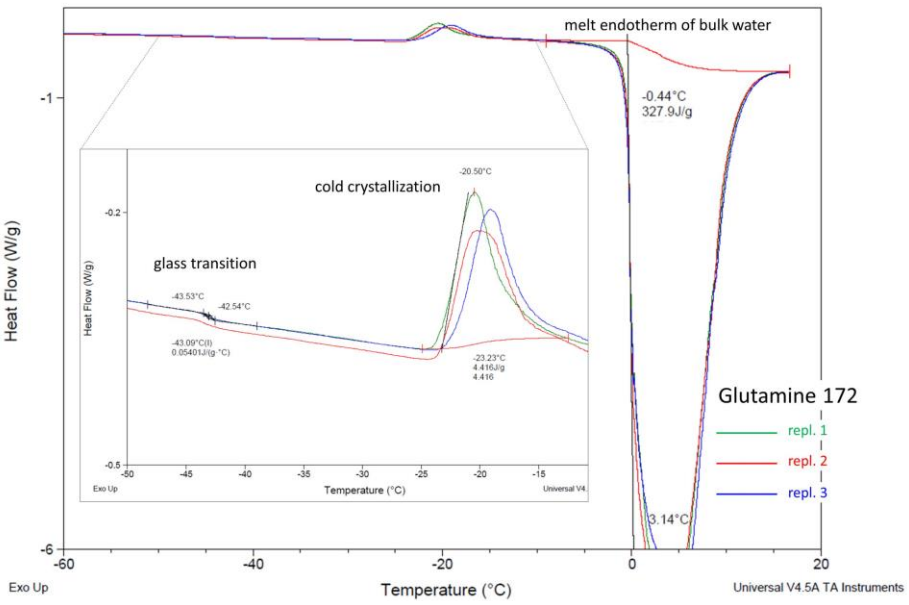


**Supplementary Figure A12: Analysis of DSC heating curves of glutamine solution (172 mmol.kg$^{-1}$) in water.** Three technical replicates of the same solution are shown in different colours. Replicate 1 (green line) is quantified as an example. The complete dataset of DSC analysis results can be found in Table A2.

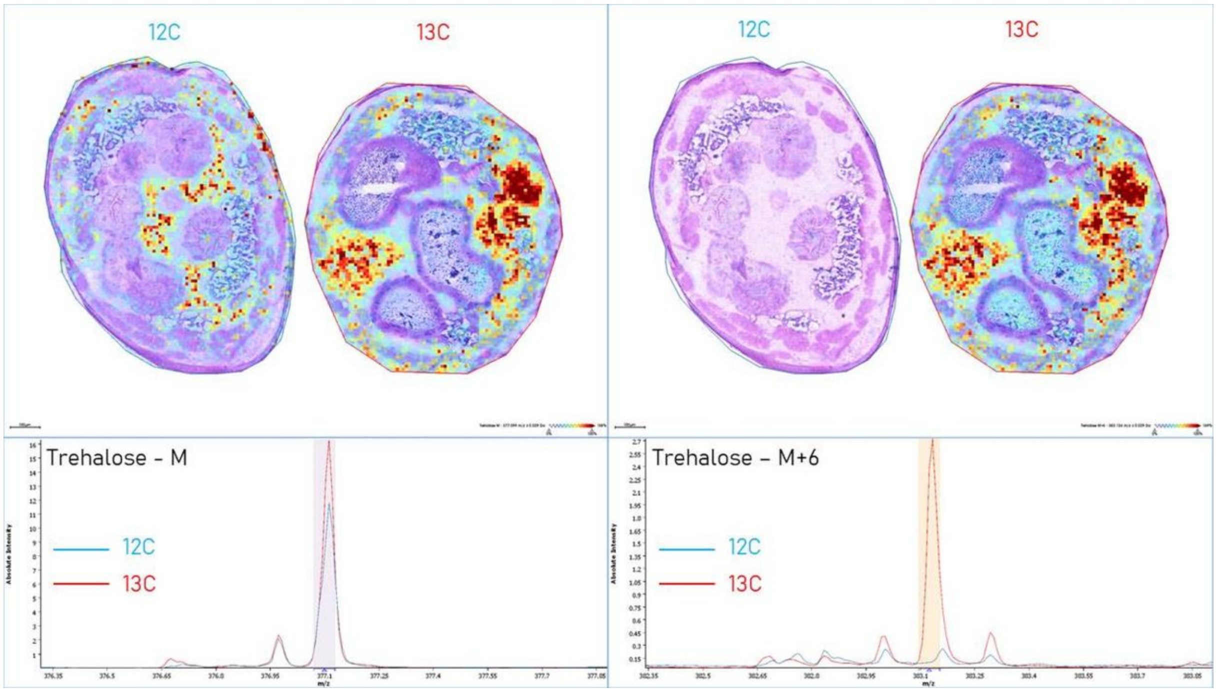


**Supplementary Figure A13: Validation of trehalose m/z peak identity using a $^{13}$C6 experiment.**
LD larvae at 17 days of age were fed a $^{13}$C6-labelled glucose enriched diet (13C, red font) for 24 hours, while the control group was fed standard diet (12C, blue font). The larvae were then processed for MALDI-MS/TOF analysis as described in Materials and Methods. The left pair of sections shows the signal of trehalose (M, m/z 337.101), the right pair of sections shows the signal of the trehalose molecules with one $^{13}$C6-labelled glucose incorporated into its structure (M+6, m/z 383.1).

**Notes:**
Larvae fed a standard 12C diet show a distinct signal from trehalose (M) in hemolymph and no signal from $^{13}$C6 labelled trehalose (M+6).

Larvae fed a $^{13}$C6 glucose-enriched diet (13C) show that the signal from M and M+6 are co-localized at exactly the same positions (mainly in hemolymph).

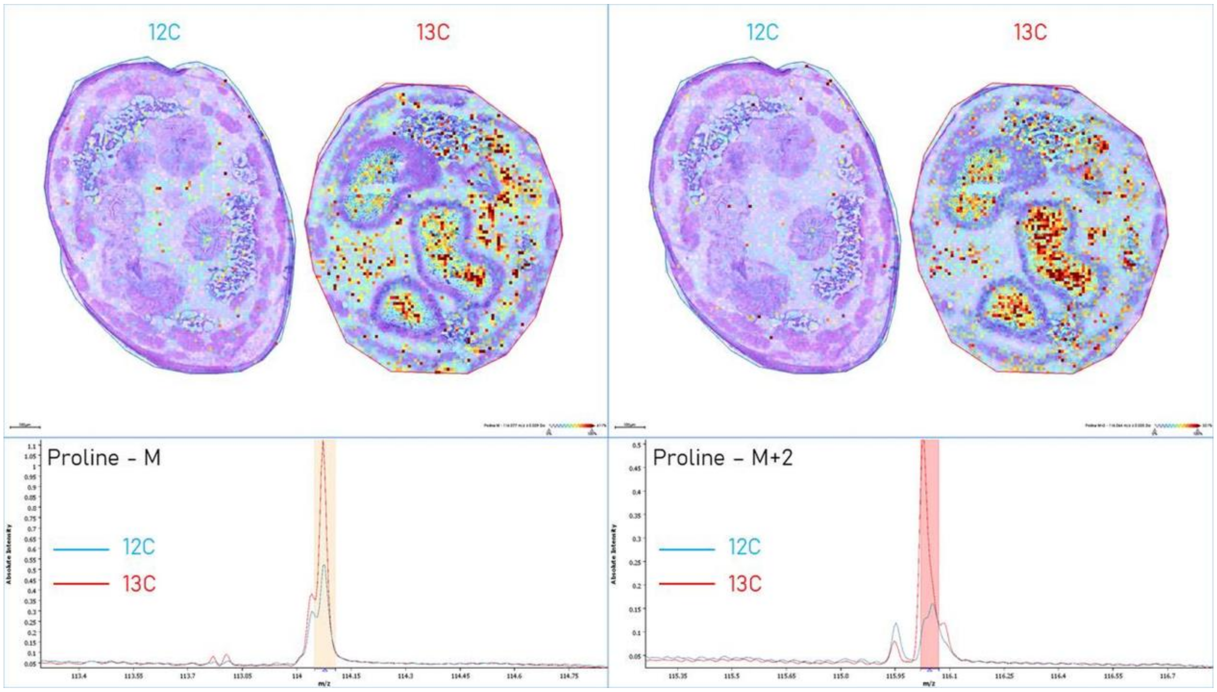


**Supplementary Figure A14: Validation of proline m/z peak identity using a $^{13}$C6 experiment.**
LD larvae at 17 days of age were fed $^{13}$C6-labelled glucose enriched diet (13C, red font) for 24 hours, while the control group was fed standard diet (12C, blue font). The larvae were then processed for MALDI-MS/TOF analysis as described in Materials and Methods. The left pair of sections shows the signal of the proline molecule (M, m/z 114.072), the right pair of sections shows the signal of the proline molecule containing two $^{13}$C-labelled atoms in its structure (M+2, m/z 116.1). Only two $^{13}$C-labelled atoms are transferred from $^{13}$C6 glucose to the proline structure as it undergoes glycolysis, TCA, and proline biosynthetic pathways.

**Notes:**
Larvae fed standard 12C diet show signal of proline (M) in the gut content, hemolymph, and fat body, while the signal of $^{13}$C6 labelled proline (M+2) is weaker or absent.

Larvae fed $^{13}$C6 glucose-enriched diet (13C) show the signal of M and M+2 co-localized at the same positions. The proline (M) signal is very abundant in the gut content of other LD larvae (Fig. A7 and Appendice), as the diet contains a relatively high concentration of proline (unpublished data).

### References used in Supplementary Figures

**Applebaum, S. W.** (1985). Biochemistry of digestion. In *Comprehensive insect physiology, biochemistry and pharmacology*, vol. 4, pp. 279-311.

**Beis, I. and Newsholme, E. A.** (1975). The contents of adenine nucleotides, phosphagens and some glycolytic intermediates in resting muscles from vertebrates and invertebrates. *Biochem J.* **152**, 23-32.

**Fujimura, Y. and Miura, D.** (2014). MALDI Mass Spectrometry Imaging for Visualizing In Situ Metabolism of Endogenous Metabolites and Dietary Phytochemicals. *Metabolites* **4**, 319-346.

**Champe, P. and Harvey, R.** (1994). Lippincott's illustrated reviews: biochemistry 2nd ed: Philadelphia: JB Lippincott.

**Mullins, D.** (1985). Chemistry and physiology of the hemolymph. In *Comprehensive insect physiology, biochemistry and pharmacology*, vol. 3, pp. 355-400.

**Olsson, T., MacMillan, H. A., Nyberg, N., Staerk, D., Malmendal, A. and Overgaard, J.** (2016). Hemolymph metabolites and osmolality are tightly linked to cold tolerance of Drosophila species: a comparative study. *J Exp Biol.* **219**, 2504-2513.

**Rozsypal, J., Moos, M., Šimek, P. and Koštál, V.** (2018). Thermal analysis of ice and glass transitions in insects that do and do not survive freezing. *J Exp Biol.* **221**, 170464.

**Sies, H.** (1999). Glutathione and its role in cellular functions. *Free Radic Biol Med.* **27**, 916-921.

**Štětina, T., Hůla, P., Moos, M., Šimek, P., Šmilauer, P. and Koštál, V.** (2018). Recovery from supercooling, freezing, and cryopreservation stress in larvae of the drosophilid fly, *Chymomyza costata*. *Sci Rep.* **8**, 4414.

**Thompson, S. N.** (2003). Trehalose—the insect 'blood'sugar. *Adv In Insect Phys.* **31**, 205-285.

**Tuthill II, B. F., Searcy, L. A., Yost, R. A. and Musselman, L. P.** (2020). Tissue-specific analysis of lipid species in Drosophila during overnutrition by UHPLC-MS/MS and MALDI-MSI. *J Lip Res* **61**, 275-290.
